# High-density sampling reveals volume growth in human tumours

**DOI:** 10.1101/2023.12.10.570995

**Authors:** Arman Angaji, Michel Owusu, Christoph Velling, Nicola Dick, Donate Weghorn, Johannes Berg

**Author notes:** Kishony lab, Faculty of Biology, Technion, Haifa 3200003, Israel (Christoph Velling).

## Abstract

In growing cell populations such tumours, mutations can serve as markers that allow tracking the past evolution from current samples. The genomic analyses of bulk samples and samples from multiple regions have shed light on the evolutionary forces acting on tumours. However, little is known empirically on the spatio-temporal dynamics of tumour evolution. Here, we leverage published data from resected hepatocellular carcinomas, each with several hundred samples taken in two and three dimensions. Using spatial metrics of evolution, we find that tumour cells grow predominantly uniformly within the tumour volume instead of at the surface. We determine how mutations and cells are dispersed throughout the tumour and how cell death contributes to the overall tumour growth. Our methods shed light on the early evolution of tumours in vivo and can be applied to high-resolution data in the emerging field of spatial biology.

---

The evolution of a solid tumour is governed by the division, motion, and death of cancer cells. Genetic mutations arising during cell divisions can serve as cell markers to track this dynamics. From the observed spatial distribution of mutations, it should in principle be possible to infer the spatio-temporal principles of tumour evolution: Is the growth rate uniform across the tumour, or does growth predominantly take place near the edge of the tumour? What is the interplay between the tissue dynamics of the tumour and its genetic evolution? These broad modes of tumour evolution affect for instance the signature of neutral evolution, the response to selection, or the number of low-frequency mutants which can confer therapy resistance [1, 2].

However, to answer such questions on the basis of genetic tumour data is challenging because only partial information is available: (i) The sequencing depth (average number of reads covering a nucleotide in NGS sequencing) is finite. This means that only high-frequency mutations are observed. (ii) Usually only a small number of samples are taken from different parts of a solid tumour, which limits the information on mutations present in the full tumour [3, 4]. (iii) Longitudinal data from ctDNA measurements are highly limited in the observable range of mutations and provide noisy frequency estimates.

Over the next years, some of these restrictions will be lifted by the advent of spatial genomics [5, 6, 7]. These techniques allow assaying the genomic information almost at single-cell level in intact tissue sections. Currently, the attainable sequencing depth is too low to identify point mutations across different parts of the tumour. However, it is clear that the coming-of-age of spatial genomics will bring new opportunities to understand the past evolution of a population of tumour cells from a late-stage snapshot. This implies a need for new tools to analyse spatio-temporal evolution, since standard tools of population genetics, like the site-frequency spectrum, are designed for spatially mixed populations and disregard spatial information.

One particular question concerns two different modes in which a tumour can grow; surface growth and volume growth. Under surface growth, the cancer cells divide predominantly at the border with healthy tissue. The potential reasons for this spatial dependence include higher nutrient levels near normal tissue, higher levels of metabolic waste products in the tumour bulk, or mechanical stress in the tumour centre [8, 9]. A faster growth rate at the edge of the tumour leads to a radially outward growth of the cell population. The surface growth mode is well-known from bacterial growth [10]. In tumours, some evidence for surface growth comes from histological stainings, which show an enhanced level of the Ki-67 protein (a cellular marker for proliferation) near a tumour surface [11, 1, 12]. However, the reverse situation has been found as well [6], with elevated Ki-67 levels near the centre of a renal carcinoma. The surface growth mode also has a long history in the modelling of tumour evolution [13, 14, 15, 16, 17, 18, 1, 19, 20, 21, 22] and has been used to analyze multi-region tumour sequencing data [19, 21, 2, 23, 24, 25, 26].

In volume growth, on the other hand, cancer cells grow irrespective of their location in the tumour: although each cell has a physical location, and upon division its offspring is in a similar location, location does not affect cell division or death. Under volume growth, subclones can originate from any location in the tumour [6]. As a result, under volume growth mutation frequencies evolve exactly in the same way they would do in a well-mixed population, even though the resulting tumour will generally be spatially heterogeneous (see below). Mixed population models have been used extensively to model all aspects of tumor evolution, from tumorigenesis to the formation of metastases and the response to therapy [27, 28, 29, 30, 31, 32]. They also form the basis of almost all population-genetic approaches to analyzing tumour data [33, 34, 35].

To tell between these two different evolutionary dynamics, we use high-resolution data on the spatial distribution of mutations found in solid tumours. We use whole-exome mutation data obtained from hepatocellular carcinomas and published previously [36, 24]. From these snapshots of late-stage tumours we infer how the tumours grew in earlier stages. To this end, we develop metrics of intra-tumour heterogeneity which leverage the information from the spatial position of all samples.

## Methods

### Spatially resolved data

We analyze two datasets where large numbers of samples (*>* 100) were taken from different hepatocellular tumours, once from a two-dimensional section [36], and once from a three-dimensional microsampling [24].

In Ling *et al*. [36], 285 samples of a planar section of a hepatocellular carcinoma of diameter 35 mm were taken and analyzed. Each sample consisted of about 20000 cells [36] determined by cell counting. 23 of these samples were subjected to whole-exome sequencing (average read depth of 74 reads per nucleotide), and loci with mutations found in these 23 samples were probed by genotyping (by Sequenom and Sanger sequencing) in all 285 samples [36]. Figure 1A shows the positions of the samples and the mutations found at each position. The data thus effectively consist of a set of 285 samples at high spatial resolution, but with limited genomic information (genotyping), and a subset of 23 samples at high genomic resolution from whole-exome genomic sequencing, but limited spatial resolution. We reanalyzed the whole-exome data as described in SI S4.2. The genotyped data consists of 35 mutated loci. Calling variants jointly on all 23 whole-exome samples and filtering as described in SI S4.2 returned 217 mutated loci.

**Figure 1:**
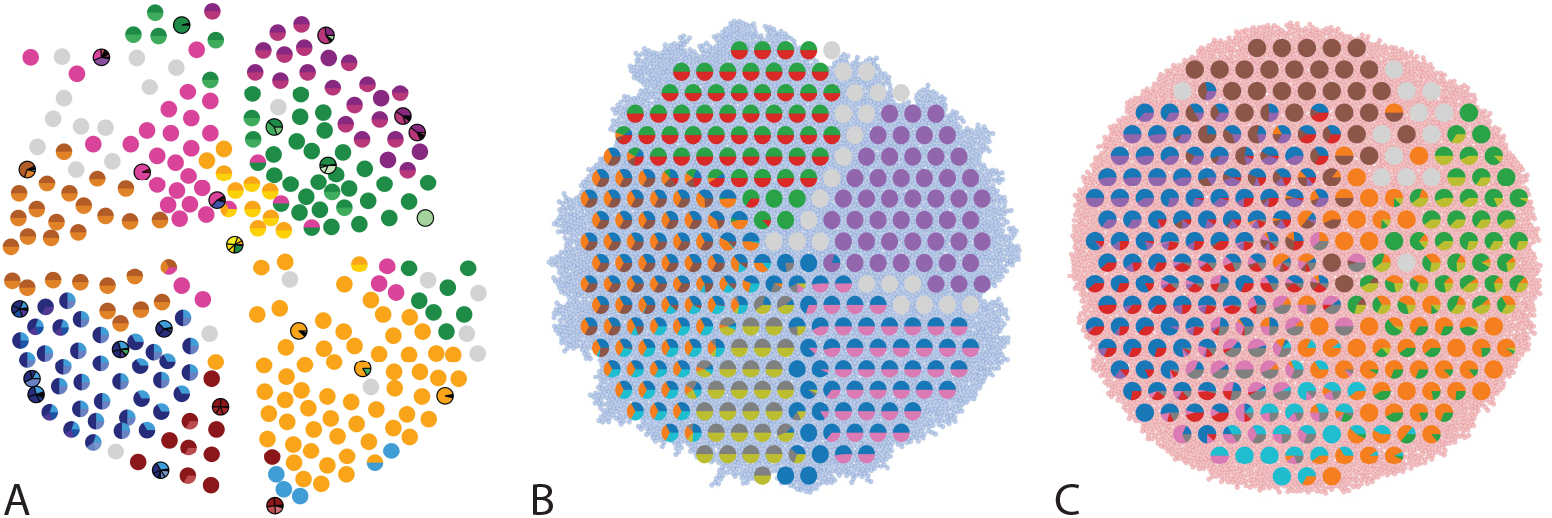
Multi-region sampling of a hepatocellular tumour and cell-based simulations. A shows the spatially resolved sequencing data of 285 samples of a hepatocellular carcinoma analyzed by Ling *et al*. [36]. Each sample is indicated by a small pie chart in which colors indicate specific mutations, and slice sizes indicate the mutation frequencies within each sample. The 23 samples highlighted by a black outline were also subjected to whole-exome sequencing. The samples form a honey-comb structure, because the tumour slice had been cut into four quadrants, see Fig S1 in [36]. B and C Results of a cell-based simulation in the surface growth mode (B) and the volume growth mode (C). In each case, 280 evenly spaced samples were taken from the population of 10000 cells of a 2d simulation, see text. The most frequent mutations are shown as in (A), superimposed on the structure of the simulated tumour.

In the second data set, Li *et al*. [24], samples were taken from multiple planar sections of two different hepatocellular tumours from the same patient. This gives a three-dimensional picture of the genetic heterogeneity of these tumours. 169 samples were taken from 11 sections of tumour T1, which was approximately of 20 mm diameter, and 160 samples from 6 sections of tumour T2, which was of approximately of 15 mm diameter. Each sample consisted of ca. 3200 cells. 16 (9) samples from T1 (T2) were whole-genome sequenced and the remainder genotyped using the Ampliseq method. Li et al. report 906 (564) mutations used for genotyping of T1 (T2), of which 563 (259) pass our filters.

### Computational model of volume and surface growth

We use an off-lattice cell-based model, well-known from simulations of tissue mechanics [37, 38, 39, 40], which we endow with mutations entering the population at a constant rate per cell division.

In this model, each tumour cell is described by a sphere centered on a particular point in 3d space or 2d space, as well as a genome. Each cell division introduces a new point adjacent to its parent in a random (uniformly distributed) direction, and each cell death removes a point. At division both mother and daughter cell can acquire mutations, the number of mutations is Poisson distributed with mean *µ*. The spherical are not allowed to overlap and can push one another out of the way to achieve this. This off-lattice approach differs from cellular automaton models of tumour growth, where cells occupy discrete lattice sites [1, 2]. Specifically, in off-lattice models like the one we use, cells can gradually push one another out of the way during growth, see below. Cells cannot move by themselves but as they divide overlaps are introduced and resolved by pushing the neighboring cell.

We implement the resolution of overlaps introduced by a cell division using a ‘width-first’ pushing-algorithm, where the immediate neighborhood of the newly placed cell is resolved first before proceeding to the new overlaps this step introduces. A fast search first identifies overlaps with the current cell in its close neighborhood. The cell then pushes its neighbor along their connecting axis only by the overlap plus a small margin which acts as a buffer and avoids many tiny recurring overlaps. The neighboring cell is then added to a queue for subsequent iterations. Neighborhoods are shuffled such that there is no particular order in which cells are pushed. After each overlapping cell in the neighborhood is pushed and added to the queue, the search for overlaps is repeated on the first cell in the queue, adding new pairs to the queue, which are in turn iteratively resolved until the queue is empty. Thereby each cell division can set off a cascade of rearrangements of cell positions. This level of microscopic detail of course comes at a computational cost. Using off-lattice cell-based modelling, population sizes of several ten thousand cells can be simulated, compared to billions of cells with a cellular automaton [1].

In regions of high cancer cell density, the division rate may be reduced (for instance due to lack of nutrients, toxic metabolic products, or mechanical stress). We effectively encode such potential spatial effects in the rate at which individual cells divide: the cell division rate *b* = *b*(*ρ*) depends on the local density of cells *ρ* = *ρ*(**r**). For simplicity, we consider a division rate *b*(*ρ*) which decreases in a straight line from *b*(*ρ* = 0) = *b*_0_ to *b*(*ρ* = *ρ*_*c*_) = 0, see Fig. S1. Setting *ρ*_*c*_ = ∞ leads to a constant division rate for all cells, irrespective of their local density. This growth mode is termed volume growth. Lower values of *ρ*_*c*_ lead to an increased growth rate in parts of the tumour where the local density of cancer cell is low, and hence enhance the growth rate at the surface of the tumour relative to the bulk of the tumour. Surface growth is characterized by a *ρ*_*c*_ equal to the density defined by the minimal cell distance. By changing *ρ*_*c*_ one can thus tune the dynamics of this model continuously from volume to surface growth. To describe the genetic changes in the tumour, we use an infinite-sites model and a constant mutation rate per cell division. Figure 1B and C show the resulting tumor sections for the surface and volume growth mode, respectively.

For a detailed description of the model, its implementation and performance in comparison with variants of the kinetic Monte Carlo and pushing algorithms, see SI S1. The code is available as a Julia package at https://github.com/aangaji/TumorGrowth.

## Results

### Volume versus surface growth

Under surface growth, cells divide predominantly near the edge of a tumour. Under volume growth on the other hand, cells divide uniformly across the entire tumour mass. To distinguish these two modes in empirical data, we look at the angle between a line from the tumour centre to a parent clone and from the parent to their offspring, see Fig. 2A and SI S2. Under surface growth, offspring tend to lie radially outward from their parents, leading to a distribution of these direction angles centered around zero. Under volume growth, cells divide isotropically, leading to a uniform distribution of angles. The distribution of direction angles is a model-independent metric, it does not rely on a particular model of cell dynamics, population dynamics or sampling. It is also robust against the presence of selected subclones: if a subclone grows isotropically from some point, but at an elevated rate, it will contribute to a flat distribution of angles like all other clones.

In Fig. 2B we show the distribution of direction angles found in the spatially resolved data of Ling *et al*. [36]. The empirical data clearly show a uniform distribution of the direction angle, and thus no evidence for radially outward growth caused by faster growth near the tumour’s edge. For comparison, we look at the distribution of direction angles in simulations of different growth modes. For surface growth, the histogram of direction angles shows a pronounced maximum at *θ* = 0 produced by the radial outgrowth of clones (Fig. 2C), in a simulation of volume growth, we find a flat distribution (Fig. 2D). These simulations were done with samples placed in the same positions as in the real tumour to eliminate effects from uneven sampling.

**Figure 2:**
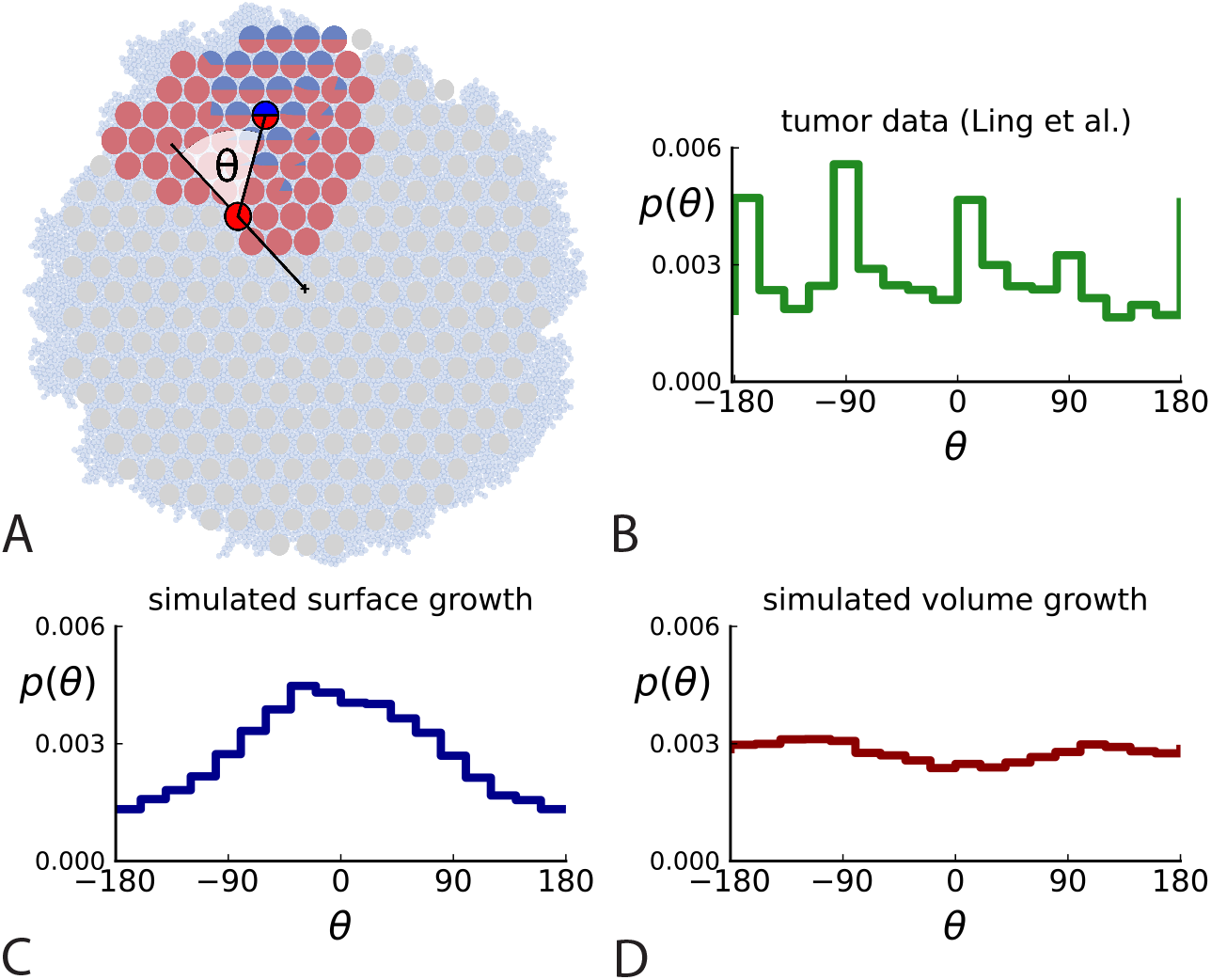
Relative position of mutants under different growth modes. A The direction angle *θ* quantifies the direction of a new mutant clone relative to its parent clone. In this illustration, a new mutant clone indicated in blue appears and grows radially outward on a red parental background, resulting in an angle *θ* near zero. Each pair of cells indicated in red and blue contributes to the distribution of *θ*, with the statistical weight of each mutant clone adding to one (see SI S2). B The distribution of angles *θ* for different mutant clones found in the spatially-resolved data of Ling *et al*. [36]. The peaks at *θ* ≈− 180^deg^, − 90^deg^, 0^deg^ come from individual clones. The contributions from different clones to the distribution of angles is shown in Fig. SI S2. C and D show the corresponding distribution of angles for numerical simulations. Subfigure C shows simulations of surface growth, resulting in a distribution of direction angles with a pronounced maximum near zero. Under volume growth (D) a nearly flat distribution is seen. For C and D, simulations were run in three dimensions with a maximum population size of 40000 cells grown at division rate *b* = 1, a rate of cell death *d* = 0.4 and *d* = 0.8 for surface and volume growth respectively, and a whole-exome mutation rate *µ* = 0.3 before taking a two-dimensional cross-section of 280 samples mimicking the sampling procedure in [36]. (The different death rates were chosen to make the extinction probabilities of the populations comparable for the two cases. Changing these rates did not affect the distributions of angles.)

This result is corroborated by the average number of mutations of a clone as a function of its distance from the tumour centre. Under surface growth and neutral evolution, the number of cell divisions since tumorigenesis, and hence the number of mutations found in a cell, increases with distance from the tumour centre. However, we do not find this signature of surface growth in the tumour data. Instead, we find that the number of mutations at different distances from the centre is compatible with volume growth (see Fig. S11).

Similarly, the dataset of Li *et al*. [24] with three-dimensional sampling also shows the signature of volume growth, both in the distribution of direction angles (SI S2.2) and in the number of mutations as a function of distance from the tumour centre, see SI S3.3. This result is at variance with a recent spatio-phylogenetic analysis performed by Lewinsohn *et al*. [41], which finds a signal of surface growth on the same data. In SI S3.4, we show that the findings of [41] are compatible with volume growth; on artificially generated volume-growth data sampled like in Li *et al*., the algorithm used in [41] also returns a spurious signal of surface growth.

### Spatial dispersion of cells

Another aspect where surface and volume growth lead to qualitatively different behaviour is the spatial dispersion of mutations. Cells carrying a particular mutation can in principle form a tightly spaced colony or be dispersed throughout the tumour mass, see Fig. 3A. It turns out this dispersion of mutants differs between surface and volume growth.

**Figure 3:**
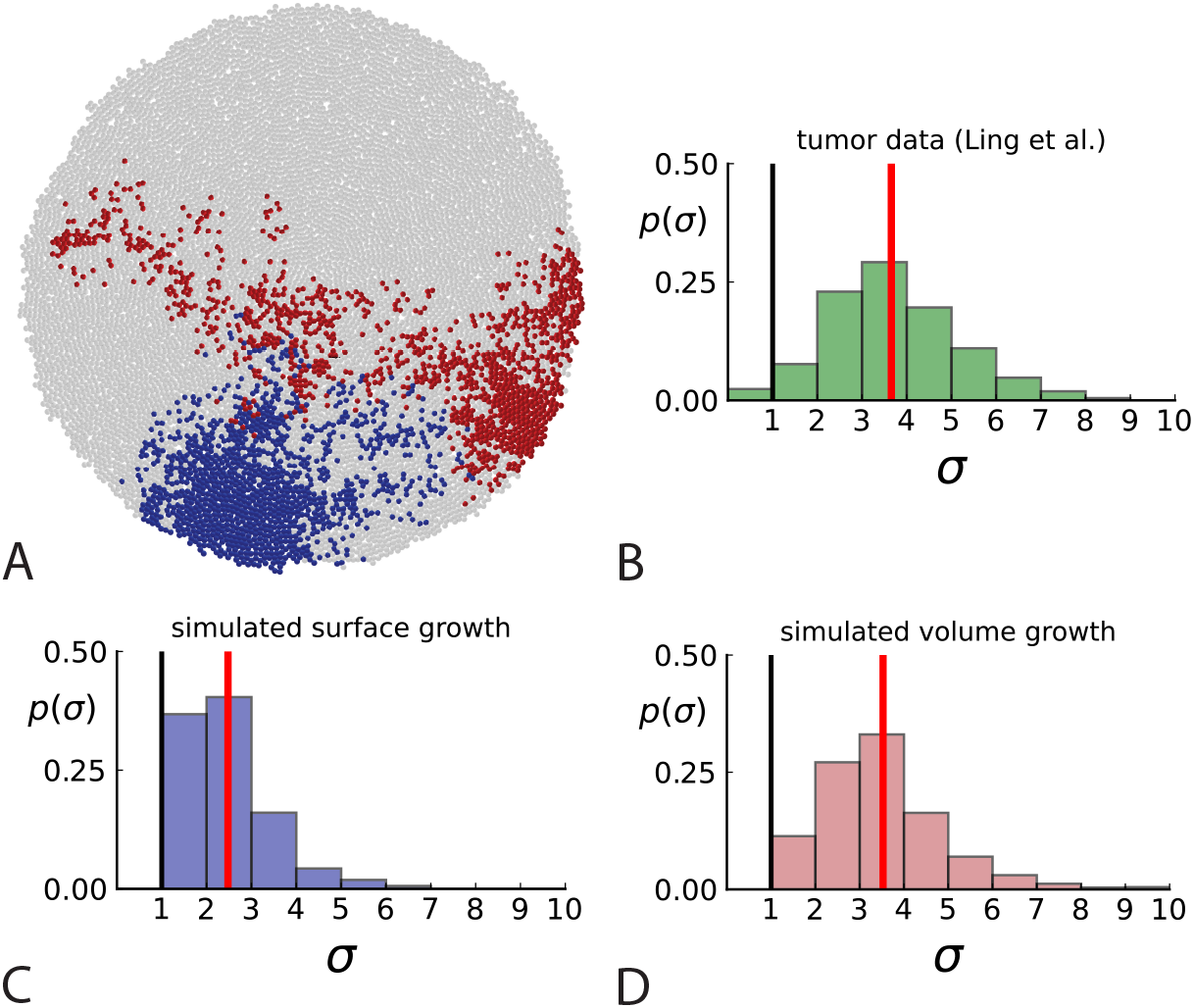
Dispersion of mutations within the tumour. (A) Cells with a particular mutation can form tight spatial clusters within a tumour (simulated example here: mutant shown in blue), or they can be more widely dispersed (mutant shown in red). We quantify the dispersion of a mutation using the dispersion parameter *σ*, see text and SI S4. In this illustrative example, the blue mutation has a small dispersion parameter *σ* = 1.3, the red one has *σ* = 2.5. (B) Histogram of the dispersion parameters *σ* across 217 mutations in the whole-exome data of [36]. The red line indicates the mean of the histogram. (C) and (D) show the corresponding histograms for simulations of surface growth and volume growth, respectively. The simulations were run in 3D with populations grown up to 40000 cells before taking 23 evenly spaced samples from a 2D cross-section. Only mutations with a whole-tumour frequency larger than 1*/*40 were considered, mimicking the limited sequencing resolution in the Ling *et al*. data. Simulation parameters were division rate *b* = 1, mutation rate *µ* = 0.3, and death rates *d* = 0.4 and *d* = 0.8 for surface growth and bulk growth, respectively.

To quantify the spatial dispersion of a mutation, we define the dispersion parameter *σ*: For a given mutation, we compute the average distance of all pairs of samples carrying that mutation and divide by the average distance these cells would have if they were arranged next to one another in a spherical shape (the tightest configuration possible). Values of *σ* much larger than one can arise when a mutation is scattered throughout the tumour, with many cells that do not carry that mutation located between cells that do. The dispersion *σ* can be estimated from mutation frequencies in the whole-exome sequencing data of [36], see SI S4 for details.

Fig. 3B shows a histogram of the dispersion parameter *σ* for 217 mutations in the whole-exome sequencing data of [36]. We find an average value of the dispersion of approximately 3.5, but the values for specific mutations can be much higher than that (up to 9). Fig. 3C and D show the corresponding distributions for simulations of volume and surface growth respectively. The distribution of dispersion parameters found under volume growth agrees with that in the empirical data (Kolmogorov-Smirnov statistics 1.13), whereas surface growth generates lower values of the dispersion parameter (Kolmogorov-Smirnov statistics 5.74 indicating a poorer match with the empirical data, see SI Section 4.3 for details). Also in the three-dimensional dataset of Li *et al*. [24] we find higher values of the dispersion parameter than expected under surface growth (SI S4.4). However, the larger distances between samples in 3 dimensions, compared to the samples in a single plane, turn out to limit the accuracy with which the dispersion parameters can be computed. A higher number of samples would be needed to make the distances between samples comparable between the two- and three-dimensionally sampled data.

Volume growth offers a simple mechanism causing the dispersion of mutations: Tumour cells displace one another when new cells are born and grow. In this way, two cells which are initially close to one another and share a mutation can be pushed apart by the birth and growth of adjacent cells that do not share the mutation. The further apart a pair of cells becomes, the higher the probability that they will be moved even further apart due to the motion of the increasing number of cells between them. This amplification of initially small distances leads to an instability that can generate the large values of the dispersion *σ* found both in the data of [36] and in numerical simulations of the volume growth model (Fig. 3B and D, respectively). On the other hand, under surface growth, cells do not push each other apart, leading to lower values of the dispersion parameter. Also, high values of the dispersion parameter cannot arise spuriously due to positive selection, as selected clones would enter the population later than neutral ones (given the same final frequency), and thus do not have as much time to be moved within the tumour.

The dispersion of cells in a growing population via this pushing effect is well known in tissue mechanics [42, 39]. In the context of cancer, it provides a mechanism for the clonal mixing found by ultra-deep sequencing of multiple samples from a solid tumour [19, 43, 21], which leads to mutations that occur at high frequencies in some samples being present at low (but non-zero) frequencies in other samples. Dispersion and clonal mixing thus arise as a by-product of volume growth. This is in contrast to surface growth, where a specific dispersal mechanism has been postulated to account for the observed spatial distribution of mutations [1].

To probe how this migration of cells between different regions of a tumour affects multi-region sampling, we look at the number of mutations detectable in a limited number of samples. Specifically, we consider the fraction of mutations present in a certain subset of samples (relative to the mutations appearing in all the samples taken together) and ask how this fraction is affected by migration. Fig. 4A shows the fraction of mutations detectable in a given number of samples randomly picked from the 23 whole-exome samples of [36]. One finds that typically about 75% of all the mutations can be detected with only 5 samples. We repeat this analysis but do not count mutations in a sample that occur with a higher frequency in some other sample. The rationale is that these mutations have arisen elsewhere (where they are present at higher frequency) and have migrated into the sample, where they are now present at a lower frequency. With such mutations removed, 5 samples typically only contain about one fifth of the mutations. This is in line with 5 samples representing only about one fifth of the 23 whole-exome samples. Migration of cells thus makes a single sample capture a far larger share of the genetic variability than expected from its share of the sampled tumour volume and provides an a-posteriori rationale for assessing the genetic variation found in a tumour on the basis of only a few samples.

**Figure 4:**
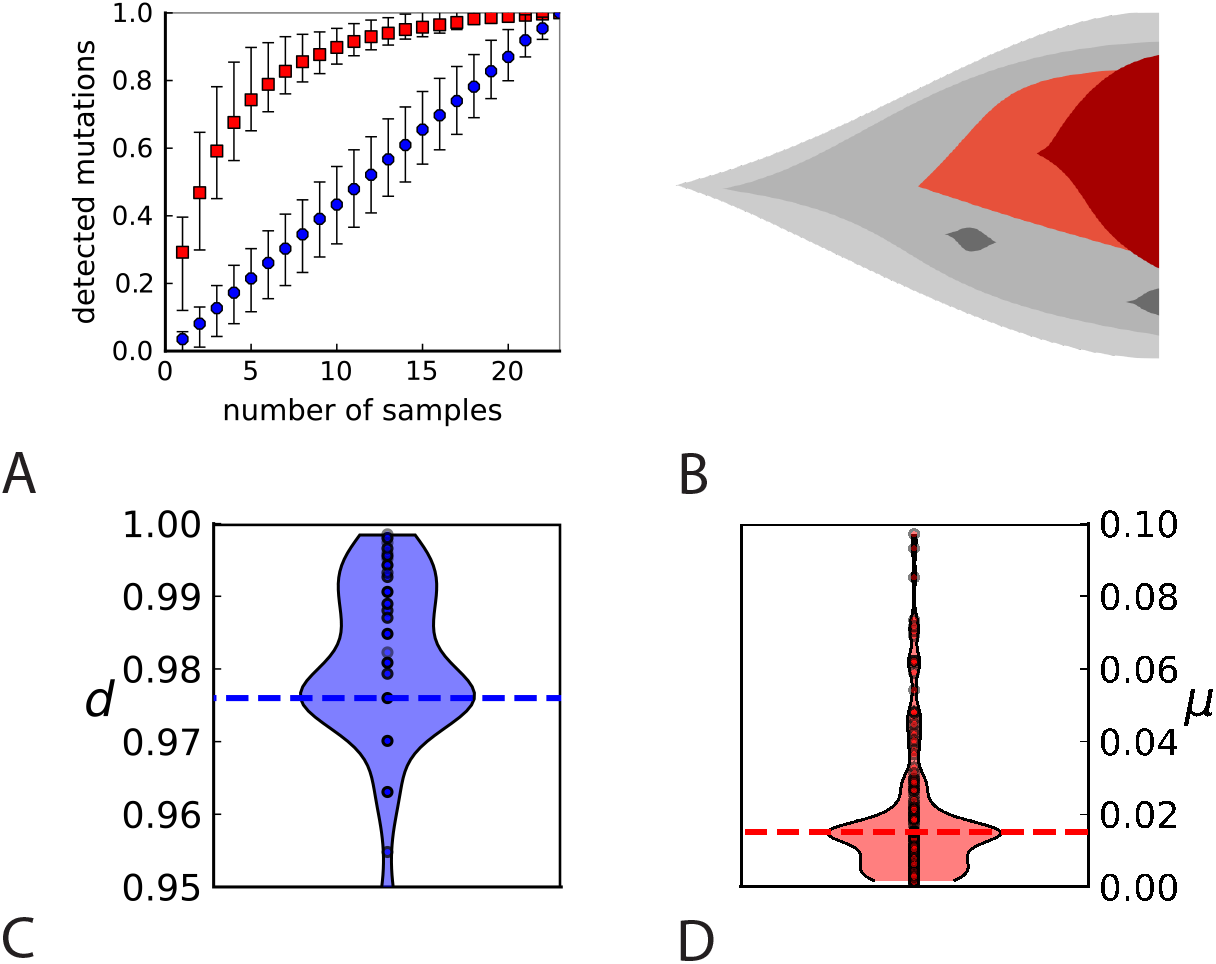
Rate of cell death and the mutation rate. (A) We ask how a limited number of samples identify mutations. We pick a subset of the whole-exome sequenced samples of Ling *et al*. [36] and plot the fraction of mutations present at least in one of these samples against the number of samples in the subset (red symbols, fractions are relative to the number of mutations present in at least one of the 23 samples. Mutations must be supported by at least 5 reads at a coverage of at least 150). The procedure is repeated (blue symbols) with those mutations removed that occur in some other sample with a higher frequency than the frequencies with which the mutation occurs in the subset of samples. Error bars indicate the range of the 95-percentile. (B) The schematic Muller plot shows how cell death leads to the loss of clones, some of which have extant offspring. Time runs on the horizontal axis, the vertical shows a number of different clones indicated by different colours. In the example shown here, the clone shown in light red becomes extinct, leaving behind its darker-shaded offspring clone with no parental clone. The rate of this loss of parental clones depends on the rate of cell death, and can be used for inference, see text. (C) shows the inferred rate of cell death and (D) the inferred rate of mutation per generation. The violin plots show how the inferred values vary when subsampling different fractions of all mutations and scaling the inferred mutation rate correspondingly, with the dashed lines indicating the mean inferred values. (The fraction of mutations sampled ranges from 0.5 to 0.9, with the results shown separately in SI 9.)

### Cell turnover

At low rates of cell death, tumours grow nearly at the rate at which tumour cells divide. Higher rates of cell death slow down the net growth of the population and lead to a constant turnover of cells. The relative rates of cell birth and death have been studied in cell cultures since the classic work of Steel [44] using thymidine labelling. Also in cell cultures, live cell imaging has been used to track individual cell divisions [45]. In the following, we use the genetic record in spatially resolved genomic data to probe the role of cell death *in vivo* during early tumour evolution.

Cell death removes cells, some of which have had offspring prior to death. This can lead to clones without extant parents (Fig. 4B), and the rate at which such ‘orphan clones’ arise can be used to infer the rate of cell death. We consider two metrics which quantify how frequently parental cells are removed from the population by cell death. The clone turnover gives the fraction of genotypes whose parental genotype is no longer extant in the tumour. The clade turnover gives the fraction of clades that coincide with their ancestral clade. (A clade is defined by a given mutation. Without backmutations, it comprises all individuals carrying this mutation.) These two metrics depend on the rate of cell death (relative to the birth rate) and on the mutation rate per cell division. Increasing the rate of cell death increases both clade and clone turnover, whereas increasing the mutation rate leads to a larger clone turnover only. These relationships have been determined analytically for a simple model of a growing population [46]. Inverting these relationships, the relative rate of cell death and the mutation rate can be determined from these metrics, see SI S5 and [46] for details and extensive tests.

Fig. 4C and D show the violin plots for the relative death rate and the mutation rate per cell division inferred in this way from the genotyped data of Ling *et al*. [36]. We find a rate of cell death nearly as high as the rate of birth (relative death rate 0.975 with 90% confidence interval [0.83, 0.995]). Hence during its early stage, the tumour was balanced nearly perfectly between growth and extinction. The inferred exome-wide mutation rate *µ* per cell division is 0.015 with 90% confidence interval [.002, 0.1]. This result is at variance with a recent analysis based on the distribution of mutational distances between samples. Werner *et al*. [47] find low rates of cell death across many cancer data sets. In SI S6 we show that this discrepancy arises because different rates of cell deaths are compatible with a given distribution of pairwise distances between samples.

To obtain the mutation rate per nucleotide we need to divide *µ* by the effective genome size, that is the number of sites in the genome that could have acquired detectable mutations. For many sites only few or no reads are retrieved during sequencing, so dividing *µ* by the size of the whole exome would underestimate the mutation rate. Ling *et al*. apply a set of criteria to filter out sites with poor coverage ([36], SI): A site must have 1) a total coverage of more than 150 reads when summed over all samples, 2) at least one sample with a coverage of 10 reads or more, and 3) a coverage of 6 reads or more in the normal tissue sample. We find that 4.8 *×* 10^5^ sites in the exome fulfill these criteria, and the resulting mutation rate estimate per nucleotide is 1.6 *×* 10^−8^ with 90% confidence interval [2*e* − 9, 1*e* − 7]. Comparing this result to a mutation rate estimate from cell cultures of healthy human fibroblasts of 2.7 *×* 10^−9^[**?**], this would mean the tumour mutation rate was enhanced by a factor of 5. We note that the relatively small size of the tree likely affects these estimates, and that analyses of larger trees in the future may lead to more accurate results.

### Mutational signatures and temporal heterogeneity

Somatic mutations in cancer tumours are caused by different mutational processes, which can be identified by their mutational signatures [48]. To investigate which processes were active in the tumours analysed here, we decomposed the mutational profiles of all three tumours into mutational signature components. Figure 5 shows that the strongest signature in all tumours is the single-base-substitution signature 22 (SBS22). This signature has been associated with exposure to the exogenous mutagen aristolochic acid [49], a herbal component used in traditional Chinese medicine. (The cancer patients of the Ling *et al*. and Li *et al*. lived in China.) To ask if there is a pseudo-temporal heterogeneity in the mutational processes, we divided the mutations from each tumour into clonal (early) and subclonal (late) mutations. While the Li *et al*. tumours maintain similar mutational signatures over pseudo-time, we find that the exposure to aristolochic acid of the patient from Ling et al. appears to have ceased over the course of tumour evolution: while the mutational signature SBS22 dominates the clonal mutations, it is absent in the subclonal mutations, see Fig. 5. The probability of not observing subclonal SBS22 due to sampling noise (*p*-value) was estimated to be less than 10^−4^, see SI S9.

**Figure 5:**
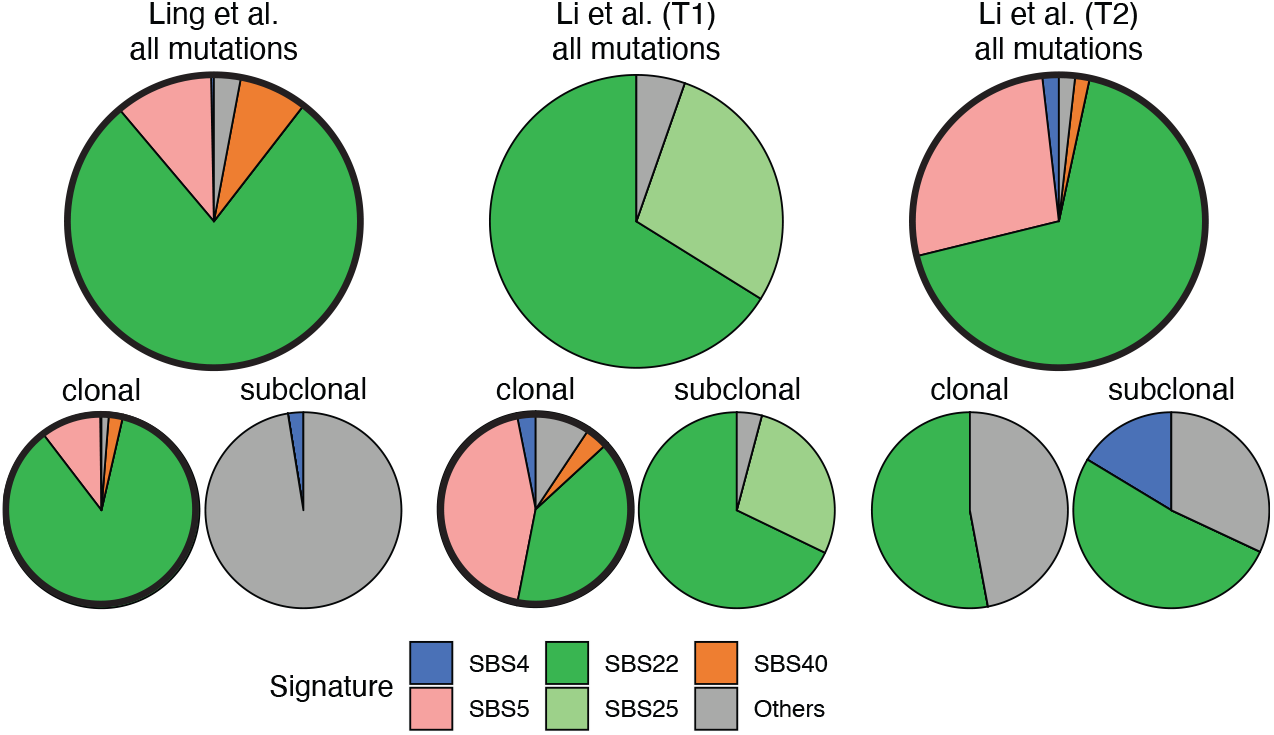
Mutational signature decomposition. Relative weights of single-base substitution (SBS) mutational signatures [48] in all three tumours were derived with SigNet Refitter where possible (highlighted pie charts), otherwise with non-negative least squares [50] (Methods). (left) Ling et al., (centre) tumour T1 of Li et al., (right) tumour T2 of Li et al. Top: all mutations, bottom: mutations stratified by their clonality. Signature SBS22 is associated with exposure to aristolochic acid. In the Ling et al. data, this signature is prominent among clonal mutations, but absent in subclonal mutations. Also shown are signatures with relative weight larger than 1% and attributed to endogenous mutational processes (SBS5 and SBS40), tobacco smoking (SBS4) and a signature that resembles SBS22 (SBS25). All other signatures were combined into a single category (“Others”).

### Population size

The results on the mode of evolution, migration, and cell turnover characterize the early stages of the tumours analysed here. The reason is that in an exponentially growing population, late-stage mutations have frequencies too low to be detected: Under neutral evolution and neglecting fluctuations in clone sizes, the final frequency (in the entire population) of a mutation that arose when a tumour consisted of *N* cells is given by *f* = 1*/N* [29]. A lower limit on the frequency *f* thus corresponds to an upper limit on the population size. In order to determine the population size the tumour had when the mutations we analysed here arose, we generalize this result to include the effects of spatial sampling and stochastic population dynamics, see SI S7. The higher the spatial resolution, the lower the whole-tumour frequency of a detectable mutation can be. For this reason, a high spatial resolution means that late-coming (low frequency) mutations can be detected. We find that the mutations detected in the samples with high spatial resolution (surface growth/volume growth analysis and genetic turnover) arose when the tumour consisted of about 80000 cells. The mutations in the whole-exome samples used to measure the dispersion of mutations arose when the tumour size was around 7000 cells.

## Discussion

### Modes of tumour evolution

We have used genetic data taken from hepatocellular tumours at high spatial resolution [36, 24] to analyse the early evolution of a tumour *in vivo*. We are not the first to link spatial tumour sampling with evolution: on a larger spatial scale, phylogenetic comparison of samples from primary and metastatic tumours has elucidated the dynamics of metastasis [50, 51, 52, 53]. Analyses of multiple samples from the same tumour have established intra-tumour heterogeneity and found neutral evolution in many cases [3, 19, 36, 34, 21] (see [54] for a critique), as well as instances of selection [35, 55]. The mode of growth of a tumour affects the resulting genetic diversity of a tumour [25, 26].

Here, we have used the spatial information contained in data based on hundreds of samples from single tumours. We found that the tumour evolved under uniform volume growth; there was no evidence of spatial constraints leading to radially outward growth. We also found a substantial turnover of cells with a rate of cell death comparable to the rate of cell division.

The rate of cell death being comparable to that of cell division implies that the tumour grew much more slowly than suggested by the rate of cell division. Instead, in its early stage, the tumour was balanced precariously between continued growth to macroscopic size and shrinkage: A small increase in the rate of cell death, or a small decrease in the growth rate would have caused the small net growth rate to become negative, possibly even leading to the extinction of the tumour.

The high spatial resolution also allows to track changes in the mutational processes over evolutionary time far more easily than in bulk data. Based on the high-resolution data of Ling *et al*. [36] we found a drastic difference between the mutational signature of early and late mutations. In this data, the dominant mutational signature in early mutations is associated with an exogenous mutagen (aristolochic acid) found in herbal traditional Chinese medicine and thought to cause liver cancer [56]. This signature is absent in late mutations, compatible with a discontinued dose of the mutagen. In the data set of Li *et al*. [24], the same signature is dominant both in the early and late mutations, compatible with a continuous dose of the mutagen.

### Mutation dispersion and sampling

We have found mutations in the tumour that are widely dispersed throughout the tumour; as a result they can have low frequencies in one particular sample but high frequencies in other parts of the tumour. We find the same level of dispersal of mutations in off-lattice simulations of growing tissues, where it is due to cells pushing each other out of the way as a part of the growth process. Hence, cells that were initially adjacent to each other and share a mutation can get pushed apart during the course of tumour growth and end up with a large distance between them. We found that this effect can be highly advantageous when assessing the mutations in a tumour from a limited number of samples: cells carrying a particular mutation can migrate into a spatial region that is later sampled. Mutations might thus be present at low frequency in a particular sample, because the cells carrying it migrated from afar, rather than because the mutation either arose late during the growth of the tumour [34] or was under negative selection [57].

Our results are based on three well-encapsulated hepatocellular tumours, resected and sampled in two and three dimensions [36, 24]. Potentially, different types of tumours or tumours at different stages may exhibit other growth modes. For instance, tumours that are not encapsulated might have a more heterogeneous environment, and evolve under surface growth. Also, the spatial and genomic resolution of any data set imposes limits on how late stage events can be observed. We estimate that the data analysed here capture the evolutionary dynamics up to 80000 cells. Increasing the genomic and spatial resolution will allow to identify mutations of lower frequency and thus characterize the *in-vivo* evolution beyond the early stages. Future techniques to sequence resected tumours at higher spatial and genomic resolution than currently possible will allow to trace key later-stage evolutionary changes like angiogenesis, effects of the tumour microenvironment, or the development of genetic instability.

## Supporting information

Supplementary Information (single file)

## Acknowledgments

This work was funded by the Deutsche Forschungsgemeinschaft (DFG, German Research Foundation) grant SFB1310/2 - 325931972. We acknowledge support of the Spanish Ministry of Science and Innovation through the Centro de Excelencia Severo Ochoa (CEX2020-001049-S, MCIN/AEI /10.13039/501100011033), and the Generalitat de Catalunya through the CERCA programme. We also acknowledge funding by the Spanish Ministry of Science and Innovation through grants PGC2018-100941-A-I00 and PID2021-128976NB-I00. This work was also partially funded by the FWF Austrian Science Fund (Erwin-Schrödinger postdoctoral fellowship, J4366). We thank Xuemei Lu and Chung-I Wu for discussions on the data sets [24] and [36], respectively, and Martin Peifer and Michael Lässig for discussions. Many thanks to Alison Feder and Nicola Müller for discussions on their SDevo algorithm.

