## Supplementary Information (single file) for "High-density sampling reveals volume growth in human tumours"

**S1 Computational modelling of volume and surface growth** **2**

**S8 Inference of clones** **37**

**S9 Subsampling of mutations and mutational signatures** **38**

#### S1 Computational modelling of volume and surface growth

We use a cell-based and off-lattice computational model of neutral spatial tumour growth in three dimensions. The model is based on a continuous-time many-type branching process well known from the population genetics of asexual reproduction [1], coupled to a simple model of spatial dynamics. Under this model, an individual cell can die, divide, and acquire neutral mutations at division. Cells inherit the spatial position from their parents, and a simple pushing algorithm to avoid overlapping cells is used to simulate the tissue mechanics. The model is implemented by a rejection kinetic Monte Carlo algorithm as outlined below. In this section, we introduce the dynamics of cell division and spatial displacement, specify the spatial sampling in Section S1.2, and in Section S1.1 we discuss the performance of this algorithm compared to rejection-free kinetic Monte Carlo algorithms such as the Gillespie algorithm. The code to the model is available as a Julia package at `https://github.com/aangaji/TumorGrowth`.

In our model, the growth dynamics of a cell can differ depending on its local environment. Under volume growth, the birth and death rates of cells are independent of their spatial position, under surface growth we make the rate of cell birth depend on the local cell density. Specifically, the rate $b$ at which a particular cell divides depends on the local cell density $\rho$ via a function $b(\rho)$. The local density $\rho$ at a position $\mathbf{x}$ is computed as a weighted sum over cells $i$ in proximity of $\mathbf{x}$, each contributing a Gaussian weight by their distance

$$\rho(x) = \sum_{\{i \mid |\mathbf{x}-\mathbf{x_i}|<7[\text{cell radii}]\}} \frac{1}{\sigma\sqrt{2\pi}} e^{-(\mathbf{x}-\mathbf{x}_i)^2/(2\sigma^2)} . \tag{S.1}$$

We set the width of the Gaussian $\sigma$ to 4 cell radii and evaluate the sum over $i$ over cells within 7 cell radii. This dependency of the cell birth rate on the local neighborhood is the reason why updating the rates of all cells at each time step (as required by a rejection-free method) would be prohibitively expensive.

The rejection kinetic Monte-Carlo algorithm for a density-dependent off-lattice spatial population dynamics is as follows:

1. Pick a cell $i$ uniformly
2. Update cell birth rate $b_i = b(\rho_{nn})$
3. Draw $r$ uniformly from the interval $(0, b_{\max} + d)$
   - $r < d$ : cell dies
   - $d < r < b_i + d$ : cell divides
     → each cell draws new mutations $m \sim \text{Poisson}(\mu)$
     → resolve overlaps by pushing
   - $b_i + d < r < b_{\max} + d$ : cell skips its turn
4. Increment time by $\Delta t \sim \text{Exp}(\frac{1}{(b_{\max}+d)\cdot N})$

The function $b(\rho)$ is chosen such that rate of birth is maximal at $\rho = 0$ and zero at a density threshold $\rho_c$, and (for simplicity) decreases in a straight line with $\rho$, see Figure S1A. This dependence encodes potential effects neighbouring cancer cells may have on the division rate of a cell, for instance due to the buildup of toxic metabolites or mechanical stress. At high values of $\rho_c$ (relative to the maximum density achieved without overlapping cells, see below), the birth rate is independent of the local cell density, at low values of the threshold the birth rate depends on the local density $\rho$ and becomes zero at $\rho_c$. Thus, by setting a high threshold $\rho_c \to \infty$ the model produces volume growth, and a low value of $\rho_c$ leads to surface growth. Under surface growth, the growth rate is zero or nearly zero throughout the tumour, except at the surface, where the local cancer cell density is smaller than in the tumour bulk. By tuning the parameter $\rho_c$, we can continuously vary the mode of growth between surface growth and volume growth.

The cell death rate and the mutation probability at division are the same across all cells. As a result, under volume growth cells divide and die independently of their spatial position and the position of other cells. The population thus evolves like a mixed population would, although every cell has a distinct position in space and this position is inherited from one generation to the next.

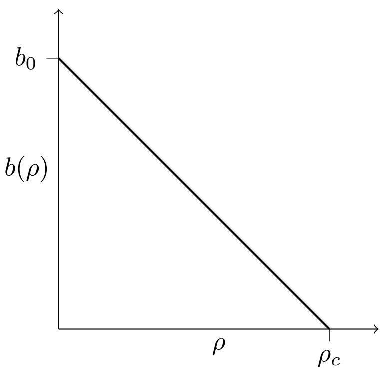

**Figure S1: Density-dependent birth rate.** A schematic plot of the birth rate $b(\rho)$ decreasing with the local density of cells $\rho$, see text.

During the simulation, cells are uniformly drawn from the population for birth and death. Importantly, this approach allows us to update a cell's birth rate only when it is selected for division as the total rate of events - comprising birth, death and null-event - is uniform across cells and constant $b_{\max} + d$. On the other hand, this means that a cell with a low birth rate might also be rejected (null-event), i.e. neither die nor divide.

After each draw of a cell, time is incremented. Time steps are exponentially distributed random variables with the mean total rate given by $\tau \sim \text{Exp}\left(\frac{1}{(b_{\max}+d)\cdot N}\right)$, where $b_{\max} = b(\rho = 0)$ and $N$ is the current population size.

At division, a new cell is placed randomly on the surface of the dividing cell. The resulting overlaps between cells are resolved by pushing overlapping cells away from each other. This step of the algorithm is computationally expensive, so keeping track of immediate neighborhoods is essential, see Section S1.1 on the computational performance.

The details of the pushing algorithm are as follows: First, the close neighbourhood of the newly placed cell is searched for overlaps. Any overlapping cell is pushed away from the new cell by the overlap plus a small margin which acts as a buffer and avoids many tiny overlaps. The cell is then added to a queue for subsequent iterations. Neighborhoods are shuffled such that there is no particular order in which cells are pushed. After each overlapping cell in the neighborhood is pushed and added to the queue the search for overlaps is repeated on the first cell in the queue, adding new pairs to the queue, which are in turn iteratively resolved until the queue is empty. In this 'width-first' approach, overlaps are first resolved within a given neighborhood and new neighborhoods are appended to the queue, as opposed to a 'depth-first' approach which follows a sequence of pushes by immediately checking a pushed cell's neighborhood for new overlaps and adding these to the front of the queue (which then operates like a stack). This can be efficiently implemented by recursion but has no noticeable performance advantage and results in the same growth patterns.

A commonly used procedure to increase the computational performance in such collision models is to rasterize continuous space. Each cell is assigned to a position on a grid which has a lattice constant of two cell radii. When searching for neighbours - both when pushing and updating the density dependent birth rate $b(\rho)$ - a quick selection of the close (Moore) neighbourhood on the grid returns candidates for which to measure the precise distance. After a cell is either pushed or dies, its position on the grid is updated.

At cell division, each of the two cells - parent and offspring - acquires new mutations whose number is drawn from a Poisson distribution with mean $\mu/2$. These mutations are taken to be neutral and do not affect the birth or death rates.

Our model combines a simple computational off-lattice framework combining genetic mutations in a growing population with a simple tissue dynamics. Additional features, such as selection or different growth laws (for instance Gompertzian growth) can be implemented easily. The model avoids the well-recognized problems that arise in lattice-based models: volume growth on a fully occupied lattice necessitates artificial steps like expanding all distances by some factor while keeping the lattice spacing constant [2], or moving a column of cells one step in one particular direction to generate empty sites [3]. However, the computational cost of the off-lattice model means that we can only simulate populations of a few ten thousand cells, compared to billions of cell that can be simulated on a lattice [4].

#### S1.1 Algorithm performance and CPU runtimes

Kinetic Monte Carlo (KMC) is a stochastic simulation of a discrete Markov process under continuous time. Under KMC, cells are picked to potentially replicate or die with a certain probability, and after each event time is incremented by a stochastic waiting time. In a rejection or null-event KMC (rKMC) scheme, the cell that has been drawn for update might not change its state (e.g. replicate or die) after a given time step, in contrast to rejection-free KMC (rfKMC) where all events lead to a change of state.

The advantage of rKMC compared to a rfKMC algorithm is that one does not need to know the rates of all cells at each step. Instead, the drawn cell's rates are updated just in time and normalized by a constant, maximum rate, which is larger than each cell's total rate $b_i + d \leq b_{\max} + d$. The maximum rate should be as small as possible to minimize the number of null-events (rejections) and can be conveniently set to $b_{\max} + d$ using the maximum birthrate at $\rho = 0$.

Rejection-free methods, on the other hand, require global updates of the relevant rates at every step. In our model, this would involve computing the local density at every cell. Such an update scales with population size and would be computationally prohibitive. Rejection KMC, on the other hand, allows for local updates and only computes the birth rate $b$ of a single cell per step, at the cost of null-events. However, these null-events do not involve the computationally expensive pushing step (cell movements) and add little to CPU runtime even at low density thresholds $\rho_c$ and high death rates $d$. For a detailed review on the two classes of KMC algorithms, see [5]. The rKMC approach has already been used in non-spatial simulations of populations of cancer cells [6].

A 3d simulation to 40000 cells at turnover rate $d/b = 0.2$ takes 5 minutes under surface growth but 16 minutes under volume growth on an AMD Ryzen 5 3600 CPU at 3.6 GHz. At high turnover rate $d/b = 0.975$ a simulation under volume growth takes from 19 up to 28 minutes. The time required for a simulation is almost fully spent on moving cells to resolve overlaps. Drawing birth and death events on the other hand is very fast even at high death rates $d$. High death rates mainly prolong run times by increasing the number of births necessary to reach the final tumour size which entails more pushing steps. This is particularly noticeable for volume growth simulations, where cells predominantly divide and push from within the tumour volume, whereas overlaps under surface growth are more quickly resolved because they affect fewer cells.

#### S1.2 Spatial sampling in simulations

The tumour data used here stems from a large number of samples taken from histopathological sections of resected tumours [7, 8]. To compare the results of our simulations with this data, we take samples from the simulated populations in a way that mimicks the empirical sampling.

Ling *et al.* take samples from a single planar section through the tumour [7]. To mimick their sampling scheme, we first cut a two-dimensional plane out of a three-dimensional simulated cell population, and then take small, circular 'punch' samples on a triangular lattice with adjustable number of samples, sample size, and spacing. Given the desired number of samples $n$, we consider a triangular lattice on the plane with a suitable lattice spacing and place samples on the lattice points. The triangular lattice has the highest packing factor $\frac{\pi}{2\sqrt{3}}$ compared to other lattices and fits the most samples into a given area. Using the packing factor we estimate the lattice constant such that $n$ primitive cells of the lattice cover the area of the plane. The number of samples can fluctuate a little from run to run, depending on the precise shape of the tumour and the orientation of the lattice relative to the tumour. The number of cells in the plane scales poorly with total population size, and at 40000 total cells, the plane contains about 2600 cells given a narrow plane width of 3 cell radii. To ensure that there is free space between samples (like in [7]), we set the number of cells per sample to 5 when taking 285 samples, but have 20 cells or more per sample when 23 samples are taken. The numbers of samples are picked to be the same as in Ling *et al.* for the spatially-resolved data obtained by genotyping and the genomically resolved date obtained by whole-exome sequencing (WES), (see main text and Methods: Spatially resolved data). We always compare the results obtained from this spatial sampling in the cross section of 3d simulations to sampling on 2d simulations of the plane, where for a population size of 10000 cells we take larger samples of 20 cells for 285 samples. The results, however, do not depend much on this size differences of samples and cell populations but rather on the frequency resolution of mutant frequencies (only mutations with frequency larger than a cutoff set by the sequencing depth are retained) and the number of samples taken.

Data from samples taken in 3d from two hepatocellular carcinomas (tumours T1 and T2) from a single patient is discussed in a recent publication by Li *et al.* [8]. Similar to Ling *et al.*, hundreds of samples were taken from these tumours (169 from tumour T1 and 160 from T2), but different slices out of the upper hemispheres of the two tumours were taken before taking samples within each slice. Again some of these samples were then analysed by whole-genome sequencing (16 in T1 and 9 in T2), and the remaining 153 and 151 samples were genotyped based on the mutations found by WGS. The multi-region sampling scheme of Li *et al.* is different from Ling *et al.* in that it takes samples from several planes of a tumour hemisphere as opposed to sampling from only a single cross section. To imitate the sampling scheme of Li *et al.*, we consider slices in the upper hemisphere of our 3d simulated tumours at distances specified by Li *et al.*. From these slices we then take small 'punch' samples at the coordinates specific to the tumours T1 and T2, both for WGS and genotyped samples. Samples in the two tumours are placed rather unevenly, in contrast to the more uniform sampling by Ling et al. For this reason, we place the samples in simulations close to the empirical positions of the samples given in Li *et al.*.

### S2 Surface and volume growth: direction of mutants

We construct a simple metric to distinguish surface growth and volume growth. It is based on the directions of newly occurring mutations with respect to their parental background. If the tumour predominantly grows on the surface, then new mutants appear radially outwards relative to their parental clone. All pairs of parental and offspring cells are assigned a direction angle and a statistical weight such that every mutation $m$ contributes equally to the metric.

#### S2.1 Algorithmic description

The metric is based on lineages of clones. Clone A is an ancestor of clone B if A's mutations are a true subset of B's mutations. We therefore need to determine the set of clones. This is straightforward for single cells where clones are genotypes. Samples containing many cells, however, may display a mixture of different clones. We use a simple clustering scheme where mutations that coincide across samples are clustered into clones, see Section S8. We compared the results of this clustering scheme also to those of the LICHeE algorithm for the inference of multi-sample lineages [9], which produces very similar sets of clones and virtually identical final results.

The direction angles of new mutations are computed as follows:

1. Infer clones and their lineage (see Section S8)
2. for each sample $s_k$ that has cells of clone $\{m_i\}$
   for each ancestral clone $\{m_j\} \subset \{m_i\}$ and each sample $s_l$ that has cells of the ancestral clone
   (a) determine distances $\vec{\Delta}_{k,l} = \vec{p}_k - \vec{p}_l$ between $s_k$ and $s_l$
   and $\vec{\Delta}_{l,\text{cm}} = \vec{p}_l - \vec{p}_{\text{cm}}$ between $s_l$ and the tumour centre of mass
   (b) skip if $|\vec{\Delta}_{k,l}|$ is larger than the distance between $s_l$ and the surface
   (c) determine the direction angle $\theta_{k,l}^{m_i,m_j} = \angle(\vec{\Delta}_{l,\text{cm}}, \vec{\Delta}_{k,l}) \cdot \text{sign}(\Delta^x_{l,\text{cm}}\Delta^y_{k,l} - \Delta^x_{k,l}\Delta^y_{l,\text{cm}})$
   (zero if vectors are aligned i.e. offspring lies radially outward relative to the ancestor)
   (d) assign weight $w_{k,l}^{m_i,m_j} = \left(\sum_{m_j}\sum_{s_k\in\{m_i\}}\sum_{s_l\in\{m_j\}} 1\right)^{-1}$ (one over the number of valid pairs)
3. plot the histogram of $\theta_{k,l}^{m_i,m_j}$ with weights $w_{k,l}^{m_i,m_j}$
4. from $c = \left\langle w_{k,l}^{m_i,m_j} \exp(i\theta_{k,l}^{m_i,m_j}) \right\rangle$ compute mean angle as $\arg(c)$ and radius as $|c|$

**Step 1.** In words, when a mutation $m_i$ enters the population, it defines a new clone $\{m_i\}$ whose genotype consists of the new mutation $m_i$ and all its ancestral mutations. The clone $\{m_i\}$ is the most recent common

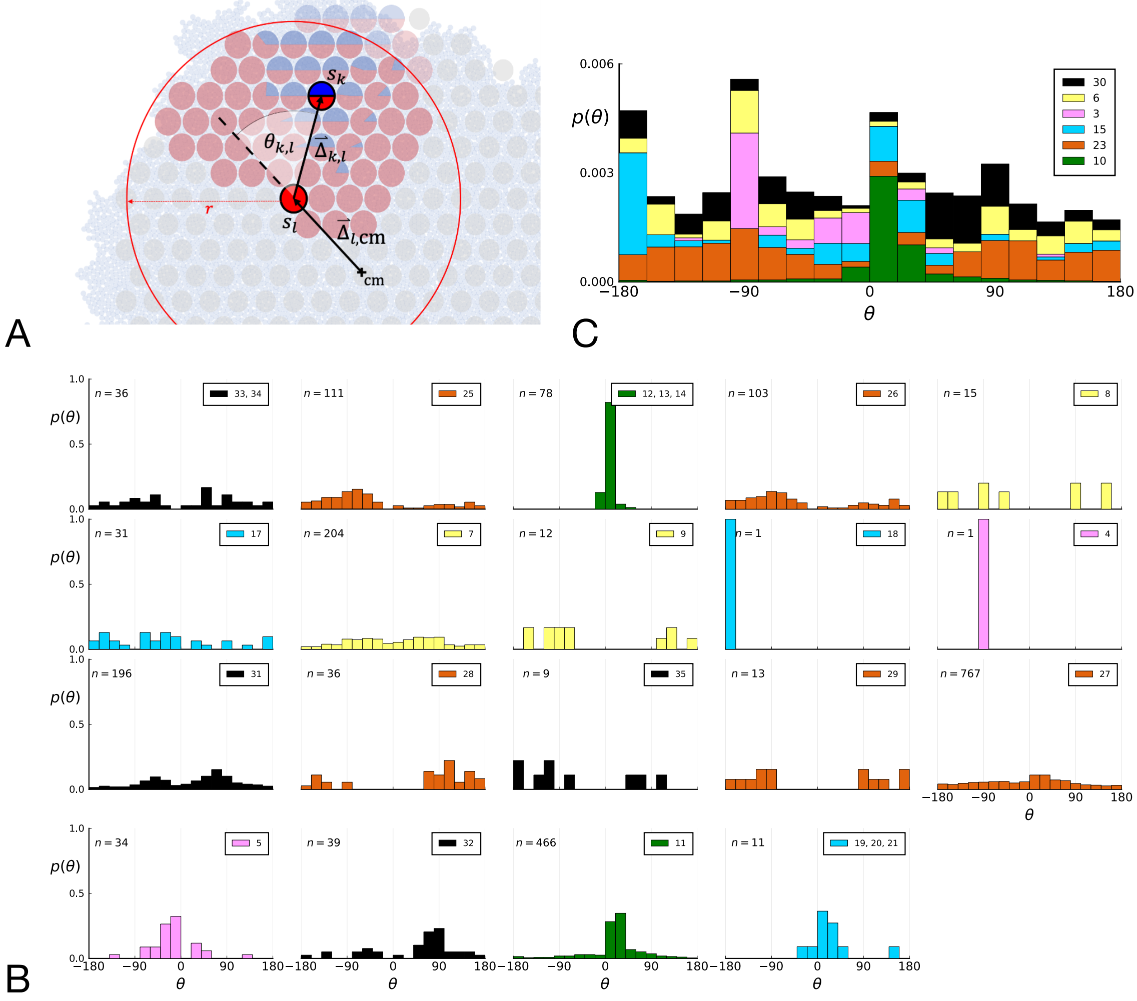

**Figure S2: Parent-offspring direction angle algorithm illustration and contributions of different clones.** A This cutout of Fig 2A showcases the different variables used in step 2 of the algorithm to calculate the angles $\theta$. A new mutant clone with mutations $\{m_i\}$ indicated in blue appears and grows radially outward on a red parental background with mutations $\{m_j\} \subset \{m_i\}$. For a given parental sample $s_l$ belonging to the parental background, we consider every sample $s_l$ belonging to the offspring clone within a distance $r$ from $s_l$, where $r$ is the distance between $s_l$ and the tumour surface. $\theta_{k,l}$ for the pair $s_k$ and $s_l$ is the angle between the arrow $\vec{\Delta}_{l,\mathrm{cm}}$ pointing from the center of mass cm to $s_l$ and $\vec{\Delta}_{k,l}$ connecting $s_l$ to $s_k$ as calculated in step 2c. A weight is assigned to $\theta_{k,l}$ such that the total weight of all pairs where the blue clone is the offspring is 1; this way each clone contributes equally to the distribution of $\theta$. B The distribution of angles $\theta$ in the Ling *et al.* data shown Fig. 2 B of the main text comprises contributions from different clones. These contributions are shown here separately for each clone. On each subplot, the top-right label gives the private mutations of the clone and on the top-left label gives the number of parent-offspring pairs contributing to the histogram. The color indicates which of 6 distinct clades the clone belongs to. Crucially, there is no bias for a particular value of $\theta$ across clones. An artefact of giving equal weights to clones are the sharp peaks when there are few parent-offspring pairs. In particular clones 4 and 18 (panel 9 and 10) each have only 1 parent-offspring angle. C This figure shows the contribution of the 6 distinct clades (clade colours as in Subfigure B) to the total angle distribution in Fig. 2B of the main text. Histograms of subfigure B of clones belonging to the same clade are added and the resulting clade histograms are stacked (weights of the stacked histogram are divided by the total number of clones for normalisation).

ancestor of all clones containing mutation $m_i$. We ask how the radial position of this clone relates to its ancestral clones. (2) Given an ancestral clone $\{m_j\}$ we take one sample that has cells of the offspring clone $\{m_i\}$ and one that has cells of the ancestral clone, (2a) draw a line between the two and (2c) measure the direction angle $\theta$ between this line and the line connecting the sample with the ancestral clone to the tumour centre of mass. If tumour growth occurs on its surface (interface with normal tissue), the offspring grows outward, and we expect this angle to be biased towards zero. The direction angle is symmetric with respect to the line between parent and centre of mass, so to recover the whole $-180°$ to $180°$ range we multiply it by a term which distinguishes clockwise from counterclockwise. We repeat this measurement for all pairs from the two clones and again with all other ancestral clones following the lineage backwards in time. (2d) The weight associated with each direction angle $\theta$ is one over the total number of ancestor-offspring angles for clone $\{m_i\}$, such that, after summing over all samples with cells of the ancestral clone, every mutation contributes with equal weight.

In the case where frequencies of clones are known in each sample, one can assign a weight $w_{k,l}^{m_i,m_j} = f_k^{m_i} \cdot f_l^{m_j}$ to the sample pair $k \in \{m_i\}, l \in \{m_j\}$, where $f_x^m$ is the frequency of clone $\{m\}$ in sample $x$. The product of frequencies $f_k^{m_i} \cdot f_l^{m_j}$ is proportional to the number of offspring-ancestor pairs of cells between the two samples. Dividing each weight $w_{k,l}^{m_i,m_j}$ by the sum $\sum_{k,l,m_j} w_{k,l}^{m_i,m_j}$ over all weights involving clone $\{m_i\}$ as offspring again imposes a normalization where each mutation contributes equally to the distribution of direction angles.

**Step 2** Part b) accounts for a simple geometric effect: If a sample containing offspring is placed randomly around a sample containing ancestors, the direction angle $\theta$ is uniformly distributed, however, more cells lie inward than outward from any point except the centre of the disk resulting in a purely geometric bias towards $\pm 180°$. The restriction to only consider offspring samples within the distance between ancestor sample and tumour surface mitigates this bias by only considering as many samples inward from the ancestor sample as lie outward from it.

**Step 3** Directly plotting the weighted histogram of direction angles $\theta$ gives a picture of the radial direction of growth across all mutations, (4) but one can also compute a mean angle and radius as summary statistics of the distribution. The expression for $c$ is the weighted sum over all unit-length 2d vectors pointing in the directions $\theta_{k,l}^{m_i,m_j}$, resulting in an average vector pointing towards the direction of growth and whose length quantifies the strength of this bias, i.e. a radius $|c| = 1$ means that all offspring have a direction angle $\bar{\theta} = \arg(c)$ relative to the ancestor, while $|c| = 0$ corresponds to a uniform placement of the offspring relative to the ancestor (in this case $\bar{\theta}$ is indeterminate).

We compare 3d simulations of 40000 cells under volume growth ($\rho = \infty$) and surface growth ($\rho = 6$) both for sampling single cells and with a sampling scheme that imitates the experimental data of Ling *et al.* For each simulated tumour, we take a thin planar cross-section through the centre of mass. (The case of 3d sampling is treated in Section S2.2.)

For single cells, the distribution of $\theta$ can be measured directly for the single cells in the plane, resulting in many offspring-ancestor pairs. To study the effect of sampling, we employ the spatial sampling scheme described in Section S1 and take 285 samples of 5 cells each from 2d cross-sections of the 3d simulated tumour. The same sampling is repeated on 2d simulations without taking a cross section allowing for larger sample sizes of 20 cells. Finally, we compute the direction angles $\theta_{ij}^{\{m_i\}}$ and weights $w_{ij}^{\{m_i\}}$ as described above. The resulting distributions are shown in Figure S3 for 2d slices of 3d simulations and in Figure S4 for 2d simulations (where larger numbers of cells can be achieved). In both cases, we consider both the uniform sampling of single cells from the tumour and the sampling scheme mimicking [7] described in Section S1.

Both 2d and 3d simulations of surface growth clearly display the expected bias in the distribution of direction angles, compared to the flat distribution under volume growth.

The distribution of direction angles is very similar in 2d and 3d (comparing the left plots for single cells in figures S3 and S4 and similarly the right plots for samples). On the other hand, between single cells and samples in each figure, showing a small loss in signal strength (surface growth peak against the volume growth background) due to sampling noise and errors in the reconstruction of parent-offspring clone relations from samples.

Figures S3 and S4 show that the signal for surface growth can also be detected in a regime of high cell turnover (high rate of cell death $d$ compared to the birth rate). The distributions of direction angles $\theta$ in the low turnover regime are not shown here, they have the same shape as for high $d$ but show less sampling noise.

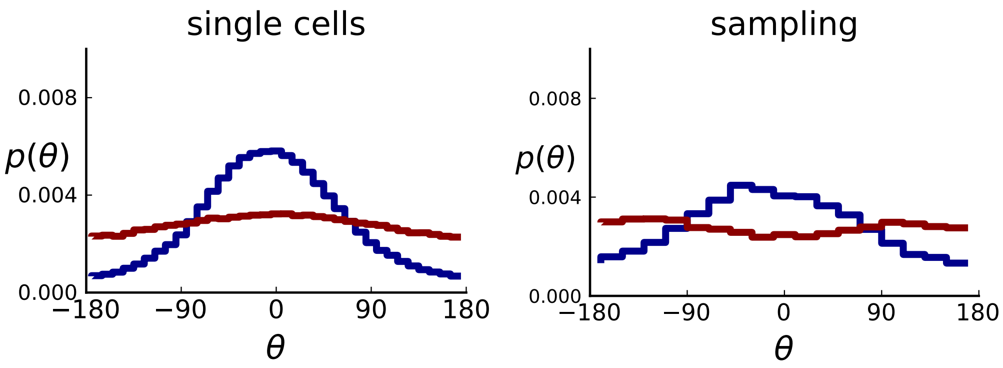

**Figure S3: Distributions of parent-offspring direction angles $\theta$ for 3d simulations under volume and surface growth.** Direction angles are taken from single cells (left) and the 2d spatial sampling scheme (right, see Section S1). Distributions of parent-offspring direction angles $\theta$ for cross-sections of 3d simulations of 40000 cells at constant division rate $b = 1$, mutation rate $\mu = 0.3$, as well as cell death rate $d = 0.8$ and $d = 0.4$ under volume growth (red, $\rho = \infty$) and surface growth (blue, $\rho = 6$), respectively. On the left, direction angles are determined for single cells (these curves are shown in Fig. 2 of the main text), whereas on the right, we additionally apply a spatial sampling with 285 samples. The cell death rate $d$ is set lower in surface growth simulations because also the rate at which cell divisions take place is much lower under surface growth due to the reduced growth in the tumour bulk. This leads to frequent population extinctions at higher rates of cell death $d$.

#### S2.2 Direction of mutants in the 3d sphere

So far, we have restricted ourselves to a planar cut through the center of a 3d tumour, motivated by the data from Ling *et al.* which was sampled from such a cross section. However, the definition of the direction of mutants can be readily extended to single cells and samples taken in 3d. An example such data is Li *et al.* [8].

The algorithm to determine ancestor-offspring direction angles as presented before is the same in 3d, with the exception that angles are not given a sign in step 2b) of the algorithm. The sign was computed relative to the $z$ axis orthogonal to the plane which defined an orientation to the direction angles and distinguished $0 - 180°$ clockwise from counterclockwise. This notion of orientation does not exist in the sphere.

More importantly, the most useful metric to quantify direction of mutants and distinguish surface from volume growth in 3d is not simply $\theta$ as on the 2d plane. To find the corresponding metric we ask what function of $\theta$ is uniformly distributed on the 3d sphere. To this end, we consider $\theta$ the polar angle in spherical coordinates. For isotropic growth of new cells the distribution of $\theta$ is specified by the volume element in spherical coordinates, and hence proportional to $\sin\theta$. Correspondingly, its indefinite integral $-\cos\theta$ and hence also $\cos\theta$ is uniformly distributed on the interval $[-1, 1]$. (Since we are looking for a function $f(\theta)$ that is uniformly distributed, the transformation of probabilities gives $\sin(\theta)d\theta = df$ and hence $\sin(\theta)d\theta = \frac{df}{d\theta}$.)

Figure S5 showcases these results using 3d simulations and sampling of single cells. Under volume growth, $\cos\theta$ shows a uniform distribution (red line). Surface growth, on the other hand leads to a bias towards the direction of the pole, $\theta = 0$, and thus a bias towards $\cos\theta = 1$ (blue line).

The set of samples taken in 3D by Li *et al.*, see S1.2, comprises samples that were whole-genome (WG) sequenced as well as samples that were genotyped. The sampling takes places at a lower spatial density than [7] and read counts are not reported for individual SNVs. In both T1 and T2, the spacing between neighboring samples is $8\% \pm 2\%$ of the tumour diameter whereas in the considerably larger tumour of Ling *et al.* (3.5cm compared to the 1.5cm of T1 and 2.5cm of T2) samples were taken at median distance of $4\% \pm 1\%$ of the tumour diameter. Measured in tumour diameters, the sampling density in Li *et al.* is about half that of Ling *et al.*, mainly due to sampling in 3d where the same number of samples is more spread out than in the plane. While 3d sampling may give a global picture of the intra tumour heterogeneity it comes at the cost of reduced spatial resolution.

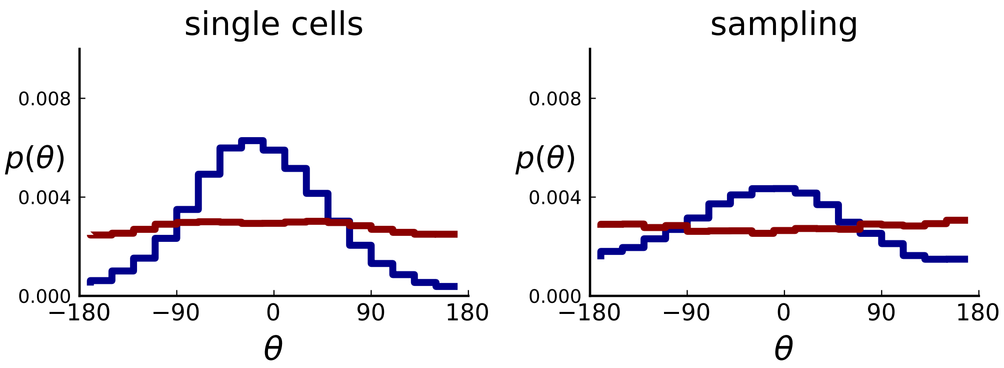

**Figure S4: Distributions of parent-offspring direction angles $\theta$ for 2d simulations under volume growth (red) and surface growth (blue).** Direction angles are taken from single cells (left) and the spatial sampling scheme (right). Parameters are as in Fig. S3 except that populations are grown to 10000 cells, no cross section is taken, and individual samples consist of 20 cells.

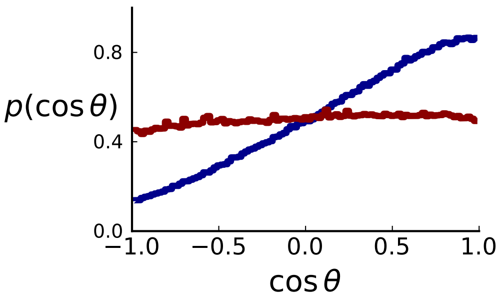

**Figure S5: Distributions of parent-offspring direction cosines $\cos(\theta)$ for single cells in 3d simulations under volume (red) and surface (blue) growth.** Direction angles $\theta$ are taken from single cells in the spherical tumour. Simulation parameters are as in S3 except that no cross section is taken.

However, Li *et al.* genotype a larger number of mutations than Ling *et al.* - 906 for T1 and 565 for T2 - because WGS targets a larger part of the genome than WES and they apply less stringent filtering criteria for SNVs compared to Ling *et al.* (see our discussion of the WES data in Section S4).

As explained in Section S8 on clonal inference, given the larger genotyping set of mutations (compared to the Ling *et. al* data), we use the LICHeE tool to determine clones, before identifying the inferred clones within the larger set of genotyped samples, which provide a higher spatial resolution. We then measure the distribution of ancestor-offspring direction cosine $\cos\theta$; the results are shown in figure S6. For comparison with simulations, we perform a spatial sampling of planar cuts through simulated tumours in 3d under surface and volume growth. We use the coordinates of planes and samples from tumours T1 and T2 to mimick their sampling (and potential spatial inhomogeneities therein) in our simulations. First, we extract samples for sequencing at the coordinates of WGS samples before taking samples at the coordinates specified for genotyped samples [8] to confirm the presence or absence of mutations found in the first set of samples, see Section S1.2 on simulated sampling. For the mutations found in the first 'sequencing' set we measure the distribution of $\cos\theta$, see Fig. S7

The simulation results show that sampling at the density of the Li *et al.* data (which is much lower than in the planar sampling by Ling *et al.*) introduces a radial weak inward bias under volume growth.

The inward bias also affects the distribution under surface growth - a similar effect as in sampling from the cross Section S4 and S4. This makes it harder to distinguish surface and volume growth in simulations mimicking the sampling in Li *et al.* in Fig. S7.

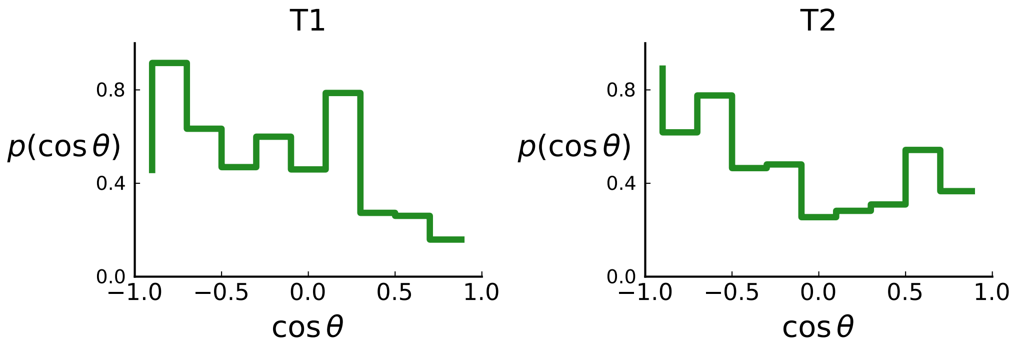

**Figure S6: Distributions of parent-offspring direction cosines** $\cos(\theta)$ **for the Li *et al.* 3d sequencing data.** Direction angles $\theta$ are computed from high-spatial-resolution (genotyped) samples of the tumours T1 (left) and T2 (right), after having applied the clustering scheme of Section S8 to obtain clones from the list of sample genotypes.

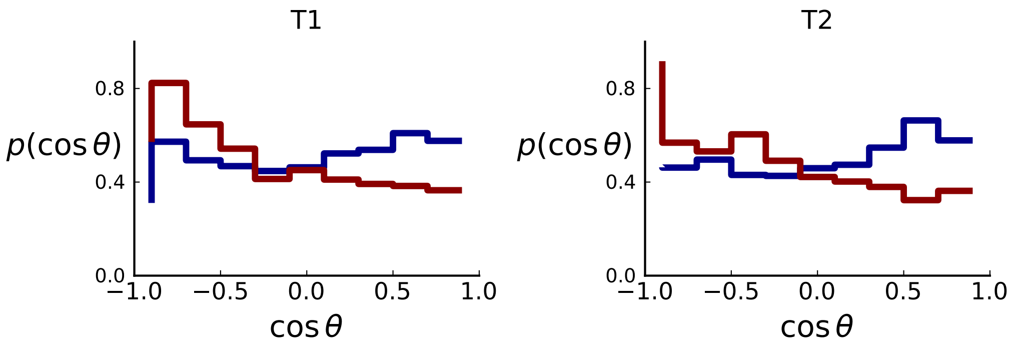

**Figure S7: Distributions of parent-offspring direction cosines** $\cos(\theta)$ **for sampling mimicking the spatial sampling of Li *et al.* [8] of 3d simulations under volume (red) and surface (blue) growth.** Direction cosines are taken from samples in layers of 3d tumour hemispheres following the sample coordinates of Li *et al.* for tumour T1 (left) and T2 (right) for direct comparison with figure S6. Simulation parameters are as in Figure S3, but no cross-section is taken. Volume growth shows a small downward trend, corresponding to a small radially inward bias. This may be due to the geometric effect discussed in Section S2.1 (Step 2).

#### S3 Mutation density on rings

Surface growth and volume growth can also be distinguished by looking at the accumulation of mutations with increasing distance to the tumour centre of mass, without the need to explicitly infer parent-offspring relationships. Under volume growth, mutations are distributed uniformly in space, whereas under surface growth cells at the surface accumulate mutations at a constant rate as the population grows radially. The latter results in an increasing mutation density (per volume or per area) as a function of distance from the centre of mass.

In this section, we look at the mutation density as a function of distance. We plot the number of mutations found within a ring of radius $r$ and width $w$ against $r$ and normalize by the number of cells or samples in the ring. To this end, we

1. set the origin to the tumour centre of mass
2. select the ring width $w$ at least larger than the distance to nearest neighbours
3. for each ring with radius $r_i$
   (a) take all samples within $r_i$ and $r_i + w$

(b) count the mutations in each sample and sum up all counts

(c) divide the total count by the number of samples in the ring to obtain the density

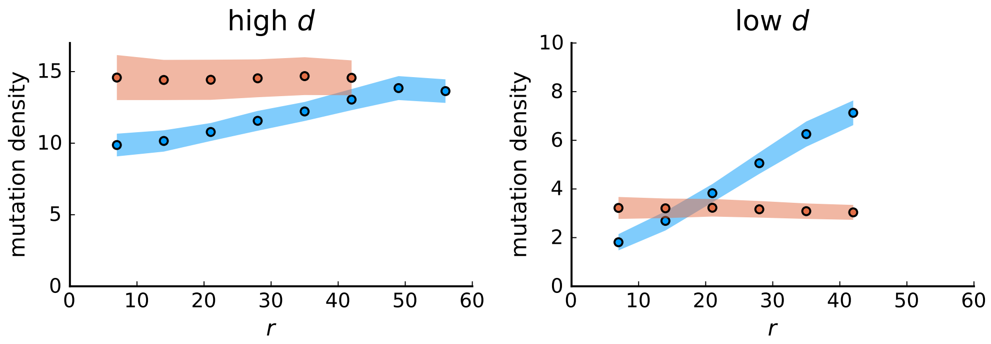

**Figure S8: Mutation density curves for sampling in 3d simulations.** We show the mean mutation density within a ring of radius $r$ and width $w = 7$ (see text). Each curve is averaged over 20 simulations, coloured ribbons indicate the standard deviations from the mean. For each simulation under volume growth (red, $\rho = \infty$) and surface growth (blue, $\rho = 6.0$), we take a cross-section and apply a spatial sampling with 285 samples (high spatial resolution) as described in the text. The right hand side shows simulations with zero death rate $d = 0.0$. On the left, (relative) death rates are high, $d = 0.8$ for volume growth and $d = 0.4$ for surface growth (see also Fig. S4).

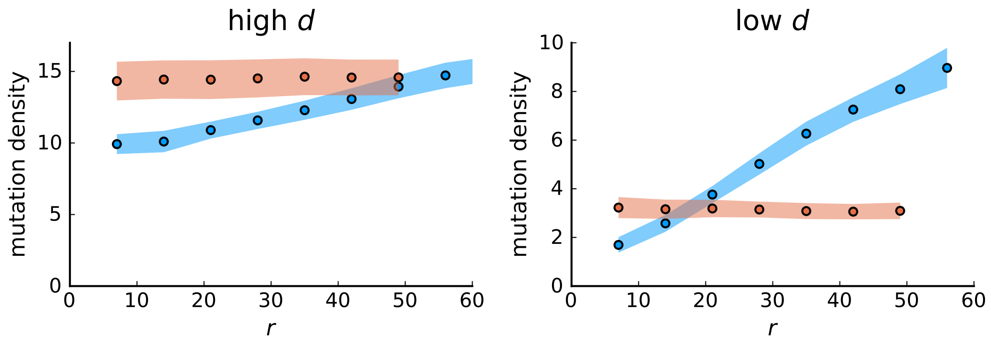

**Figure S9: Mutation density curves for single cells in 3d simulations.** Same as in Fig. S8 but for single cells under volume (red) and surface growth (blue) at high (left) and low (right) death rates.

To mimic the sampling scheme in [7], we take a thin planar cross-section through the centre of mass as described in Section S1. Given the sample number of circa $n = 285$ samples used throughout, the nearest neighbour distance is always below 7 cell diameters, which is why we choose the ring width to be 7 cell diameters for both single cells and samples.

We model stochastic effects in sequencing by setting a low coverage of 5 total reads per mutant site per sample and drawing alternate and reference reads from to a binomial distribution with the expected number of mutant reads given by the mutation frequency.

We then measure mutation densities, either for the full population of single cells in the plane or under the sampling scheme.

The mutation density versus ring radius $r$ is flat for volume growth (see Figure S8) because all cells divide at the same rate regardless of their radial position. The mutation density therefore assumes a value given by the number of mutations a cell is expected to accumulate until the given tumour size $N$: After $T$ generations (cell divisions) each cell has accumulated $m \sim \text{Poisson}(T\mu)$ mutations. Notably, a cell that is born after $T_0$ generations inherits its parents mutations and ends up with $m \sim \text{Poisson}(T_0\mu) + \text{Poisson}((T - T_0)\mu) =$

Poisson($T\mu$) mutations. Consider a ring of radius $r$ with $n$ cells. Due to the high dispersion of cells under volume growth, the number of divisions $T_i$ is independent for different cells $i$ in the ring at radius and the number of mutations is again Poisson-distributed $m_r = \sum_i m_i \sim$ Poisson($\sum_i T_i\mu$). The expected mutation density $\langle\rho_r\rangle = \langle m_r/n\rangle = \log N/(1-d/b)\mu$ and standard deviation std$(\rho_r) = \sqrt{\langle\rho_r\rangle/N_r}$ describe the mutation density curves under volume growth.

For surface growth, the mutation density increases as a straight line, see Figure S8. At zero death rate, the mutation density is compatible with a linear dependence on the radial distance. However, at a finite death rates there is a clear vertical offset, with a finite mutation density at zero distance. This is because at a finite death rate, cells die uniformly over the tumour volume, and this allows for the division of other cells in their vicinity. Hence, the higher the rate of cell death, the more cells divide also in the bulk of the tumour; the effect of spatial constraints thus diminishes with the death rate. As a result, for a model of surface growth based on local tumour cell densities, the distinction between the surface and volume growth disappears at high rates of cell death. Importantly, this is not a failure of a metric designed to distinguish between different growth modes, but both modes becoming asymptotically the same at high death rates, with cells dividing uniformly throughout the tumour volume. Specifically, the results shown in Fig. 2B of the main text and S6 of a flat distribution of direction angles (or direction cosines in the 3D case) show that there is no enhanced cell growth near the edge of the tumour. However, it is possible that surface growth would manifest itself in a tumour that grows more slowly.

#### S3.1 A model of surface growth with explicit spatial dependence

To probe the distinction of surface growth and volume growth at different rates of cell death, we briefly discuss an alternative model of surface growth. Instead of spatial constraints leading to a density-dependent growth rate (see Section S1), we make the rate of cell division explicitly depend on the spatial position of each cell. For concreteness, we consider a division rate that is zero in the tumour centre and increases with the radial distance $r$ from the tumour centre to $b = 1.$ at its edge,

$$b(r) = \frac{b}{1 + \exp\left(-\frac{r-(R-w/2)}{w/s}\right)} \, . \tag{S.2}$$

$R$ denotes the radius of the tumour, which changes during the simulation, while the parameters edge division rate $b = 1$, edge width $w = 15$ and the profile steepness $s = 10$ are initially set. Under this model, cells in the bulk cease to divide as the edge proceeds to grow outwards. Consequently, the mutation density in the bulk does not increase over time. Figure S10 shows that under the model where the cell division rate depends on the local density, the surface growth mode has a flat mutation density curve in the high death rate limit because the high turnover allows cells in the tumour bulk to divide at the rate of cell death. In contrast, the mutation density profiles of simulations with explicit spatial dependence of the division rate $b$ on the radial position cells collapse to a line with positive slope when rescaled by the effective mutation rate $\mu/(1-d/b)$. The dependence of the scale on the turnover rate $d/b$ is simply given by the number of generations $T$ to reach size $N$ with $T = (1-d/b)\log(N)$.

On the other hand, when cells at the center cannot divide even at low density, any positive cell death rate $d > 0.$ leads to a regions without cells (mimicking a necrotic core), which gets larger with higher turnover rate $d/b$.

Simulations of this model of explicit spatial dependence produce the same signature of surface growth in their distribution of directional angles as found in Section S2 for the model of surface growth.

#### S3.2 Mutation density in the Ling *et al.* data

Figure S11 shows the mutation density versus the distance from the tumour centre for the 285 samples from Ling *et al.* which have been genotyped. The resulting curve is compatible with a flat line, corresponding to a growth mode where new mutations arise uniformly throughout the volume.

Throughout this analysis, we focused on the mutation *density*, rather than the number of mutations found in a ring. The reason is that the area of a ring of fixed width increases linearly with the radius $r$ of the ring (for a small width). For this reason, under volume growth, the *number of mutations* in a ring increases linearly with its radius.

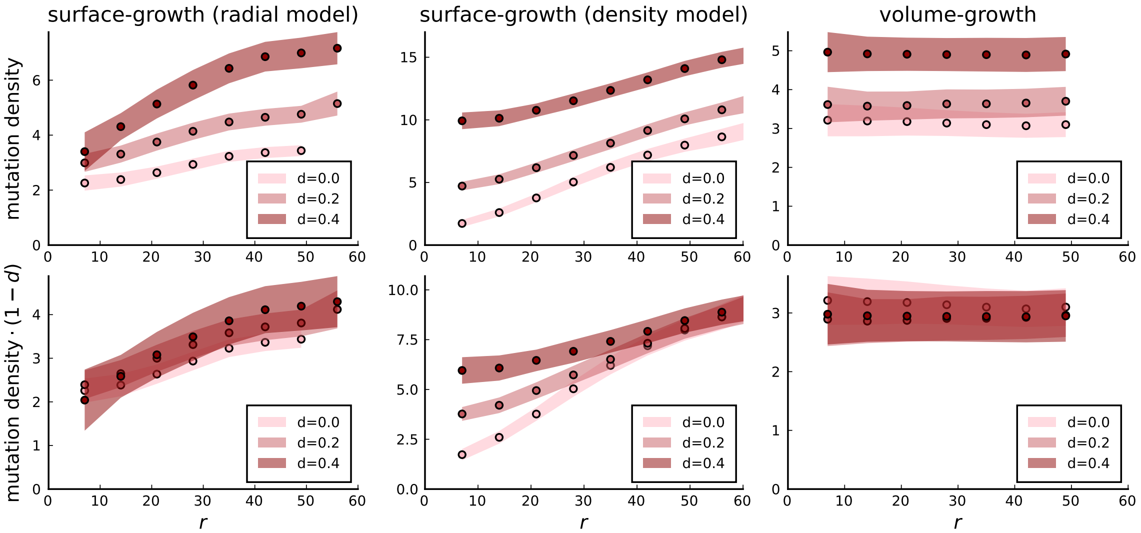

**Figure S10: Mutation density profiles under different growth models.** The mutation density is shown as a function of the ring radius for the model of position-dependent surface growth (with a division rate (S.2)nearly zero below the surface), for the density-dependent surface growth model, and under volume growth. Rescaling mutation densities by $(1 - d/b)$ collapses the radial model and the volume growth model to a single curve. Settings for 3d simulations under all models are as in S9: $N = 40000$, $b = 1.$, $\mu = 0.3$.

#### S3.3 Mutation density on shells in 3d

Extending the measurement of the radial change in mutation density to samples taken in a 3d spherical tumour requires only a minor adjustment to the definition as presented in the beginning of section S3. Instead of a ring of radius $r$ we now consider a spherical shell of width $w$ containing the samples with a radial distance within $r$ and $r + w$ from the tumour centre. The mutation density is the ratio between total count of mutations summed across samples and the number of samples in a given shell.

Figure S9 shows the mutation density on rings as a function of the radius for single cells of planar cuts through simulated 3d spherical tumours. The corresponding curves for single cells on spherical shells at different radii of the simulated tumours are virtually identical to the mutation density in rings of the cross section. As expected, the density of mutations is independent of the radius for volume growth and increases with radius under surface growth.

However, imitating the sampling scheme of Li *et al.* in simulations, as explained in Section S2.2, shows a poorer distinction between the two modes of growth, see figures S12 and S13. The radial increase in mutation density under surface growth is dampened in particular at the tumour surface. Under volume growth the curve is still flat (within the error margins). The mutation density calculated for tumours T1 and T2 has a considerably larger decrease near the surface. The slight decrease at high $r$ seen both in the empirical data and the simulations might be caused by the uneven sampling for whole-genome sequencing: WGS samples are taken from few slices mostly inside the tumour and might miss mutations that appear near the surface.

We note here that these results appear to contradict findings in Li *et al.* [8] concerning the distribution of mutations across the tumour. Specifically, Li et al. state that ”peripheral regions not only accumulated more mutations, but also contained more changes in genes related to cell proliferation and cell cycle function” and ”Phylogenetic trees show that branch lengths vary greatly with the long-branched subclones tending to occur in peripheral regions”. We find that the classification of samples into ‘centre’ and ‘periphery’ by Li et al. does not coincide with the distance of samples from the centre of the tumour, see Figure SI S14. For tumour T1, there are several samples labeled as ‘centre’ are further from the centre as some samples classified as ‘periphery’, for example L1. Furthermore, the samples labeled ‘centre’ have mutations that belong to a single clade (see Fig. 5 in [10]), which suggest that the classification scheme mixes spatial and genomic information.

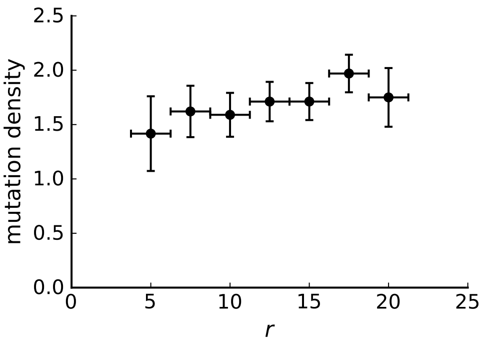

**Figure S11: Mutation density curve of genotyped samples** The 285 samples from Ling *et al.* that have been genotyped have a mean nearest neighbour distance of about 1.5mm. Choosing a ring width of 2.5mm allows us to define 8 rings around the samples' centre of mass. The horizontal error bars indicate that samples can fall anywhere in this 2.5mm window. Vertical error bars were estimated using the standard deviation of the mutation density under volume growth derived in the text $\mathrm{std}(\rho_r) = \sqrt{\langle\rho_r\rangle / N_r}$, where we use the measured value of $\rho_r$ in place of $\langle\rho_r\rangle$ to estimate the error. We found in numerical simulations that this yields a lower bound on the error in 3d simulations with sampling. The curve can be considered flat if the change in value is of the same scale as the error. Our null model is volume growth, therefore, a lower-bound of the error estimate favors rejecting flatness of the curve (the signature of volume growth), thus making the estimate conservative. The flat shape of the mutation density curve is again consistent with volume growth rather than surface growth. Rings at radius 50 (centre) and 450 (edge) contain only 2 samples each and were thus dropped.

#### S3.4 SDevo analysis on simulated volume growth

In this section, we compare our finding of volume growth in the Li *et al.* data [8] to the recent result of Lewinsohn *et al.* [10]. In a phylogenetic analysis of the Li *et al.* data [8], Lewinsohn *et al.* found a significantly elevated rate of cell birth (relative to death) near the edge of the tumour, compared to the tumour's core. This is at variance with our result on the same data, which showed no signal of the outward growth associated with surface growth. We show that the result of Lewinsohn *et al.* [10] is compatible with volume growth: we demonstrate that the phylogenetic analysis of [10] produces a signal of surface growth also on artificial data from a volume growth model, when samples are placed precisely as in the empirical data by Li *et al.* and analysed as in [10].

We grew a population from a single cell to size 40000 in 3d at a relative rate of cell death $q = 0$ and on average $\mu = 10$ mutations per generation. The growth mode was volume growth as described in SI Section S1. Repeating the analysis at $q = .2, .4, .6, .8$ gave similar results. Samples of typically 30 cells were taken from the same positions they were taken from in tumours T1 and T2 of [8], respectively. Samples were classified following the classification used in Extended Data Fig. 7 of Lewinsohn *et al.*. In this scheme, the samples with distance from the centre higher than 0.9 of the tumour's radius are classified as 'edge samples', those with distance less than this threshold are classified as 'core samples'. This classification differs from that that used in the main text of Lewinsohn *et al.*, which uses the classification introduced in Li *et al.* [8] briefly discussed in this SI Section S3.3. As the latter classification is not based on distance alone, we will not pursue it here.

We then applied the algorithm SDevo developed in [10] with the parameters provided in xml-template-files by Lewinsohn *et al.* on the Github page `https://github.com/blab/spatial-tumor-phylodynamics`.

Figure S15 shows the resulting distribution of inferred birth rates in the edge and core parts of the tumours across 30 different simulated tumours grown under volume growth. The mean ratio of the inferred birthrates in the edge versus centre samples was 11.47 (95% interval $[0.03, 49.70]$), even though for volume growth this ratio should be one, or close to one. Specifically, this ratio is higher, and hence the spurious signal for surface growth is stronger, than what Lewinsohn *et al.* found in the empirical data: Extended Data Fig. 7 of [10] reports the estimated birth rate ratio (edge/core) of 1.15 for tumour T1.

For samples placed as in tumour T2, we found a mean growth rate ratio of 6.98 (95% interval $[0.07, 33.12]$),

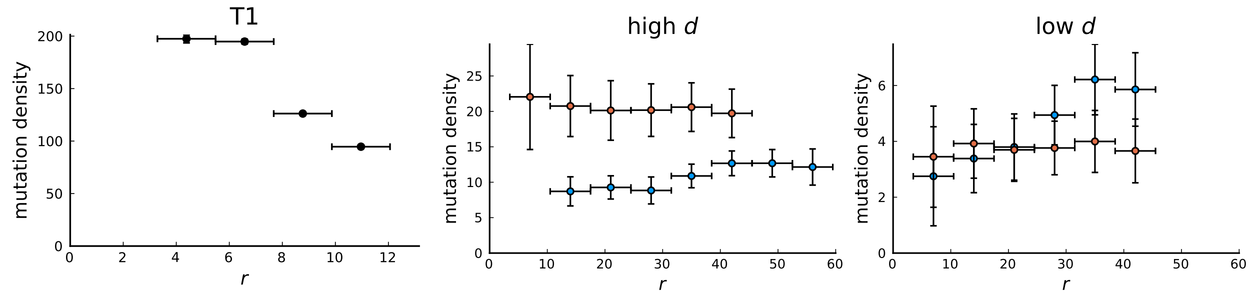

**Figure S12: Mutation density curve of tumour T1 (left) and T1-like sampling in simulations (centre and right).** The plots show mutation density for spherical shells at different distances $r$ from the tumour center. The left plot is based on the 153 genotyped samples from 9 slices of T1 that report the presence or absence of mutations found by whole-genome sequencing of 16 samples. The other two plots show mutation density as a function of shell radius $r$ in simulations of 3d spherical tumours under T1-like sampling. Within each simulation 16 samples are taken from 3 slices for sequencing and 153 samples from 9 slices to check for the presence of the detected mutations. The positions of slices and samples are specified by T1. Parameters of the 3d simulations are as in Fig. S8.

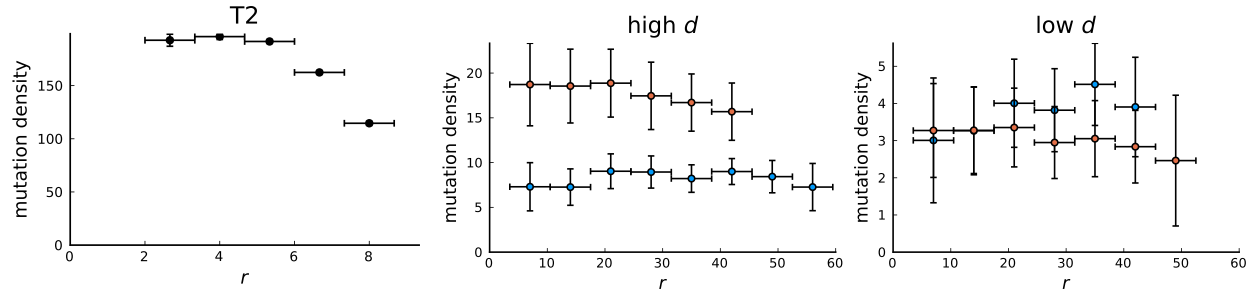

**Figure S13: Mutation density curve of tumour T2 (left) and T2-like sampling in simulations (centre and right).** The plots show mutation density for spherical shells at different distances $r$ from the tumour center. The left plot is based on the 151 genotyped samples from 6 slices of T2 that report the presence or absence of mutations found by whole-genome sequencing of 9 samples. The two plots for simulations (middle and right) are analogous to Fig. S12 but with T2-like sampling.

compared to 3.89 reported for tumour T2 in Extended Data Fig. 7 of [10]. From Figure S15 we conclude that in many cases of artificial tumours grown under volume growth, SDevo detects a higher birth rate in samples near the edge of the tumour, and therefore does not reliably differentiate between surface and volume growth.

A potential cause for this result is the sampling probabilities. The inference of birth-death model parameters from a phylogenetic tree generally depends on estimates of the probability that an individual from a (possibly large) population was sampled [11]. Correspondingly, SDevo requires estimates of the sampling probabilities in the core/edge regions of the tumour as input parameters. From the xml files referenced above we found these parameters were set to 0.2 and 0.1 for the core and edge, respectively. We estimated the sampling probabilities by measuring tumour volume in units of closely packed spheres of the size of samples. Specifically, we calculated the number of samples in a particular part of the tumour (core or edge) divided by the number of spheres of the size of samples that fit into the corresponding volume (edge or core region of a particular tumour) under close packing. This gave $f_{\text{core}} = .00006$ and $f_{\text{edge}} = .00012$ for T1 and $f_{\text{core}} = .00004$ and $f_{\text{edge}} = .00024$ for T2, much lower than the sampling probabilities used in [10]. The absolute sampling probabilities (not just the relative one between edge and core) can in principle affect the inference results, because the inferred death rate depends non-homogeneously on the sampling probability [11], and birth and death rate are inferred relative to one another. However, there is a difference even in the relative sampling probability of core and edge regions between our estimates and the parameters used in [10]; the ratio of sampling probabilities is 2 (core relative to edge) in [10] and about 1/2 in our estimate for T1 and 1/6 for

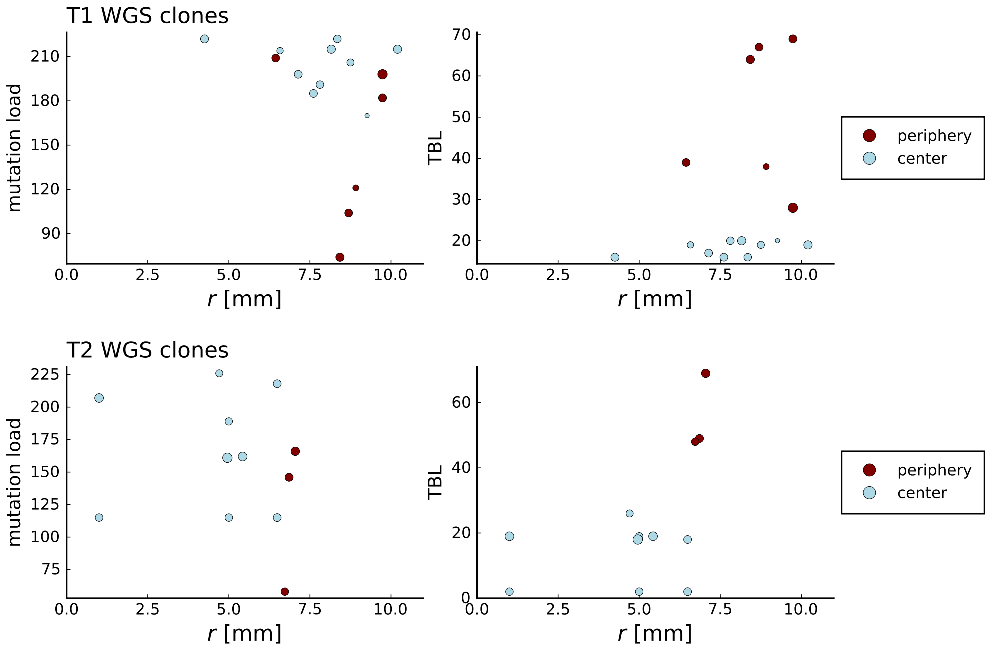

**Figure S14: Clone sizes and terminal branch lengths (TBL) for genotyped mutations in Li et al. WGS samples.** The plots on the left (right) show the number of mutations (terminal branch length) of clones in the WGS samples of tumour T1 (top) and T2 (bottom) against the radial position relative to the tumour center of mass in mm on the x-axis. The size of markers indicates abundance of the clone within the sample and colors are determined by the classification of samples by Li et al. into "periphery" and "center". Samples marked as peripheral have higher terminal branch lengths but in the case of T1 are not located closer to the tumour boundary than those marked as central.

T2.

To test this hypothesis by running SDevo with the sampling probabilities as estimated here, however, this not not lead to qualitatively different results. Further work would be needed to determine the capabilities and the limitations of the SDevo algorithm [10].

### S4 Spatial dispersion of cells

We quantify how clones are dispersed throughout the tumour, using the WE-sequencing data of [7] and [8]. Rather than looking at sum of spatial distances between cells with a particular mutation, we define a dispersion parameter as these sum of distances relative to those in a dense packing of cells. The dispersion parameter is one for a (hypothetical) dense and spherical arrangement of cells carrying a particular mutation, and larger than one if these cells are dispersed within the tumour (with cells not carrying that mutation between them). For each mutation $m$ we compute the dispersion parameter $\sigma_m$ as follows:

#### S4.1 Algorithmic description

1. Obtain all mutant clades and compute whole-tumour frequencies
2. Discard mutations that occur in a single sample or fall below a resolution threshold $f_{\text{res}}$
3. Estimate the sample density $\rho_s$ as the number of samples devided by the sampled surface area
4. For each mutant clade $m$ do
   (a) obtain the frequencies (cancer cell fractions) $f_i^m$ of mutation $m$ in all samples $i$ of the clade

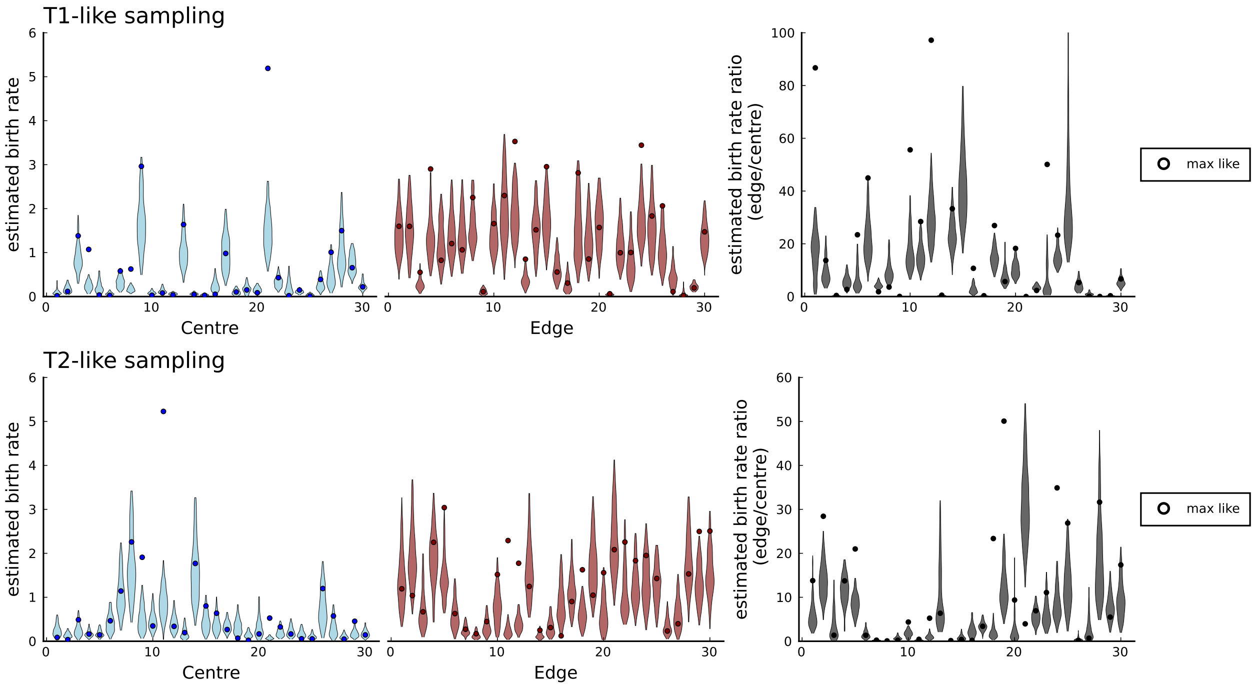

**Figure S15: Birth rates for periphery and center inferred with SDevo from simulations of volume growth with spatial sampling.** We run 30 simulations in 3d to a population size $N = 40000$ at rates $b = 1.$, $d = 0.$, $\mu = 10.$ and take punch samples for deep sequencing at positions specified in [8] for tumours T1 and T2. We only call mutations exceeding a 0.3 cellular fraction threshold within a sample as specified in [10] and use the template xml file provided by Lewinsohn *et al.* to generate inputs for the SDevo algorithm. Each simulation shows a violin plot of inferred birth rates (95% interval of last 500 sampled states of the MCMC-chain in SDevo) for samples labeled as 'centre', samples labeled as 'edge', and the ratios of the two rates, as in Fig. 5 of [10]. A circle marker for each violin plot indicates the values at the maximum likelihood, which often falls outside the 95% interval of the last 500 sampled states.

(b) compute the dense packing radius $r_m = \sqrt{\sum_i f_m^i/(\pi\rho_s)}$

(c) compute the dense packing dispersion [1]

$$\sigma_m^{\text{dense}} = 128/(45\pi) r_m \tag{S.3}$$

(d) compute pairwise distances $d_{ij} = |\vec{p_i} - \vec{p_j}|$ and weights $w_{ij}^m = f_i^m f_j^m$

(e) compute the relative dispersion $\sigma_m = (\sum_{j,i<j} w_{ij}^m d_{ij} \;/\; \sum_{j,i<j} w_{ij}^m) \;/\; \sigma_m^{\text{dense}}$

5. plot the (unweighted) histogram of the values $\sigma_m$

[1] To compute the mean distance of points on a disk, we consider two points on the disk with distance $r$ (in units of the disk radius $R$) at an angle $\theta$. We note that the probability density $\delta(r)$ is proportional to the area of the disk of radius $R$ with points that have a distance $r$ to another point on the disk at angle $\theta$. This area is given by the intersect of the disk to the disk shifted by $r$ in direction $\theta$. This intersect is segmented by the shift-axis and the orthogonal axis into four identical circle segments of area $4\int_{r/2}^{1}\sqrt{1-x^2}dx$. Considering distance vectors for all $\theta$, the density function $\delta(r)$ is proportional to an additional factor $r$ for the circumference of $r$-vectors. Normalizing by integration from $r = 0$ to 2 yields

$$\delta(r) = \frac{r}{\pi}\left((4\arctan\left(\frac{\sqrt{4-r^2}}{r}\right) - r\sqrt{4-r^2}\right).$$

The mean value of $r$ can then be computed as

$$\langle r\rangle = \int_0^2 r\delta(r)dr = \frac{128}{45\pi}.$$

Frequencies of mutations within single samples and within the whole tumour are crucial to all quantitative analysis in this work. What we call mutation frequencies are cancer cell fractions (CCF) of mutations in sequencing data which estimate the fraction of cells carrying that mutation. Given a sample with variant allele frequency $f_{\rm vaf}$ (VAF), sample purity $\rho$ and ploidy $p$ we get the mutation's multiplicity $m$ by rounding $u = f_{\rm vaf} \cdot (p \cdot \rho + 2 \cdot (1-\rho))/\rho$ to the nearest (nonzero) integer and the cancer cell fraction $f = u/m$. The whole-tumour frequency of a mutation is its tumour wide cancer cell fraction, computed as the average over samples and weighted by the sample purity (assuming equal sample sizes) $\bar{f} = \sum_i f_i \cdot \rho_i / \sum_i \rho_i$ [12].

To put the algorithm in words, in (1-2) we consider clades of mutations above a whole-tumour frequency cutoff $f_{\rm res}$. For a given clade defined by mutation $m$, we measure the spatial distance $d_{ij}$, in step 4d between pairs of samples i,j that contain $m$ and assign a weight that estimates the number of single cell pairs that carry $m$ between both samples. The absolute dispersion of mutation $m$ is defined as the weighted average over these pairwise distances. But different mutations have different whole-tumour frequencies and would naturally take up areas of different sizes. We therefore measure dispersion relative to the value of dispersion a clade would have if all cells were densely packed in a disk, as given by step 4e. The dispersion for dense packing can be computed as the mean distance between all points on a disk of radius $r_m$ by equation (S.3) in step 4c.

As in the algorithm for distinguishing surface from volume growth (see Section S2), every mutation contributes with equal weight to the computed distribution, here the distribution of dispersion $\sigma_m$. Summary statistics of the dispersion parameter $\sigma_m$ such as the mean over mutations, or its distribution over mutations show the extent of dispersion and cell migration within the population. To compare dispersion distributions observed in empirical data to those seen in spatial simulations under comparable sampling in the surface and volume growth regimes, we use the (approximate) two-sample Kolmogorov-Smirnov test between the experimental and the simulated distribution. The Kolmogorov-Smirnov (KS) statistic is the maximum distance between the two cumulative distribution functions (cdf) $F$, $D_{n,m} = \sup_\sigma |F_{1,n} - F_{2,m}|$ with the number samples $n$,$m$ of the two empirical distributions respectively. $D$ is a measure of how distant the two distributions are from one another and can be used two rank pairs of distributions by similarity. When normalized by the factor $\sqrt{nm/(n+m)}$ and given a rejection level $\alpha$ (here we use the default $\alpha = 0.05$), a p-value can be computed from the statistic $D_{n,m}$ which states how likely it is that the empirical distributions are sampled from the same underlying distribution - that is, a low p-value indicates that the two underlying distributions are significantly different.

The choice of mean pairwise distance as a measure of dispersion is conceptually simple, but other choices are also possible. Similar measures such as the radius of gyration (root mean square radial displacement) $\sigma_m^{\rm gyr} = \sum_i f_i^m |\vec{p}_i - \langle \vec{p} \rangle_m|^2 / \sum_i f_i^m$ (around the clade center of mass $\langle \vec{p} \rangle_m = \sum_i f_i^m \vec{p}_i / \sum_i f_i^m$ ) yield the same qualitative results and their dispersion under dense packing can readily be calculated as an expectation value over the disk.

#### S4.2 Processing of whole-exome sequencing data

Fastq files of the sequences were downloaded from `https://bigd.big.ac.cn/search/?dbId=gsa&q=PRJCA000091&page=2` [7]. Fastq files were processed using our in-house mapping pipeline based on GATK4 (version 4.1.0.0 from DockerHUB). We used GRCh37.p13 as the reference genome and gencode.v19 as the annotation file. Singletons, and reads with less than 70 bp were discarded. Additionally, we excluded known variants in the human population (taken from https://data.broadinstitute.org and ftp://ftp.ncbi.nih.gov). We used the interval bed file S02972011 from SureSelect all human exon version 3. Variant calling and initial filtering was performed with Mutect2 (GATK), using the joint variant caller mode and applying the optional filter.

We focus on the high-depth WES data instead of the set of genotyped samples for two reasons: Firstly, because dispersion, unlike the directional growth bias and the turnover, is based on individual mutations alone, we do not need to infer clones or parent-offspring relationships and can effectively leverage the sample frequencies from whole-exome sequencing. Secondly, the distribution of dispersion values turns out to be very noisy for the set of 35 mutations probed by genotyping in the high-spatial-resolution data. However, we have measured the dispersion parameter for this small set of mutations both in the WES samples and the genotyped samples and found that their respective dispersion distribution shows the same large tail as the dispersion distribution of the 217 subclonal SNVs detected by Mutect (see main text, Fig.3).

To compute the dispersion parameter for the WE-sequencing data of Ling *et al.*, we call mutations using

Mutect 2 which jointly calls on all 23 samples. We obtain 238 subclonal SNVs by imposing the following filters: a given SNV, 1) passes Mutect's filters, 2) has no supporting ALT reads in the normal sample, 3) has 5 or more supporting reads in total, 4) has a total of 150 or more reads (coverage), and 5) has a whole-tumour frequency (CCF) $\bar{f}_m$ between 0.025 and 0.3 (subclonal). The criteria 2)-4) are adopted from the variant calling by Ling *et al.* 2) and 3) eliminate false positives, 4) assures accuracy in the estimation of mutant frequencies while at the same time also reducing the effective genome size (see main text, "Cell turnover"). Setting a lower bound on the number of supporting alternate reads, filter 3) determines the resolution of mutant frequencies.

Regarding the cutoffs on whole-tumour frequencies 5), the CCF-sprectrum consists of two clearly distinct clusters: Clonal mutations forming a large gaussian peak near $\bar{f}_m = 1$ and subclonal neutral mutations at low frequencies following a power-law decay, with very few mutations falling around $\bar{f}_m = 0.5$ inbetween the two profiles. We found that imposing an upper cutoff at $\bar{f}_m = 0.3$ fully removes clonal mutations while recovering most subclonal mutations. The distribution of $\sigma$ turns out to depend on the minimum whole-tumour frequency of mutations, therefore, we also need to set a cutoff on $\bar{f}_m$ at the lower end of the SFS CCF-spectrum in order to compare with simulations. This dependence is linked to the size of the tumour when specific mutations arose, see S7. The distribution of $\sigma$ turns out to depend on the minimum whole-tumour frequency of mutations, therefore, we need to set a cutoff on $\bar{f}_m$ at the lower end of the SFS in order to compare with simulations. This dependence is linked to the size of the tumour when specific mutations arose, see S7.

We choose a minimum whole-tumour frequency of $f_{\text{res}} = 1/40$, because this is the frequency in the SFS below which the number of detected mutations drops rapidly, as can be seen from the saturation in the cumulative SFS (see Fig. S21). The cumulative SFS counts the number of mutations with frequency larger or equal to $f$ as a function of $1/f$. It is monotonically increasing and would extend to the smallest mutation frequency but effectively flattens below the sequencing resolution $f_{\text{res}}$. Beyond $f_{\text{res}} = 1/40$ the sequencing disproportionately misses low frequency mutations which would be relevant to the comparison with simulations at cutoffs lower than $f_{\text{res}}$ for the dispersion parameter.

#### S4.3 Dispersion in two dimensions

We compare the dispersion parameter in the data of Ling *et al.* to simulations of populations growth to 40000 cells under volume growth ($\rho = \infty$) and surface growth ($\rho = 6$) which we sample to mimic the spatial sampling of [7] as described in Section S1.2. For each simulated 3d tumour, we take a planar cross-section through the centre of mass and cover the plane with a triangular lattice that fits 23 samples of 20 cells each at its nodes. To mimic stochastic effects in sequencing, in each sample and for each mutant we draw a binomially distributed number of alternate reads given the true frequency and a read depth of 20 reads. These choices follow from the sequencing depth in the WES data of Ling *et al.*, which has a mean coverage of 20 reads per sample. In the simulations, samples consist of 20 cells, which allows for a sequencing depth that is sufficiently high to resolve whole-tumour mutation frequencies even below 1/40. In fact, given the limited sequencing depth we found that increasing the sample size has no effect on the distribution of dispersion parameter.

We compute whole-tumour frequencies $\bar{f}_m$, retain mutants with $\bar{f}_m > 1/40$ - the cutoff used for mutations in the WES data of Ling *et al.* and determine their dispersion distribution as described above.

The normalized Kolmogorov-Smirnov statistic and log-p-value between the data and simulations in 3D with sampling (Fig. S18) are 5.742 and $-65.3$ for surface growth and 1.13 and $-1.9$ for volume growth, confirming that the volume growth regime fits the observed distribution of dispersion parameters much better than the surface growth regime. (A perfect fit, leading to a KS statistics of zero, is not expected, for instance because of fluctuations due to the finite number of mutations.)

The same methods apply for uniformly sampled single cells ($f_m^i = 1$), see Fig. S17, and for the sampling scheme used for Fig. S16. However, simulations show that the absolute scale of dispersion $\sigma_m$ strongly depends on sampling density, namely, the value of mean dispersion (in the volume growth limit) increases as the sampling density gets reduced. It is therefore import to reproduce the sampling settings when comparing simulations to sequencing data.

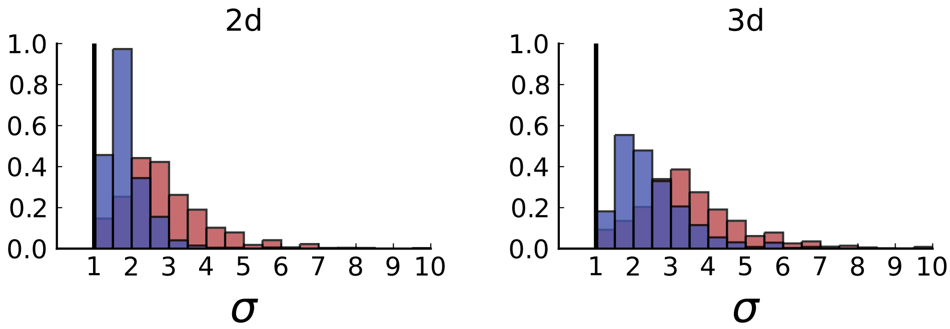

**Figure S16: Dispersion distributions in simulations including spatial sampling - 2d (left) and 3d (right).** Histograms of the dispersion parameters under spatial sampling with 23 samples with a sequencing model mimicking the whole-exome sequencing of Ling *et al.* Samples are taken from populations grown to 10000 cells in 2d (left) and cross-sections of 3d simulations of populations grown up to 40000 cells (right). As in Section S2, we use the cell division rate $b = 1$, mutation rate $\mu = 0.3$, as well as cell death rates $d = 0.8$ and $\rho_c = 6$ for volume growth (red) and $d = 0.4$ and $\rho_c = \infty$ for surface growth (blue), respectively (see also Fig. S4).

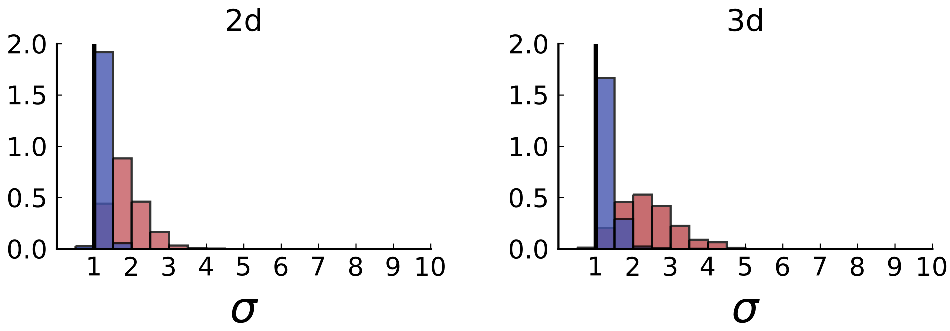

**Figure S17: Dispersion distributions in simulations with sampling of single cells - 2d (left) and 3d (right).** Same as in S3 but for single cells under volume (red) and surface growth (blue) at the same parameters $b, d, \mu$.

##### S4.4 Dispersion in three dimensions

Just like the measure of the direction of mutants in S2.2, the dispersion can also be computed for samples taken in three dimensions. Only the mean distances of cells in a densely packed sphere $\sigma_m^{\text{dense}}$ is different from that of the two-dimensional disk used in steps 4(b) and 4(c): Given a 3d sample density $\rho_s$, a mutant occurring with frequencies $\{f_m^i\}$ in samples $\{s_i\}$ could be packed into a sphere of radius $r_m = \left(\sum_i f_m^i/(4/3\pi)\right)^{1/3}$. The mean distance of points within this sphere is $\sigma_m^{\text{dense}} = 36/35 r_m$. [2]

Based on the corresponding definition of the dispersion parameter, we measure the dispersion of mutants

[2] To see this, consider two points $\mathbf{x}$ and $\mathbf{y}$ within the sphere of radius $R$ on shells of radius $r_1 < R$ and $r_2 < R$ and with an angle $\theta = \angle(\mathbf{x}, \mathbf{y})$. The distance between the two points is $\sqrt{r_1^2 + r_2^2 - 2r_1 r_2 cos(\theta)}$, so the average distance for points within the sphere becomes

$$\langle|\mathbf{x} - \mathbf{y}|\rangle = \int_0^R f_{r_1}(r_1') \int_0^R f_{r_2}(r_2') \int_0^\pi f_\theta(\theta') \sqrt{r_1'^2 + r_2'^2 - 2r_1' r_2' cos(\theta')} dr_1' dr_2' d\theta',$$

where the density functions $f_{r_1}(r') = 3r'^2/R^3$, $f_{r_2}(r') = 3r'^2/R^3$ are proportional to the surface area of the sphere with radius $r'$ and the density function $f_\theta(\theta') = \sin(\theta)/2$ is proportional to the circumference of the circle defined as the rim of the cone

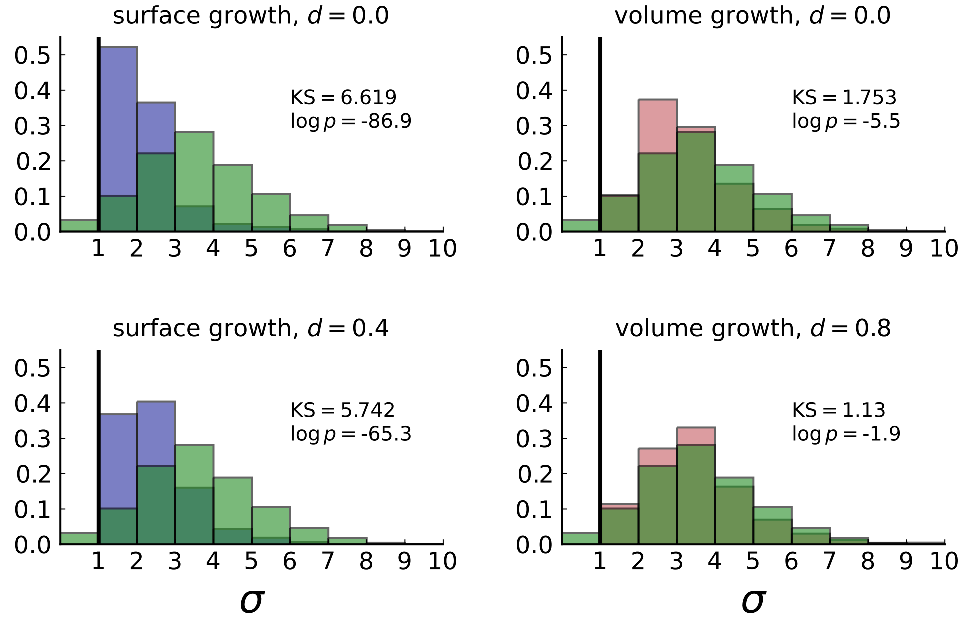

**Figure S18: Normalized Kolmogorov-Smirnov distance between dispersion distributions of Ling *et al.* data and simulations.** Each frame compares the histogram of the dispersion values $\sigma$ of subclonal mutations in the WES data (in green) to that of simulations in 3d with spatial sampling under volume (red) and surface (blue) growth at low and high turnover rates $d$. Simulation settings are as in S16, right. The KS statistic serves as a measure of distance between the two distributions with smaller values indicating that the data is more likely to match the model. The dispersion observed in the data fits with simulations of volume growth at a high rate of turnover (KS = 1.13, bottom right).

in the whole-genome sequencing (WGS) data of Li *et al.* The data consists of 16 samples from tumour T1 and 9 samples from tumour T2. The cumulative SFS reaches a plateau at frequency $f_{\rm res} = 1/30$ for T1 and $1/18$ for T2, which is the respective whole-tumour frequency resolution, as discussed for the Ling *et al* in the previous Section S4.2. The frequency resolution is lower than in Ling *et al.*, in agreement with the expected linear scaling between the inverse frequency resolution and the number of samples $n$ which can be written as $(nf_{\rm res})_{\rm Li} = (nf_{\rm res})_{\rm Ling}$ given that both studies sequenced at almost the same read depth per sample.

Whereas in the case of the tumour data of Ling *et al.*, samples were taken nearly spatially uniformly for subsequent deep sequencing analysis, in the two tumours T1 and T2 only few of the slices were used for WGS. To account for this uneven sampling we use the same sample positions in simulated 3d tumours: For each simulation we take slices at specified heights and sample each slice at the positions of the WGS samples from tumours T1 and T2. The resulting distributions of dispersion values for mutants for T1 and the corresponding numerical simulations are shown in Fig. S19. The dispersion values observed for tumour T1 are high compared surface growth simulations (as in the 2d data by Ling at al.), and the KS statistics is not compatible with surface growth, see Fig. S19.

In the case of T2, there are insufficient mutations that appear in a sufficiently large number of samples. Out of 150 mutations in T2, only 2 appear in more than 4 samples. (Out of 308 mutations in T1, 58 appear

with side length $\max(r_1, r_2)$ and opening angle $\theta'$. The inner integral yields

$$\int_0^{\pi} d\theta' \sin(\theta')/2\sqrt{r_1^2 + r_2^2 - 2r_1r_2cos(\theta)} = \frac{1}{6r_1r_2}((r_1+r_2)^3 - |r_1 - r_2|^3),$$

and the outer integrals can be computed by distinguishing the two cases $r_1 < r_2$ and $r_2 < r_1$ resulting in

$$\langle |\mathbf{x} - \mathbf{y}| \rangle = \frac{36}{35} R.$$

in more than 4 samples.) The dispersion parameter becomes unreliable when the number of samples involved is low; a few samples from a spherical distribution of mutants typically yield artefactual values of $\sigma$ higher than one, due to the sparse and uneven distribution of samples in space.

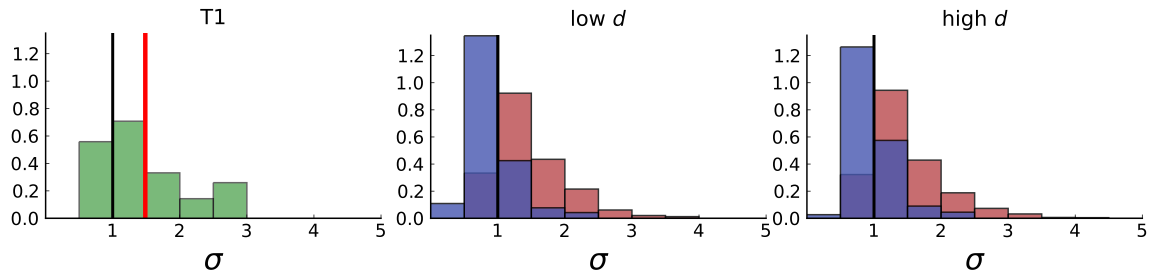

**Figure S19: Dispersion distribution of tumour T1 and T1-like sampling in simulations.** The left plot shows dispersion values of 308 mutations with frequencies larger than the frequency resolution $(f_{\mathrm{res}})_{\mathrm{T1}} = 1/30$ based on the 16 WGS samples of T1. The plots on the right show dispersion in simulations of 3d spherical tumours under T1-like sampling, meaning that, from each simulation 16 samples are taken from 3 slices, where the positions of slices and samples are specified by T1. Parameters of the 3d simulations are as in Fig. S16. The normalized KS statistic and log-p-value between the data and simulations are 7.272 / 7.267 and $-105.1$ / $-104.9$ for surface growth (low/high $d$) and 1.652 / 2.189 and $-4.8$ / $-8.9$ for volume growth (low/high $d$).

### S5 Cell turnover

We infer the cell death rate and mutation rate by tracking the dynamics of clades and clones. A clone describes the set of cells with a particular genotype. In the absence of back mutations, a clade describes the set of cells carrying a particular mutation. (With back mutations, a clade is the set of descendants of a particular mutant cell, including the original mutant itself.) Under (volume) exponential growth, a clade is expected to grow exponentially at the growth rate $\lambda = b - d$ and maintain on average a constant frequency within the tumour [13]. A clone grows in a similar way, but at a rate which is reduced to mutations which generate new clones.

However, clone and clade sizes are also subject to fluctuations due to genetic drift: Clones and clades can become extinct even in growing populations, and the rate at which that happens depends on the rate of cell death (relative to the rate of birth) and the mutation rate. We use the fraction of clones with extinct parental clones and the fraction of offspring clades which coincide with a given ancestral clade to infer both the mutation rate and the relative rate of cell death.

Any mutation we encounter when tracing back the lineage of an offspring clade constitutes an ancestral clade. A clade coincides with an ancestral clade if all clones within the ancestral clade are also part of the offspring clade (a clone carrying mutations $m_i$ is part of every clade $i$ defined by mutation $m_i$). Under an infinite site model, this situation refers to a pair of coinciding mutations $m_1, m_2$, i.e. any cell carrying $m_1$ also carries $m_2$ and vice versa - $m_1$ and $m_2$ are the branching points within the lineage from which ancestor and offspring clade originated, respectively. A clone with extinct parental clone necessarily implies a clade that coincides with its most recent ancestral clade. Clades however can coincide with several ancestral clades if these underwent successive sweeps and ended in the fixation of the offspring clade. So more generally, for $n-1$ ancestral clades coinciding with the offspring lineage there is a set ${m_i}_{i=1}^n$ of mutations that is common to all cells carrying any of the mutations $m_i$.

We begin with a small but illustrative example of clone and clade turnover in Section S5.1 before introducing the turnover based inference scheme in Section S5.2. We then test this scheme on simulated data (also under spatial sampling) in Section S5.3. For a derivation of analytical expressions and a discussion of the inference model used in this section, see Section S5.4 below and [14].

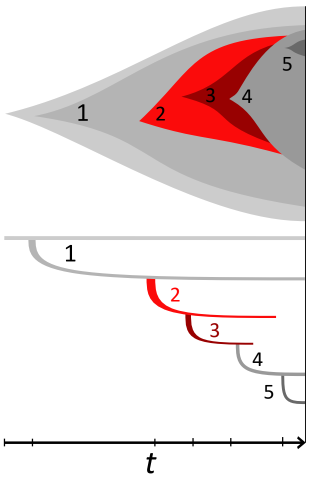

**Figure S20: Example phylogeny and turnover.** This illustration shows a growing population of cells that accumulates 5 mutations, on the top showing the muller plot of the 5 clones' sizes and below the corresponding phylogeny against time $t$. The growth rate is positive but under stochastic dynamics clones and clades may become extinct. The clones carrying mutations $\{1,2\}$ and $\{1,2,3\}$ went extinct and their offspring clones/clades add to the clone/clade turnover, see text. Specifically, the clone $\{1,2,3,4\}$ lost its parental clone, similarly the clade defined by mutation 3 coincides with clade 2 and clade 4 in turn replaced both clades 2 and 3.

#### S5.1 Clone and clade turnover example

We consider a population in which mutations $1,2,3,4,5$ arose in that order (the order is given only to illustrate the example, see S20). In this population, clones carrying mutations $\{\}$, $\{1\}$, $\{1,2,3,4\}$, $\{1,2,3,4,5\}$ survived. Clone $\{1\}$ has an extant parental genotype, namely $\{\}$. None of the extant genotypes carry exactly three mutations out of $\{1,2,3,4\}$. The parental clone of this genotype thus became extinct. Finally genotype $\{1,2,3,4,5\}$ has an extant parent in $\{1,2,3,4\}$.

Hence 1 out of 3 clones have extinct parents, resulting in a clone turnover of $W_c = 1/3$. In order to compute the clade turnover we go over all mutations $1,2,3,4,5$, which each define a clade. The only ancestral clade of mutant 1 is the original clade which gave birth to all clades. The clone $\{\}$ survived, therefore clade 1 does not coincide with its ancestor. Mutation 1 hence contributes one to the denominator and zero to the numerator of the clade turnover. Mutation 2 does not coincide with either of its two ancestral clades (origin and 1) and thus adds two to the denominator and zero to the numerator.

The clade of mutation 3 is necessarily also distinct from the ancestral clades of mutation 2 - however, its clade has replaced that of 2 and therefore adds three to the denominator and one to the numerator. Note that the clone $\{1,2,3\}$ does not need to be extant; as long as clade 3, which includes its subclades 4 and 5, replaces clade 2 it adds to the turnover. The clade of 4 in turn coincides with both 2 and 3, resulting in a contribution of four to the denominator and two to the numerator. Lastly, clade 5 has the original clade and mutants $1,2,3,4$ as ancestoral clades and coincides with neither of them, adding five to the denominator and zero to the numerator. The clade turnover is hence $W_l = 3/15$.

In this example we assumed that we know the order in which the mutations occurred, but in fact the clade turnover can be computed without knowledge of the phylogenetic tree - we refer to [14] for the tree-free algorithm.

#### S5.2 Algorithmic description

We consider a population of cells dividing at rate $b$ and dying at rate $d$. The population starts from a single cell and the dynamics lasts for a time period $T$. With probability $\mu$ a mutation occurs in one of the daughter cells at division. The rates of birth and death are the same for all cells, meaning that the population is under volume growth (S1, $\rho_c = \infty$).

Extinctions are not *directly* metric from a single snapshot. Clones and clades that go extinct only leave evidence that they existed through their mutant offspring. We calculated the probability for a clone to have its parental clone become extinct [14],

$$\text{clone turnover}: \quad W_o(d/b,\mu,bT) = \frac{\mu(d/b + (\frac{\mu}{2})^2)}{(1-\frac{\mu}{2})^2(1-d/b-\frac{\mu}{2})}\frac{1-e^{-2(1-d/b-\frac{\mu}{2})bT}}{1-e^{-2\frac{\mu}{2}bT}}\ , \tag{S.4}$$

the probability for a clade to replace an ancestral clade,

$$\text{clade turnover}: \quad W_a(d/b, N) = \frac{d/b}{2\log N}(1 - \log(N)^{-2}) \;, \tag{S.5}$$

see Section S5.4 for a brief derivation. The clone turnover $W_o$ depends on the mutation rate, as mutations lead to the establishment of new clones. The clade turnover, however does not depend on the mutation rate. From the empirically measured turnover of clones and clades it is thus possible to infer the mutation rate and the rate of cell death (relative to the growth rate). The model assumes exponential growth and neutral mutations in a infinite site scenario. We infer clones by a simple clustering of coinciding mutations (see Section S8) as was done in Section S2. Again we find no significant changes when using LICHeE [9] instead.

The computation proceeds as follows:

1. estimate the population size as $N = n^{3/2}\frac{2}{\pi^{1/2}3^{1/4}}\frac{1}{2}$ given the number of samples in the plane $n$
2. clade turnover:
   (a) remove mutations that occur in less than 2 samples from clones
   (b) compute the fraction $\hat{W}_a$ of pairs where offspring clade and ancestor clade coincide
   (c) solve $\hat{W}_a = W_a(d/b, N(1 - d/b))$ for $d/b$
3. clone turnover:
   (a) remove clones that have mutations that occur in less than 2 samples
   (b) compute the fraction $\hat{W}_o$ of clones with missing parental clone
   (c) solve $\hat{W}_o = W_o(d/b, \mu, \log(N(1 - d/b))/(1 - d/b))$ for $\mu$
4. optional: subsample a fraction $0 < L < 1$ of mutations and repeat from (4) to infer $d/b, \mu$ and compare the results for different values of $L$

**Step 1** Both clade and clone turnover depend on the effective population size $N$, which depends on the resolution frequency as well as relative cell death rate $d/b$ through (S.28), as discussed in detail in sectionS7. To put the points from sectionS7 briefly: $N$ depends on the inverse frequency resolution $f_{\text{res}}$ which in turn relates to a set frequency cutoff within the sampled plane through eq. (S.27). The dependence of $N$ on $d/b$ is due to the stochastic extinction of clades from fluctuations in population size. Extinction is frequent at high rates $d/b$ and increases the expected population size of a surviving population by a factor $1/(1 - d/b)$. This $d/b$ dependence of $N$ is therefore included in the inference steps 2c and 3c.

**Steps 2a and 3a.** We set the frequency cutoff to $2/n$, where $n$ is the number of samples, by demanding that a mutation typically occurs in 2 or more of the $n$ genotyped samples. This cutoff is required to avoid cases where clones that are not sampled contribute erroneously to the turnover. The effective population size $N$ then is the size of the tumour at which, under neutral evolution, mutations occur which can just be recovered in the genotyped samples. Mutations that are found in only one sample thereby typically occurred later in time, after the population reached size $N$, and do not contribute to the observable measures of turnover. These mutations are removed from all clade genomes (2a) and their clones are removed from the set of clones. Plugging the frequency cutoff $2/n$ into eq. (S.27) gives the expression for $N$ in step 1.

**Steps 2b and 3b.** Here we calculate clade and clone turnover. See Section S5.1 for an example of how to determine clade- and clone-turnover from the set of surviving clones. We again refer to [14] for more detailed instructions.

**Steps 2c and 3c.** The clade and clone turnover calculated from the data are compared to the analytical expressions (S.5) and (S.4) to infer rates of mutation and cell death. First, $d/b$ is inferred from the clade turnover and then used in 3c to obtain the mutation rate $\mu$.

As an additional consistency check, the inferred rates can be used to compare a predicted cumulative SFS to the cumulative SFS obtained from the sequencing data, see also Section S4.2. Under neutral evolution and volume growth the cumulative SFS is linear with slope $\mu/(1-d/b)$ [15]. Fig. S21 shows a good match between the slope given by the inferred rates $d/b$,$\mu$ and the cumulative SFS of the WES whole-tumour frequencies of Ling *et al.*

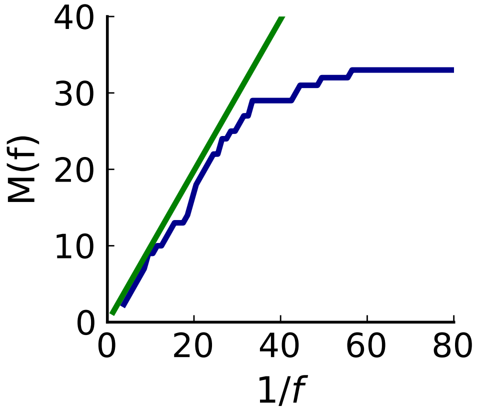

**Figure S21: Cumulative SFS of WES data compared to prediction using inferred rates $d/b$ and $\mu$.** The number of mutations with frequency larger than $f$ in the entire tumour against $1/f$ in genomically-resolved sequencing data (solid blue line). $M(f)$ flattens below the allele frequency corresponding to the sequencing resolution, $f_{\text{res}} = 1/40$. The green line indicates the expected curve under neutral evolution for the inferred rates of mutation $\mu$ and cell death $d$ [13] and provides a consistency check on our inference. Its slope is the scaled mutation rate $\mu/(1 - d/b)$.

In Section S7 we test how the choice of the population size $N$ affects the inference of mutation rate and the rate of cell death. We mimick the procedure in Ling *et al.*, where first 23 samples are taken for WE sequencing, and a small set of mutations is then probed by genotyping in a larger set of 285 samples (see Fig. SI S22). We find that 1) the number of samples $n$ required for estimating $N$ is given by the size of the large set of genotyped samples which determines the clonal composition, and 2) setting the lower cutoff of 2 supporting genotyped samples on mutations avoids biasing the turnover towards high death rates. In conclusion, the spatially-resolved set of the Ling *et al.* data reconstructs the clonal composition of the tumour, determines the effective population size $N$, and can be used to determine the rate of cell turnover $d/b$ and the mutation rate across the target genome. The preceding step of sampling at a lower spatial resolution for whole-exome sequencing, on the other hand, controls the size of the mutational target by filtering sites according to their coverage. We use the number of sites that pass these filters to calculate the mutation rate per nucleotide (see main text, "Cell turnover").

#### S5.3 Testing the inference based on clade/clone turnover

Our inference scheme for cell turnover based on stochastic clone extinction does not use spatial information. However, to test how the inference works under spatial sampling, we use data from spatial simulations under volume growth. Neutral evolution of the different clones is assumed, compatible with the original findings of Ling *et al.* [7] and the corresponding (cumulative) site frequency spectrum shown in Fig. S21.

We test the inference as specified in the steps above both in single cells sampled uniformly and using samples mimicking the sampling scheme in [7] as described in Section S1. Figure S22 compares inferred and underlying relative death rates and mutation rates in numerical simulations of volume growth ($\rho = \infty$).

In Fig. S23, we perform the same comparison, but focus on the regime of a high relative death rate. Specifically, we set the model parameters (relative death rate and mutation rate) close to the values inferred from the data of Ling *et al.* [7] and ask how well the inference scheme can recover those parameters. The average inferred rates match the underlying rates very well, but we also find large fluctuations in individual datasets. The uncertainty in the inferred mutation rate is connected to the high rate of cell death; a small change in $d/b$, say from 0.9 to 0.99 changes the number of generations needed for the cell population to grow by a factor of 10, and in turn changes the inferred mutation rate per generation by the same factor.

Employing LICHeE clustering of clones in simulations yields inferred rates $\mu$ and $d/b$ very similar to those obtained with our simple clustering scheme. As for the analysis of the direction of mutants in S2, we also compare our inference results for the simple clustering of the Ling *et al.* data to the results yielded by the

LICHeE clustering tool. LICHeE infers a split of clones that is almost identical to our clustering scheme, presumably due to the large number of samples and the low number of mutations. Consequently, turnover inference based on the set of clones by LICHeE results in a very similar, high turnover rate and the median mutation rate falls well into the error margin of inferred mutation rates for the simple clustering.

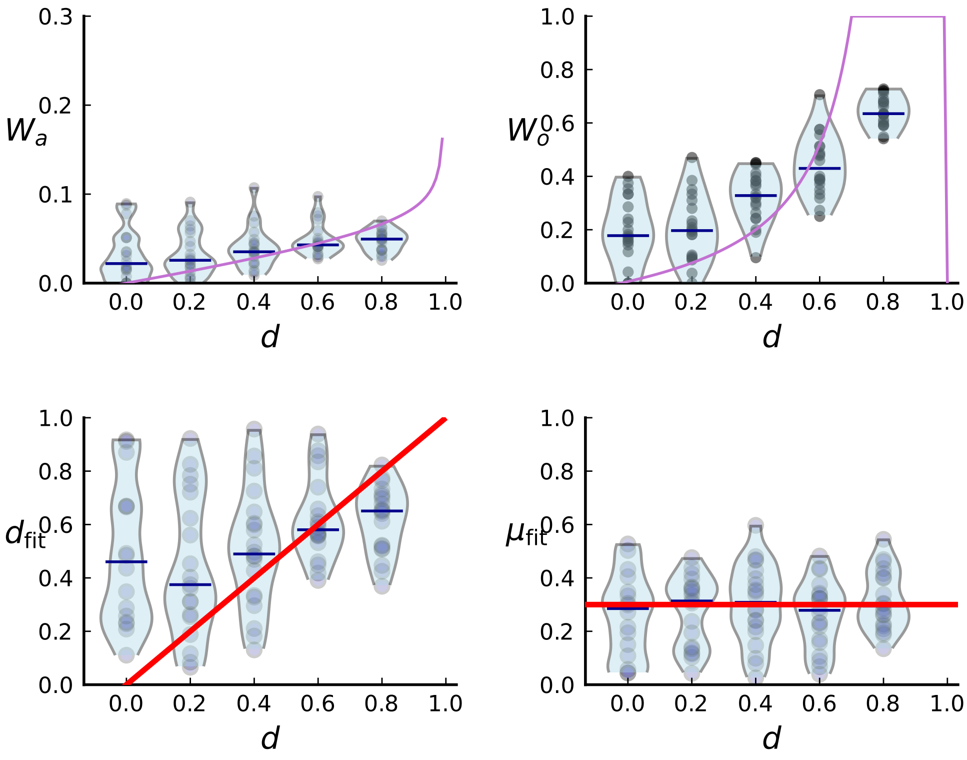

**Figure S22: Turnover inference for simulated sampling from cross sections in 3d over a range of death rates.** For each choice of death rate $d$ we simulate 20 populations under volume growth in 3d to 40000 cells each and take a cross-section. Mutations detected in a first set of 23 samples are then probed for in a set of 285 samples and used to infer $d$ and $\mu$.

We also apply the turnover-based inference to the tumours T1 and T2 analyzed by Li *et al.* and test on simulations with analogous spatial sampling schemes. The results are shown in Fig. S24. Clones of T1 and T2 were determined using the LICHeE tool for clone inference in multi-region sequencing data, see Section S8.

The turnover theory described in Section S5.4 assumes at most one mutation per cell division. The number of mutations validated by Li *et al.* is very large compared to Ling *et al.* (906 mutations for T1 and 565 mutations for T2), resulting in a genomic mutation rate that exceeds this limit. This situation can be easily dealt with by taking subsets of the validated mutations of relative size $L$, effectively shrinking the genome size by factor $L$. However, we find that the inferred mutation rates do not scale linear with $L$ - despite showing a downward trend with decreasing $L$ as required by the dependence of the clone turnover on the mutation rate. Only the inferred death rate $d$ is constant as expected.

To investigate, we use numerical simulations and again imitate the sampling of the two tumours in our 3d spatial simulations for different mutation rates, see figure S25. The inference scheme fails to infer the true rates in simulations when applying the sampling procedure of T1 and T2, whereas the planar sampling scheme of Ling *et al.* accurately infers $\mu$ (figure S22) and maintains the proper scaling in simulations when reducing the mutation rate by subsampling of mutations. We conclude that the samples in 3d data of Li *et al.* are not sufficiently dense to characterize the loss of clones with an accuracy sufficient for inference.

We suspect that the method of turnover-based inference breaks down in this case due to the low resolution

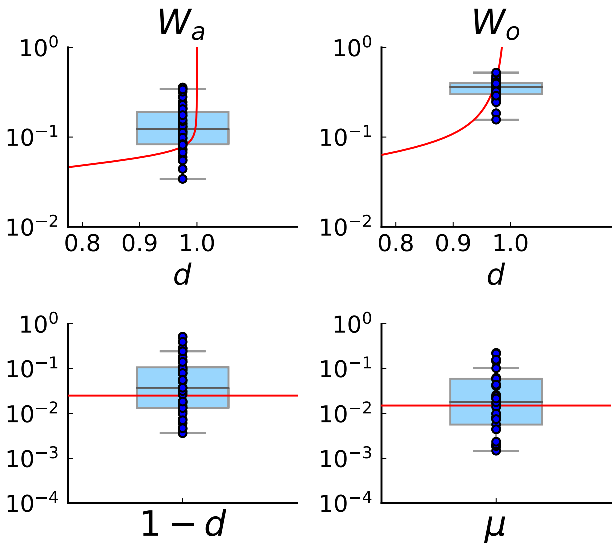

**Figure S23: Turnover inference for simulated sampling in 3d at the rates $d/b$ and $\mu$ compatible with those inferred from the Ling *et al.* data.** We run 40 simulations in 3d to 40000 cells at the rates inferred for the Ling *et al.* tumour, $d = 0.975$ and $\mu = 0.015$. Spatial sampling is performed as explained in Section S1.2 and Fig. S22. We plot the measured clade and clone turnover, $W_a$ (top left) and $W_o$ (top right), with the theoretically expected value as a function of d in red, and the inferred rates, $d$ (bottom left) and $\mu$ (bottom right), with the true value of the respective rate in red. The median of inferred rates is $d = 0.949$ and $\mu = 0.024$, but there are substantial fluctuations around these median rates.

of the clonal composition, which was particularly noticed for T2 in the previous sections. For both tumours, samples are taken from several slices of the upper hemispheres. The sampling covers the hemispheres rather uniformly, therefore, the estimate of the effective number of samples in the whole sphere $\tilde{n}$ is simply $2n$ for $n$ samples in the hemisphere. Using the same minimal frequency $f_{\min} = 2/n$ as before, the effective population size in step 1) of the turnover computation becomes $N = 2n\frac{1}{2} \approx 160$ for both tumours. This shows that taking a number of samples from the sphere that is comparable to that in the cross section implies a much lower sampling density. The low density of samples, however, in turn makes it harder to accurately infer clones their ancestry needed for inference.

An issue that goes beyond the scope of this work is the turnover inference if one or more lineages are under selection. Under selection the turnover theory no longer applies to the full population but needs to be modified to apply to individual lineages with different birth and death rates. It would be straightforward to include the effects of selection by defining such clade-specific turnover metrics, but this is outside the scope of this paper. We note that both T1 and T2 have a SFS that deviates to some degree from neutrality exhibiting potential clusters of mutations, which may be due to selection [8].

#### S5.4 Brief derivation of clade and clone turnover

The probability of eventual extinction can be obtained by first-step analysis, see [14] for details. Consider a single cell whose descendants form a clone or clade. The dynamics of clones and clades can be described by the same master equation by defining the effective rates of cell birth and death $\alpha$ and $\beta$. Clades gain cells at rate $\alpha = b$ and lose cells at rate $\beta = d$. But clones only gain a cell if neither parent nor offspring mutates at division, $\alpha = b \cdot (1 - (\mu/2)^2)$, and additionally lose a cell if both mutate $\beta = d + b \cdot (\mu/2)^2$. The probability that the originating cell and all offspring eventually die is denoted by $q$. After a single small time-step $\Delta t$, the cell either died with probability $p_0 = \beta\Delta t$, neither died nor divided $p_1 = 1 - (\alpha + \beta)\Delta t$ or divided $p_2 = \alpha\Delta t$. In each case the remaining cells each independently produce lineages which eventually go

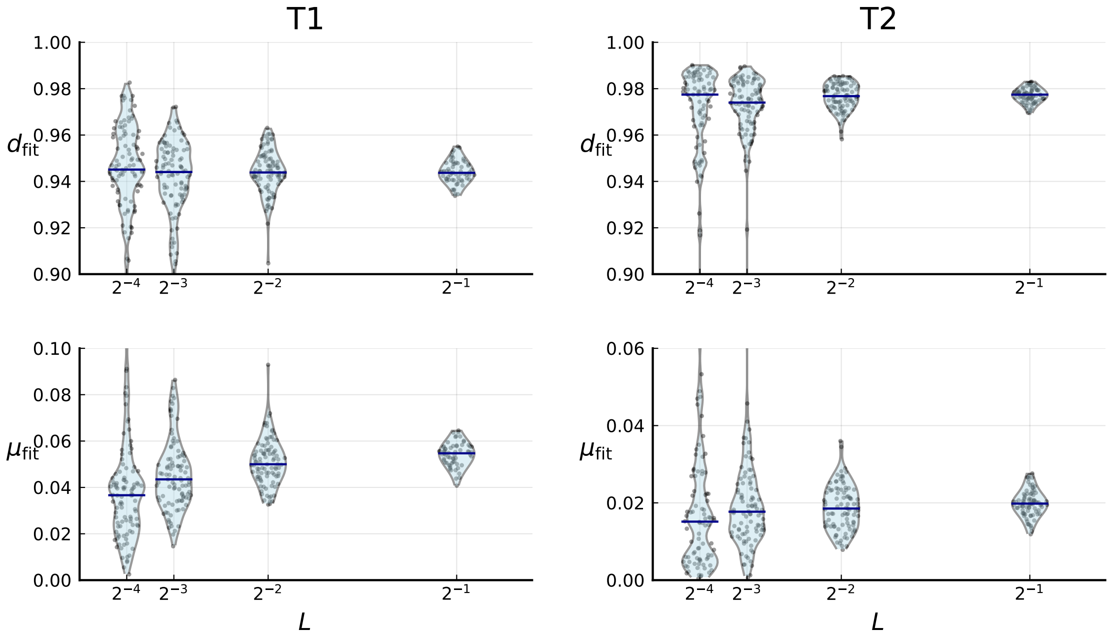

**Figure S24: Cell death and mutation rates inferred from the clade and clone turnover measured on the clones of T1 (left) and T2 (right).** For a given $L$ we repeatedly subsample a fraction $L$ of the mutations, calculate clade and clone turnover and infer $d$ and $\mu$ for each subsampled set of clones (violinplots, individual samples are shown as blue dots). Blue bars indicate the median. Clones of T1 and T2 were determined using the LICHeE tool for clone inference in multi-region sequencing data, see S8.

extinct with probability $q$. This leaves us with the following self-consistent equation for $q$

$$q = p_0 + p_1 q + p_2 q^2 \tag{S.6}$$

which can also be rewritten as $q = \frac{\beta}{\alpha+\beta} + \frac{\alpha}{\alpha+\beta}q^2$ in accordance with a time discrete branching process with probability of cell birth $\frac{\alpha}{\alpha+\beta}$ and death $\frac{\beta}{\alpha+\beta}$ ([1], chapter 3).
This equation is readily solved by $q = \min(1, \frac{\beta}{\alpha})$, which is

$$\text{for clades} \quad q = \frac{d}{b}, \quad \text{and for clones} \quad q = \frac{d + b(\frac{\mu}{2})^2}{b(1-\frac{\mu}{2})^2}. \tag{S.7}$$

Given the probability $p_n(t)$ of a clone/clade having size $n$ at time $t$, the probability that this clone/clade will eventually die out is

$$\sum_{n=0}^{\infty} q^n p_n(t) = z(q,t) \ , \tag{S.8}$$

which is also the definition of the generating function $z$ of the time-dependent probability mass function $p_n(t)$ at $q$. Instead of finding a solution for $p_n(t)$, we can obtain moments of the size distribution by taking derivatives of the generating function.

We begin by formulating the master equation governing the size of exponentially growing clones and clades. Consider a population that starts from a single cell and grows for a time $T$. Clones appear at different times and have different running times during which they can produce offspring. The probability that a cell was born at time $t$ is proportional to the population size $N(t) = e^{\lambda t}$. The normalising factor is $(e^{\lambda T} - 1)/\lambda$, yielding for large final times $T$ an exponential distribution $\lambda e^{\lambda(t-T)}$ for the running times $\tau \equiv T - t$.

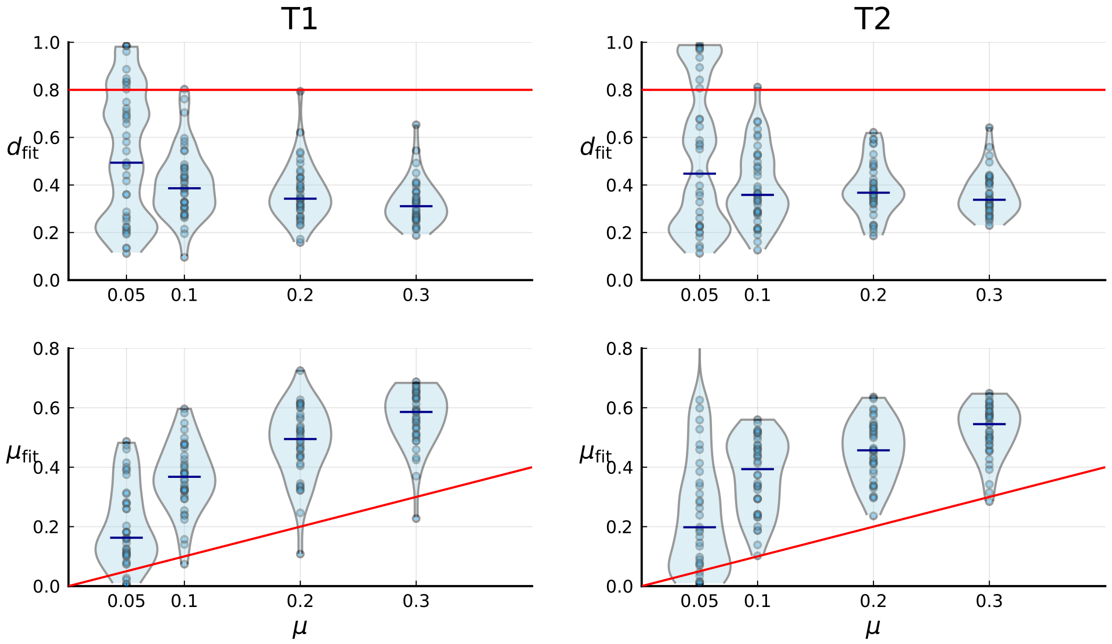

**Figure S25: Turnover-based inference of cell death and mutation rates under 3d sampling of spatial simulations.** 3d simulations are artificially sampled in a way that mimics spatial sampling, sequencing and genotyping in the tumours T1 (left) and T2 (right) by Li *et al.*, as in figure S7, explained in Section S2.2. Red lines indicate the true simulation rates. Under this sampling scheme, the inference of mutation and death rate fails, see text.

$$\partial_t p_n(t) = \alpha(n-1)p_{n-1}(t) - (\alpha+\beta)np_n(t) + \beta(n+1)p_{n+1}(t) - \lambda p_n(t)\ . \tag{S.9}$$

In addition to the first terms describing a standard birth-death process, the master equation also contains a term $-\lambda p_n(t)$ describing the exponential decay of available running time. The clone's founding cell is here born at time $t = 0$. Using the definition S.8 we get a linear partial differential equation for the generating function

$$\partial_t z(q,t) = (\alpha q^2 - (\alpha+\beta)q + \beta)\partial_q z(q,t) - \lambda z(q,t) \tag{S.10}$$

which for the boundary condition $z(q, 0) = q$ is solved by

$$z(q,t) = e^{-\lambda t}\frac{(1-q)\beta - e^{-(\alpha-\beta)t}(\beta - \alpha q)}{(1-q)\alpha - e^{-(\alpha-\beta)t}(\beta-\alpha q)}\ . \tag{S.11}$$

We defined the clone turnover as the probability for a clone to have its parental clone become extinct, and the clade turnover as the probability for a clade to replace an ancestral clade. In both cases, the offspring appears on a background of size $n$ with a probability proportional to $np_n(t)$, while the background becomes extinct with probability $q^n$. Averaging over both the time since the occurrence of the parental clone $t$ and parent size at birth $n$ defines the turnover parameter

$$W = \frac{\int_0^T dt \sum_{n=0}^{\infty} np_n(t)(\alpha/\beta)^n}{\int_0^T dt \sum_{n=0}^{\infty} np_n(t)} = \frac{q\int_0^T dt \partial_q|_{q=\beta/\alpha} z(q,t)}{\int_0^T dt \partial_q|_{q=1} z(q,t)} \tag{S.12}$$

$$= \frac{\beta}{\alpha}\frac{\lambda - (\alpha-\beta)}{\lambda + (\alpha-\beta)} \cdot \frac{1 - e^{-(\alpha-\beta+\lambda)T}}{1 - e^{(\alpha-\beta-\lambda)T}}\ . \tag{S.13}$$

Inserting the respective effective rates $\alpha$, $\beta$ for clones and clades finally yields the clone turnover (S.4) and clade turnover (S.5). For the detailed derivation we refer to [14].

### S6 Inference from mutational distances

In this section we investigate how to jointly infer the mutation rate per generation and the rate of cell death (relative to that of cell birth) from samples from a growing population using the mutational distances. Our approach is based on combining exact results of Stadler [11] on the distribution of the times since the last common ancestor in a population growing under birth-death dynamics and by Cheek and Johnston [16] on the statistics of the number of cell divisions in a sampled lineage. We compare the resulting statistics of the pairwise mutational distances with the distribution of mutational distances used recently Werner and collaborators [17] to jointly infer the mutation rate and the relative rate of cell death. Our treatment is a modification of the approach of Werner et al., and below we discuss the need for this modification.

The inference of the mutation rate per generation and the rate at which individuals die (relative to the rate of birth) is a long-standing problem. In population genetics, the times of speciation along a phylogenetic tree and the number of mutations along the tree have been used to perform this inference, using approaches based either on coalescence theory or on birth-death models [18, 19, 20]. In these approaches, the accuracy of the inferred parameters (mutations per generations, rates of a birth/death model) depend on how well the times of speciation can be reconstructed. An overview and unified perspective can be found in [21]. In the present case, due to the low number of mutations along phylogenetic branches the speciation times cannot be determined accurately.

#### S6.1 The distribution of mutational distances

Picking a pair of samples randomly and uniformly, the members of that pair differ by $m_1$ and $m_2$ mutations, respectively, from their last common ancestor. The mutations of the last common ancestor (relative to the normal tissue) are taken to be the intersection of the mutations in the two samples (this assumes no back-mutations). The distribution of the numbers of mutations $m_1$ and $m_2$ (mutational distances) across pairs of samples is the metric we will use for inference. A related approach has been taken by Ling *et al.* [7], who use both coalescent theory and birth-death models to calculate the fraction of pairs of samples that are genetically identical. The full spectrum of mutational distances has recently been used by Werner *et al.* [17] on multi-region tumour sequencing data.

The starting point is a population of cells (individuals) grown from a single cell. These cells independently divide at a rate $b$ and die at a rate $d$ (a birth-death process). Each cell carries a genome of length $L$ and during division, each genomic site of the two daughter cells is mutated with probability $\mu/L$. (In Werner *et al.* [17], $\mu$ denotes the mutation probability per site.) The population grows to a final size $N$, and the pairwise mutational distances between extant cells are measured. The aim is to infer the mutation rate and the rate of cell death relative to the rate of birth from the distribution of mutational distances.

Starting point is the probability distribution of the times of speciation under a birth death model. A result by Stadler (theorem 4.1 in [11]) gives the distribution of the time since the last common ancestor in pairs of extant cells picked uniformly from a population that has reached size $N$ as

$$\rho_{b,d,N}(t) = \frac{2be^{(b-d)t}}{N-1}\left(\frac{1}{\alpha+1}\right)^{N+1} \left[(\alpha+1)^{N+1} - \binom{N+1}{2}\alpha^2 - (N+1)\alpha - 1\right] , \tag{S.14}$$

where $\alpha = \frac{b-d}{b}\frac{e^{-(b-d)t}}{1-e^{-(b-d)t}}$. We rescale time to $\tau = bt$ obtaining the probability distribution for the rescaled time

$$\rho_{q,N}(\tau) = \frac{2e^{(1-q)\tau}}{N-1}\left(\frac{1}{\alpha+1}\right)^{N+1} \left[(\alpha+1)^{N+1} - \binom{N+1}{2}\alpha^2 - (N+1)\alpha - 1\right] , \tag{S.15}$$

with $q = d/b$ the death rate relative to the birth rate and $\alpha = (1-q)\frac{e^{-(1-q)\tau}}{1-e^{-(1-q)\tau}}$.

We now consider a pair of extant cells with time $\tau$ since their last common ancestor. The two cells and their ancestor define two lineages, with $i_1$ and $i_2$ cell divisions after speciation, respectively, since the last common ancestor. Cheek and Johnston [16] have calculated the distribution of the number of cell divisions of a lineage of age $\tau$ conditioned on survival. The distribution is affected by lineages that have undergone many cell divisions being statistically overrepresented in the set of extant cells. This is an example of the inspection paradox well known in probability [22], where for instance determining the average size of a family by polling individuals for the number of their siblings results in a biased estimate. The resulting statistics for the number of divisions $i$ found by Cheek and Johnston [16] is a mixture of Poisson distributions and in the present notation is

$$p_\tau(i) = 2^i e^{-d\tau} \frac{b - de^{-(b-d)\tau}}{b(e^{(b-d)\tau} - 1)} \sum_{j=0}^{i} \frac{1}{j!} \left( \frac{be^{(b-d)\tau} - d}{(b-d)e^{b\tau}} \left( \log \left( \frac{(b-d)e^{b\tau}}{be^{(b-d)\tau} - d} \right) \right)^j - e^{-b\tau}(b\tau)^j \right) \tag{S.16}$$

This expression follows equation (20) in the archived version (https://arxiv.org/pdf/2205.13875.pdf) of Cheek and Johnston; in the published version [16] the corresponding equation (21) differs by an erroneous factor of $(-1)^j$. The partial sum of the exponential series can be written in terms of the incomplete Gamma-function.

Finally, we look at the distribution of the number of mutations in a lineage given the number of cell divisions $i$ after speciation. Since mutations occur independently at different sites and at different divisions along a lineage, and the per-site mutation probability is small, the number $m$ of mutations after $i+1$ divisions (now including the speciation event itself) is Poisson distributed with mean $(i+1)\mu$, giving

$$P_{(i+1)\mu}(m) = \exp(-(i+1)\mu)((i+1)\mu)^m/m! \; . \tag{S.17}$$

We combine these results to estimate the distribution of the mutational distances $m_1$ and $m_2$ from the last common ancestor of a pair of cells randomly selected from extant cells in a population of size $N$, obtaining

$$P(m_1, m_2) = \int_0^\infty d\tau \rho_{q,N}(\tau) \sum_{i_1=0}^{\infty} p_\tau(i_1) P_{(i_1+1)\mu}(m_1) \sum_{i_2=0}^{\infty} p_\tau(i_2) P_{(i_2+1)\mu}(m_2) \; . \tag{S.18}$$

Similarly, the distribution of individual mutational distances (between the ancestor of a pair of cells and an element of that pair) is given by

$$P(m) = \int_0^\infty d\tau \rho_{q,N}(\tau) \sum_{i=0}^{\infty} p_\tau(i) P_{(i+1)\mu}(m) \tag{S.19}$$

These expressions neglect that choosing pairs of samples does not generate the typical number of cell divisions given the age of a lineage, but the error turns out to be small. Figure S26A shows the distribution of mutational distances for a single population ($\mu = 1, q = 3/4, N = 300$ compared to the result (S.19) (green line).

#### S6.2 Likelihood-based inference from pairwise mutational distances

Equation (S.18) serves as the likelihood function for the parameter inference. Enumerating all pairs of extant cells gives a list of mutational distances between each element of the pair and the last common ancestor, $\{m_1^\nu, m_2^\nu\}$, where the index $\nu$ runs over all pairs of extant cells. Maximizing the log-likelihood defined by (S.18)

$$\sum_\nu \log \left( P(m_1^\nu, m_2^\nu) \right) \tag{S.20}$$

with respect to the free parameters $q$ and $\mu$ for a given population size $N$ yields the maximum-likelihood estimates of the relative death rate $q = d/b$ and the genomic mutation rate $\mu$ per cell division and cell. The computation of double sums in the likelihood (S.18) can be avoided by precomputing the sum over $i_1$ for different values of $\tau$ and $m_1$ and using the same lookup-table for the sum over $i_2$.

The likelihood (S.18) differs in three points from the likelihood used in Werner *et al.* [17]: (i) We use a birth-death model, whereas Werner *et al.* use an approximation based on coalescence theory [18]. This point should be inconsequential, the correct use of either model should yield a correct inference (see below). (ii) We use the exact result by Cheek and Johnston [16] for the distribution of the number of divisions (S.16) which

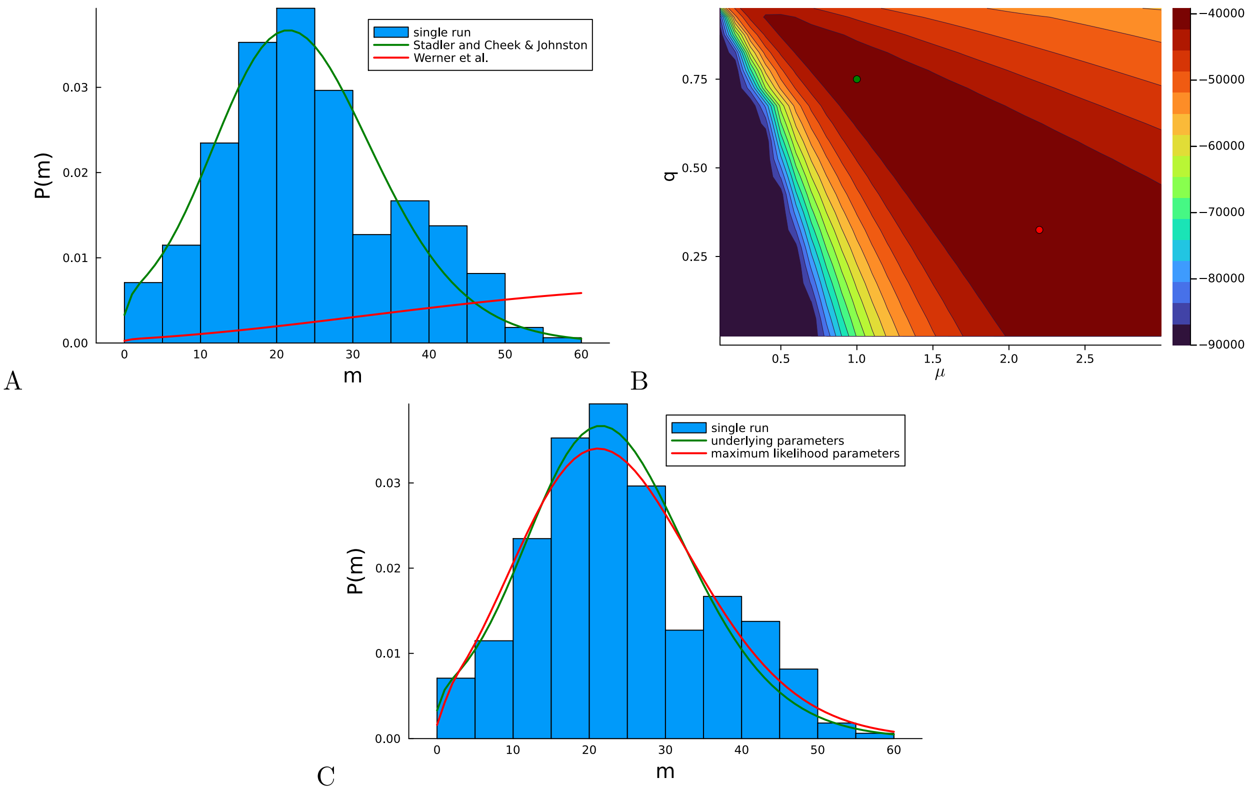

**Figure S26: Inference on artificial data using the likelihood** (S.18) **based on mutational distances.** For a concrete example, in a single run, a population is grown at rate of birth $b = 1$ and rate of death $d = 3/4$ with a genomic mutation rate $\mu = 1$ from a single cell until the population size has reached $N = 300$ cells. (A) The histogram of the mutational distances $m$ from the single population is shown in blue. The green solid line shows the corresponding distribution (S.19), the red solid line indicates the distribution of mutational distances derived in Werner *et al.* (equation (7) in [17]), see text. (B) The log-likelihood landscape defined by (S.18) with the mutation rate $\mu$ on the $x$-axis and the relative death rate $q = d/b$ on the $y$-axis. The underlying parameters $\mu = 1$ and $q = 0.75$ are indicated with a green dot, the maximum of the likelihood landscape is indicated by the red dot. The likelihood landscape shows a pronounced ridge of high likelihoods. The ridge-like shape of the likelihood landscape means that there are many different parameter values compatible with the data that yield nearly the same distribution of mutational distances (S.18). (C) The last point is illustrated by the distribution of the mutational distances: As in (A), the histogram of mutational distances $m$ is shown in blue, and the green solid line again shows the distribution (S.19) at the underlying parameters $\mu = 1$ and $q = 0.75$. The red line shows the the distribution (S.19) at the maximum-likelihood parameters (near $\mu = 2.2$ and $q = 0.33$).

accounts for the skewed statistics due to the inspection paradox. Werner et al. use a heuristic which does not account for this effect. (iii) The distribution (S.18) describes correlations between the mutational distance $m_1$ and $m_2$, which enter the log-likelihood (S.20). The method in [17] does not exploit these correlations.

Regarding point (i), neither the distribution of pairwise speciation times nor the coalescence probabilities per time can be evaluated without knowing the final population size $N$, see for instance [11] and [18] equation (5). Yet, the expression derived in [17] (equation (6) in [17]) for the coalescence probability does not depend on the final population size. The resulting distribution of mutational distances (equation (7) in [17]) is shown in Figure S26A (red line) and disagrees strongly both with our result (S.19) (green line) and numerical simulations.

Still, the likelihood of [17] can infer the model parameters correctly in the regime of large mutation rates, where there are clear oscillations in the distribution of mutational distances of period $\mu$: The number of cell

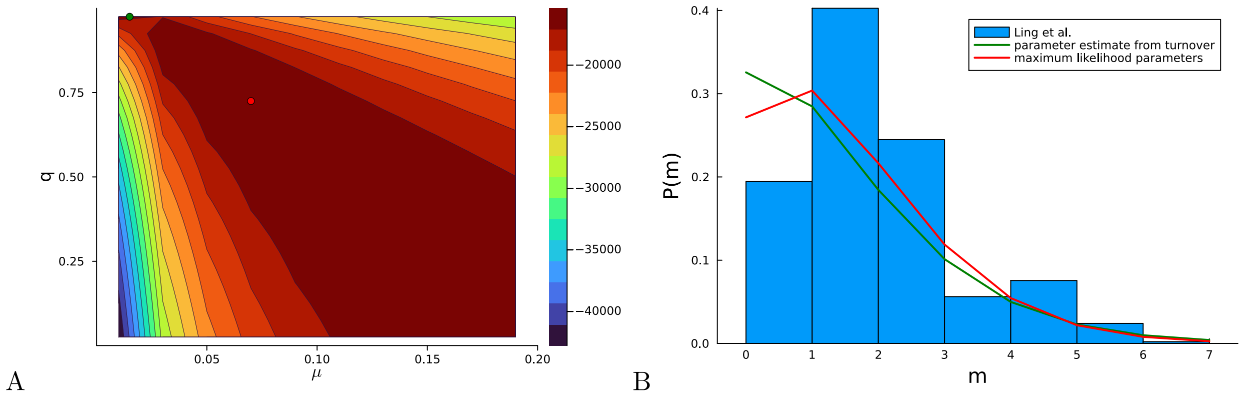

**Figure S27: Inference using the likelihood** (S.18) **on the the empirical data of Ling *et al.*** (A) The log-likelihood landscape defined by (S.18) with the mutation rate $\mu$ on the $x$-axis and the relative death rate on the $y$-axis, on the basis of mutational distances between samples in the data of [7]. The maximum of the likelihood landscape is indicated by the red dot ($\mu = 0.07$ and $q = 0.73$), the parameters previously inferred using the turnover of Section S5 ($q = 0.975, \mu = 0.015$) are indicated with a green dot (top edge left). As in the artificial data of in Fig. S26, the likelihood landscape shows a pronounced ridge of high likelihood. (B) The histogram of the mutational distances $m$ from samples of Ling *et al.* is shown in blue. The green line shows the distribution (S.19) at the parameters inferred using the turnover of Section S5. The red line shows the the distribution (S.19) at the maximum-likelihood parameters.

divisions between two cells is always integer, so the average number of mutations is an integer multiple of $\mu$. In the regime $\mu \gg 1$, one can resolve individual cell divisions in the spectrum of mutational distances, which cause distinguishable peaks at $\mu, 2\mu, 3\mu, \ldots$ (see artifical data in Fig. 2A in [17]). In this regime, the likelihood is dominated by parameters yielding the correct oscillation period. In the presence of such oscillations, the model parameters can easily be determined without using a likelihood by reading off the mutation rate from the oscillation period, and the relative death rate from a plot of the cumulative frequency of mutations versus the inverse frequency as in [15]. However, outside this regime, we found the method [17] returns incorrect parameter estimates. For whole-exome data, such as the Ling *et al.* data set and many of the data sets analyzed in [17], $\mu$ need not be much larger than one (in particular in WES data from tumours lacking the hypermutator phenotype), making a correct treatment of either coalescence probability or the distribution of speciation times necessary.

Figure S26B shows the log-likelihood (S.20) landscape for the same population of 300 cells used in Fig. S26A. A pronounced ridge-like shape is seen with a broad plateau of high likelihoods. This ridge includes the underlying parameters ($\mu = 1, q = 3/4$), although the maximum occurs elsewhere near $\mu = 2.2$ and $q = 0.3$. In general, broad maxima of the likelihood make the precise position of the likelihood maximum strongly depend on fluctuations in the data. Figure S26C shows how the distribution of mutational distances (S.19) at the maximum likelihood parameters (red line) and at the underlying parameters (green line). Both of these distributions fit the distribution of mutational distances quite well, although the fitting parameters are rather different. This is a property of the summary statistics used here, in line with the ridge of high likelihood values seen in Figs. S26B.

The same picture is also seen in the empirical data of Ling *et al.* [7]. Figure S27A shows a pronounced ridge of high likelihood. S27B shows the corresponding distribution of mutational distances (blue histogram) as well as the distribution (S.19) both for the parameters inferred using the genetic turnover (green line, see Section S5) and for the parameters maximising the likelihood (S.20) based on the mutational distances (red line). Again both parameter set lead to a reasonable fit with the distribution of mutational distances, indicating that mutational distances are not a suitable summary statistics for inference (unless the expected number of mutations per cell division is substantially larger than one, see above).

### S7 Estimating the population size $N$

The probability of coalescence or the time since the last common ancestor of two cells both depend on the population size. As a result, the distribution of the mutational distances (S.19) and hence the likelihood function (S.18) depend on the population size. Also, to infer the relative growth rate $q = d/b$ from the turnover parameter as in Section S5 we need to know the population size. Similarly, when telling surface growth from volume growth in Section S2, or tracking the spread of mutations in Section S4, we use mutations which entered the population at a specific time when the tumour had a certain size. Therefore our measures describe the growth dynamics at that particular time and tumour size, depending on the input data and mutation filters. In this section we discuss out the effective population sizes $N$ for different sets of mutations.

Due to the finite sequencing resolution this effective population size is not simply the final number of cells in the tumour population. Rare variants below the frequency resolution $f_{\mathrm{res}}$ are underrepresented or missing due to a finite sequencing coverage and stochastic errors of sequencing. Only mutations with sufficiently high frequency can be identified by sequencing, and those arose early in the evolution of the tumour when the tumour had a small size [15]. The statistics of mutations found in a large tumour with a finite sequencing depth corresponds to a smaller tumour where all mutations and frequencies are known. Here we address the question of what the effective population size of that smaller tumour is, given a particular frequency resolution $f_{\mathrm{res}}$ and test our answer in numerical simulations.

Under deterministic exponential growth - that is if one neglects fluctuations in clone sizes - the analysis is simple: each generation brings forth new mutations that have an initial frequency $f = 1/N$, where $N$ is the total population size at birth of the first mutant [15]. Under neutral evolution, new clones match the total population growth rate $\lambda = b - d$ (cell birth rate $b$ and death rate $d$) and thus maintain their frequency over time. Consequently, keeping only mutations down to a cutoff frequency $f_{\mathrm{res}}$ effectively casts the system into the early stages of tumour growth. Specifically, one observes the clonal makeup of the population at the time when mutations with frequency $f_{\mathrm{res}}$ have just appeared. Correspondingly, one would set the effective population size to

$$N = 1/f_{\mathrm{res}} \ , \tag{S.21}$$

see [15].

We now discuss a correction to this deterministic result due to stochastic fluctuations in the clone sizes. In an extreme case, such fluctuations can drive a clone to extinction. For $t \to \infty$ the scaled population size $e^{-\lambda t}N(t)$ (of a clone or an entire population) has been shown by Durrett ([1] chapter 3 theorem 1) to have a limiting distribution given by

$$\frac{d}{b}\delta_{N,0} + \frac{\lambda}{b}\left(1 - \frac{d}{b}\right) e^{-(1-\frac{d}{b})e^{-\lambda t}N} \ . \tag{S.22}$$

Here $\frac{d}{b}$ is the extinction probability, such that when conditioning on survival

$$(e^{-\lambda t}N(t)|N(t) > 0 \ \forall \ t) \overset{t\to\infty}{\longrightarrow} (1 - \frac{d}{b})e^{-(1-\frac{d}{b})e^{-\lambda t}N} \tag{S.23}$$

the expected population size becomes $\langle N(t)\rangle = \frac{1}{1-d/b}e^{\lambda t}$ for sufficiently large times $t$. The exponential term in this expression is the result due to determininistic growth, the prefactor is a correction due to fluctuations which can drive a clone to extinction. The contribution is large if the death rate $d$ is close to the rate of birth $b$, so extinctions events are frequent.

Whereas $e^{-\lambda t}N(t)$ is an exponentially distributed random variable for an ensemble of populations, but fixed for a given population, the mutant clones have an analogous relation between expectation values of the population size at birth of the mutant, $N$, and the mutation frequency $f_{\mathrm{res}}$

$$N = \frac{1}{1 - d/b}\frac{1}{f_{\mathrm{res}}} \ . \tag{S.24}$$

This gives a correction to the deterministic estimate $N = \frac{1}{f_{\mathrm{res}}}$ due to stochastic extinction events.

We test this relationship numerically in two ways. Figure S28 shows the population size at birth of a mutation against (one over) its final frequency and compares the numerical results to the deterministic estimate S.21 and the stochastic correction S.24.

Figure S29 probes how well the effective population size is described by (S.24) when low-frequency mutations up to a frequency threshold $f_{\mathrm{res}}$ are disregarded. We grow a population to size $N = 10000$ and disregard all mutations with final frequencies less than $f_{\mathrm{res}} = 0.01$. We then compare the resulting mutational distances (mean and standard deviation over each population) with those of populations grown to different final sizes *without* removing low-frequency mutants. Under the deterministic model (S.21), we expect the mutational distances to match those in a population of size $N = 100$. Instead, we find that the mutational distances match those in a population of a size given by the stochastic correction (S.24), which increases with the relative rate of cell death $q = d/b$.

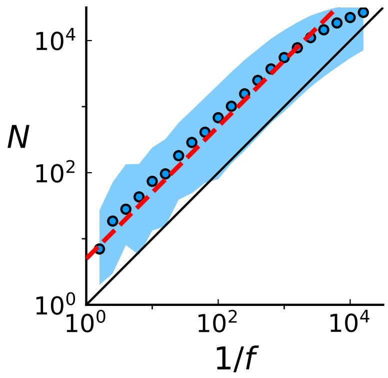

**Figure S28: Population size $N$ at birth of a mutation with final frequency $f$** Circular markers show the mean population size $N$ for a given frequency bin. The blue ribbon covers the 99$^{\mathrm{th}}$ percentile. The red line shows the relation (S.24) with the correction for fluctuating clone sizes, the black line shows the deterministic result (S.21). Mutations were collected from 20 3d spatial simulations up to a population size 40000 with birth rate $b = 1.$, death rate $d = 0.8$, and mutation rate $\mu = 0.3$.

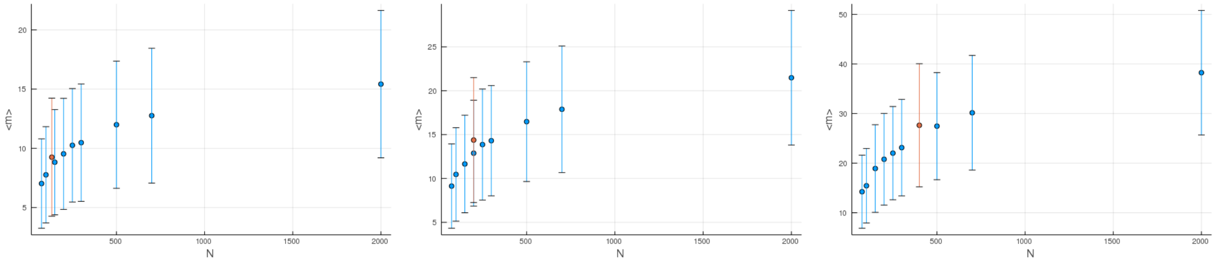

**Figure S29: The effect of disregarding low-frequency mutations.** The mean mutational distances and their standard deviations (blue markers and blue error bars, respectively) are shown for populations grown to different sizes $N$ ($x$ axis) at different relative death rates $q = 0.25, 0.5, 0.75$ (left to right). In each of these panels, the orange markers show the mean and standard deviation of the mutational distance for a population of size $N = 10000$ whose low-frequency mutants with frequency $f < 0.01$ have been discarded. For the effective population size we use (S.24). Rather than staying constant as they would under (S.21), the corresponding mean mutational distances increase with the relative death rate $q$, and this increase is compatible with (S.24), as seen from the orange markers following the blue ones. All quantities are averaged over 100 runs. Note that error bars give the standard deviations of the mutational distances, not standard errors.

#### S7.1 Frequency thresholds in the empirical data

It remains to set the minimal frequency $f_{\mathrm{res}}$ in the data of Ling *et al.* [7] and Li *et al.* [8]. $f_{\mathrm{res}}$ depends not only on the sequencing depth per sample but also the spatial sampling density. We discuss the dependency of the frequency threshold $f_{\text{res}}$ on the spatial sampling for the case of sampling within one cross section of a 3d spherical tumour, as in the Ling *et al.* data, or several slices of a hemisphere, as in the two tumours from Li *et al.*. In both cases, the idea is that only a fraction of the whole tumour is being sampled. Therefore, the rarest mutations found in the sample actually occur at a lower frequency within the whole 3d tumour and happened to fall within the sampled fraction of the tumour. We set a threshold on the frequency of mutations $f_m$ within the set of sequenced samples that is larger or equal to the frequency resolution for the given sequencing depth.

The minimal resolved whole tumour frequency $f_{\text{res}}$ is the smallest frequency with respect to the whole tumour of any mutation that occurs at a frequency greater than $f_m$ in the sampled plane. It necessarily describes a mutation that only occurs in our set of samples and has frequency $f_m$ therein. From our choice of $f_{\text{res}}$ on, we proceed to estimate the effective population size $N$ for each of the three tumours.

**Sampling from a tumour cross section (Ling *et al.*)**

Ling *et al.* [7] take samples from a cross section of the 3d tumour and detect mutations with a range of mutation frequencies within this set of samples. To find which is the minimal mutation frequency with respect to the whole tumour that still passes a threshold $f_m$ on the sampled slice, we ask how many samples under the given regular spacing would fit into a spherical tumour of the same diameter.

Given the spatially even and dense sampling we assume a triangular lattice and take the mean nearest neighbour distance between samples as the lattice constant $r$. We compute the mean spherical tumour radius $R$ using the dense packing ratio $\frac{\pi}{2\sqrt{3}}$ for the plane.

$$R = \sqrt{nr^2 / \frac{\pi}{2\sqrt{3}}} \tag{S.25}$$

Then $\tilde{n}$ is the number of samples that fill the spherical tumour of radius $R$ given the dense packing ratio $\frac{\pi}{3\sqrt{2}}$ for the 3d lattice:

$$\tilde{n} = (R/r)^3 \frac{\pi}{3\sqrt{2}} \tag{S.26}$$

$$= n^{3/2} \frac{2}{\sqrt{\pi\sqrt{3}}} \tag{S.27}$$

Finally, given a frequency threshold $f_m$ within the sampled plane for $n$ samples, we can estimate the minimal, resolved frequency in the whole tumour to be $f_{\text{res}} = f_m n/\tilde{n}$ because the total read-depth at a given site on the genome scales linearly with the number of samples from $n$ to $\tilde{n}$.

Essentially, if our lower bound on mutation frequency in the sampled plane is $f_m$ we include mutations down to the frequency $f_{\text{res}}$ with respect to the 3d tumour regardless of fluctuations in read counts. Increasing the sample size $n$ necessarily improves the frequency resolution as $f_m^{-1} \propto n$ under uniform sampling but the minimal, resolved frequency when sampling only the cross-section of a spherical tumour scales as $f_{\text{res}} \propto n^{3/2}$. Thus, the effective population size (S.24) becomes

$$N_{\text{eff}} = \frac{1}{\frac{n}{\tilde{n}} f_m} \frac{1}{1 - d/b} \tag{S.28}$$

$$= n^{3/2} \frac{2}{\sqrt{\pi\sqrt{3}}} \frac{1}{n f_m} \frac{1}{1 - d/b}. \tag{S.29}$$

In the rest of this section as well as in other sections we will simply refer to the effective population size $N_{\text{eff}}$ as $N$.

Based on this estimate and the correction for high cell turnover from equation (S.24), we use the effective population size (S.28) both for the inference of the relative death rate from the turnover, and for the analysis based on the mutational distances.

Because the frequency resolution and hence the population size estimate depends on the density of sampling we actually have two different effective population sizes: In the set of samples characterized by genotyping (high resolution, used to distinguish between volume and surface growth and for turnover inference) we consider mutations that occur in at least two of these samples. Such mutations can be assigned to clones, whereas it is impossible to confidently tell the relation of a mutation that occurs in a single sample to other mutations in its sample, see Section S8. Choosing the frequency cutoff such that mutations occur in at least two genotyped samples means that both metrics (directed growth and turnover) describe the dynamics of the set of clones up to the tumour size where a clone fails to colonize a spatially separated sample and reach detectable size therein until surgical resection. This is where the sampling density comes into play, since increasing the number of samples makes it more likely to sample a region where the clone has already reached sufficient size. We obtain the corresponding effective population size $N$ using equation (S.28): $n = 285$ samples underwent genotyping, giving an estimate of $\tilde{n} = 4000$ in 3d, the resolution is set to $n \cdot f_m = 2$ as specified in S5 because mutations must be recovered in two samples, and finally our turnover-based inference results in a turnover rate $d/b = 0.975$, giving $N = \tilde{n} \cdot \frac{1}{nf_m} \frac{1}{1-d/b} \approx 82500$.

In contrast, we can only measure the spatial dispersion of mutations that grew to detectable frequency in at least two whole-exome sequenced samples (deeply sequence samples and used to obtain cancer cell fractions and the site frequency spectrum). These samples resolve mutations up to a minimum frequency of $f_{\mathrm{res}} = 1/40$ (see S4), meaning that those mutations arose when the size of the tumour was only around $N \approx 3300$ in size, where $N$ is again the effective population size (S.28). Further details beyond this early stage of tumour development remain unresolved. Future data with a deeper sequencing and a higher spatial resolution will allow probing tumour evolution beyond this early stage.

The larger set of 285 genotyped samples targets the sites of mutations detected by whole-exome sequencing in the smaller set consisting of 23 samples. One might therefore ask whether the estimate on effective population size $N$ depends on the smaller, WES-sequenced set of samples which defines the selection of mutations or the larger genotyped set from which clones are inferred, see also Section S5. We explored this question in simulations over a range of turnover rates $d/b$ from 0 to 0.8 by applying a sampling procedure similar to that of Ling *et al.* (see S22), varying the size of the smaller (sequencing) and the larger (genotyping) set of samples, and observing how the measured clade and clone turnover $W_a$ and $W_o$ compares to the theoretical prediction which depends on $N$. How closely the measured turnover fits the theory at a given $N$ reflects in the quality of inference of $d/b$ and $\mu$. We find that, 1) the measured turnover fits the theory closest if $N$ is computed from the (larger) sample size of the genotyping set, 2) the measured turnover does not vary up to sampling noise when taking differently sized mutation-calling sets, and 3) the measured turnover decreases and increases for larger and smaller sample size of the genotyping set, respectively, as is theoretically expected for larger and smaller $N$, respectively.

**Sampling from a tumour hemisphere (Li *et al.*)**

Similarly, we estimate the effective population size for the tumours T1 and T2 from Li *et al.* [8]. In contrast to Ling *et al.* [7], samples are taken from several slices of the upper hemispheres of both tumours. Therefore, the estimate of the effective number of samples in the whole sphere $\tilde{n}$ is simply $2n$ for $n$ samples in the hemisphere.

Including the $d$-dependent correction in the estimate of effective population size $N$ for the conditioning on survival of tumours (S.24), given the estimated turnover rates $d/b = 0.944$ and 0.976, yields $N_{\mathrm{T1}} = 3000$ and $N_{\mathrm{T2}} = 6600$. Despite the high uncertainty in $d/b$, the estimated population size is lower than what we found for the clonal data of Ling *et al.* by more than an order of magnitude due to the lower density of samples in the data by Li *et al.*

### S8 Inference of clones

Our methods are based on the spatial distribution of genetic mutations in a mixed population of cells with different genotypes. It is straightforward to evaluate and interpret them for single-cell data where each sample represent a true genotype from the population. However, samples obtained by needle biopsies contain thousands of cells (about 20000 in Ling *et al.* and Li *et al.* or $\sim 3200$ in the Li *et al.* WGS samples) and generally contain a mixture of clones. This mixing of clones is particularly prevalent under volume growth, as can be seen by the dispersion of mutations, see S4. Genomic data from a large number of samples is well suited for the reconstruction of clones because the different samples already reveal to some extent the variety in genotypes. The clonal composition can be further resolved by clustering mutations by their sample frequencies on the level of single samples as well as the whole tumour (if mutation frequencies are available). For instance, two mutations might have very similar whole-tumour frequencies but do not coincide across samples (i.e. some samples have mutation A but not B) or have different frequencies in single samples and therefore do not belong to the same clone.

Clustering by frequency works best given a large number of mutations. Mutations belonging to one clone occur at equal frequency up to sampling noise and hence form a frequency cluster. The more mutations a clone has, the more confidently a clone can be identified, opposed to the random similarity in frequency between unrelated mutations. A larger number of samples on the other hand imposes stricter constraints on the clustering of mutations. Mutations must have very similar frequency in all samples to be part of the same clone, which might be hard to fulfill in the presence of sequencing noise.

We use two methods for the reconstruction of clones: As a first approach we use a simple clustering scheme where mutations that coincide across samples are clustered into clones. A simple example to illustrate this scheme mixes genotypes labeled by letters A, B, C, D into samples. The samples {AC},{AB},{BD} decompose into {[AC]}, {[A][B]}, {[BD]}, and as a rule, an ancestral clone will not be added to the sample, so {[AC]} does not become {[A],[AC]}. Despite its simplicity this scheme, when applied to artificially sampled simulations, is sufficient to recover the distributions of the direction angle (S4 and S3) and the rates from the turnover based inference already measured using simulated single cell data of the same simulations (see Fig. S22).

We further compare this simple clone reconstruction scheme to the LICHeE tool [9] designed specifically for multi-sample clone inference. LICHeE can leverage mutation frequencies of multiple samples to simultaneously cluster clones and determine their lineages and has been shown by the authors to better resolve tumour phylogenies when compared to tools designed for single bulk-samples [9].

In the case of the data by Ling *et al.*, LICHeE detects 27 clones and their phylogenetic tree from the presence profiles in the large set of 285 genotyped samples. Out of these clones, 22 agree with clones from our simple clustering, with the remaining 5 only differing each by one mutation which are ambiguous in their assignment to a clone and were removed from the set of mutations. However, both clustered sets of clones produce very similar results in the discussion of directed growth, see Section S2, and in the turnover based inference of mutation and cell death rates, see Section S5.

Li *et al.*, on the other hand, probe for the presence of 906 and 565 SNVs using genotyping in the tumours T1 and T2, respectively. Given the large number of mutations per sample, it is more likely than in the data by Ling *et al.* that the genotyping misses some mutations in a given sample or falsely reports them as present. Our simple clone reconstruction is sensitive to both false negatives and positives and very likely produces false clones here. The LICHeE tool, on the other hand, cannot resolve the constraint network defined by the large sets of genotyped sample profiles.

Instead, we apply LICHeE to the set of WGS samples, where it first clusters mutations by their measured variant allele frequencies before solving the (smaller) constraint network imposed by the WGS samples. LICHeE infers 25 and 14 clones for T1 and T2, respectively. We then proceed to recover the inferred clones in the genotyped samples by splitting the sample phenotypes into their constituent clones. This is useful for the analysis of the growth mode in S2.2, which relies on the higher spatial resolution of the genotyped samples. Testing this approach of splitting a sample into the different clones on the WGS samples from which LICHeE inferred the clones in the first place exactly recovers the split proposed by LICHeE.

### S9 Subsampling of mutations and mutational signatures

In order to estimate the statistical robustness of our estimate of the per-generation mutation rate and the relative rate of cell death, we perform the inference described in Section S5 also on subsets of the mutations. (A standard bootstrapping, where mutations are resampled without replacement is not feasible here, as it would produce duplicates of samples with zero mutational distance.) Figure S30 shows the inference results for the Ling et al. data for different relative sizes of the subset $L$ ranging from 0.5 to 0.9. No systematic drift with the sampling fraction $L$ is seen; correspondingly Figure 4C in the main text shows the results from the different fractions combined.

An unrelated point linked to subsampling of mutations concerns our observation of a particular mutational signature (SBS22) in the clonal mutations of Ling *et al.*, but not the subclonal mutations. This is compatible with exposure to the mutagen causing this particular signature during tumourigenesis, and the absence of the mutagen during the later stages of tumour growth. Here we ask how likely is an alternative scenario, namely that the signature is present also in subclonal mutations with the same statistical weight as seen in the clonal mutations, but is not detected due to the finite number of mutations (sampling noise).

| parent | offspring |
|---|---|
| [30] | [30, 33, 34] |
| root | [30] |
| root | [22] |
| root | [6] |
| [23, 24] | [23, 24, 25] |
| [10, 11] | [10, 11, 12, 13, 14] |
| [23, 24] | [23, 24, 26] |
| [6, 7] | [6, 7, 8] |
| root | [3] |
| root | [15, 16] |
| [15, 16] | [15, 16, 17] |
| [6] | [6, 7] |
| [6, 7] | [6, 7, 9] |
| [15, 16, 17] | [15, 16, 17, 18] |
| [3] | [3, 4] |
| [30] | [30, 31] |
| [23, 24, 27] | [23, 24, 27, 28] |
| root | [10] |
| [30] | [30, 35] |
| [23, 24] | [23, 24, 29] |
| [23, 24] | [23, 24, 27] |
| root | [23, 24] |
| [3] | [3, 5] |
| [30, 31] | [30, 31, 32] |
| root | [1, 2] |
| [10] | [10, 11] |
| [15, 16] | [15, 16, 19, 20, 21] |

**Table S1: Clones and their phylogeny from the Ling *et al.* data:** Each row shows one branch in the phylogenic tree inferred by LICHeE for the Ling et al. data. Columns labelled 'parent' and 'offspring' show mutations of the parental and offspring clone respectively. The mutations along the branch are the private mutations the offspring has gained, the difference between the two sets of mutations. 'root' indicates that the clone has no ancestor in the given set of clones other than the common ancestor to all clones which has the clonal mutations.

We decomposed the mutational profiles of the tumours from the two studies using our in-house method SigNet [23]. SigNet is based on an artificial neural network and predicts the weights of the COSMIC v3 mutational signature catalog [24] in individual samples. If the sample profile does not correspond to the training data, it is decomposed with non-negative least squares (NNLS). SBS1, SBS5 and SBS40 are associated with endogenous mutational processes. Since the samples that contain SBS25 were decomposed with a non-negative least squares approach (as they were classified as out-of-distribution by SigNet), it is likely that SBS25 is an erroneous assignment derived from SBS5 and SBS22 mutations [24]. The high number of signatures classified as "Others" in samples for which the NNLS approach was used is compatible with the low accuracy of this approach when the number of mutations in the sample is small.

To address the effect of subsampling on the mutational signatures, we run the in-house SigNet algorithm [23] on $10^4$ artificially generated instances with different weights of SBS22. Weights of the other signatures contributing to each simulated sample were maintained at the same relative proportions as found in the decomposition of the subclonal mutations in the whole-exome data of Ling et al. for all weights of SBS22. Each instance contained 66 mutations, the number of subclonal exomic mutations in the Ling *et al.* tumour.

The weight of SBS22 when pooling clonal and subclonal mutations was 0.78. The probability that although subclonal mutations have SBS22 at this particular weight, but the signature is not detected on the basis of

**T1**

| parent | offspring | mutations |
|---|---|---|
| 15 | 1 | 32 |
| 15 | 2 | 25 |
| 13 | 3 | 8 |
| 8 | 4 | 7 |
| root | 5 | 68 |
| 20 | 6 | 6 |
| 19 | 7 | 30 |
| 9 | 8 | 10 |
| 5 | 9 | 56 |
| 13 | 10 | 3 |
| 20 | 11 | 6 |
| 16 | 12 | 19 |
| 21 | 13 | 10 |
| 9 | 14 | 14 |
| 7 | 15 | 57 |
| 5 | 16 | 35 |
| 12 | 17 | 12 |
| 21 | 18 | 11 |
| root | 19 | 46 |
| 8 | 20 | 14 |
| 22 | 21 | 9 |
| 12 | 22 | 18 |
| root | 23 | 2 |
| root | 24 | 55 |
| 22 | 25 | 10 |

**T2**

| parent | offspring | mutations |
|---|---|---|
| 4 | 1 | 2 |
| 3 | 2 | 14 |
| 8 | 3 | 6 |
| root | 4 | 20 |
| 11 | 5 | 4 |
| 7 | 6 | 16 |
| 1 | 7 | 27 |
| 11 | 8 | 17 |
| 8 | 9 | 7 |
| root | 10 | 38 |
| 4 | 11 | 56 |
| 7 | 12 | 10 |
| 3 | 13 | 2 |
| root | 14 | 39 |

**Table S2: Li data lichee branches for tumours T1 and T2:** Each row shows one branch in the phylogenic tree inferred by LICHeE for the T1 tumour of the Li et al. data. Clones have too many private mutations to list and are therefore only counted (columns 'parent' and 'offspring' along with the number of mutations along a branch ('mutations'). 'root' indicates that the clone has no ancestor in the given set of clones other than the common ancestor to all clones which has the clonal mutations.

only 66 mutations is estimated to be less than $10^{-4}$ (the event did not occur in $10^4$ runs of the simulation). This provides an estimate of how likely the alternative scenario of undetected SBS22 is ($p$-value), leading us to reject the alternative scenario.

### References

[1] Richard Durrett. *Branching Process Models of Cancer.* Springer, 2015.

[2] Maartje van der Heijden, Daniël M Miedema, Bartlomiej Waclaw, Veronique L Veenstra, Maria C Lecca, Lisanne E Nijman, Erik van Dijk, Sanne M van Neerven, Sophie C Lodestijn, Kristiaan J Lenos, et al. Spatiotemporal regulation of clonogenicity in colorectal cancer xenografts. *Proceedings of the National Academy of Sciences*, 116(13):6140–6145, 2019.

[3] Ketevan Chkhaidze, Timon Heide, Benjamin Werner, Marc J Williams, Weini Huang, Giulio Caravagna, Trevor A Graham, and Andrea Sottoriva. Spatially constrained tumour growth affects the patterns of clonal selection and neutral drift in cancer genomic data. *PLoS Computational Biology*, 15(7):e1007243, 2019.

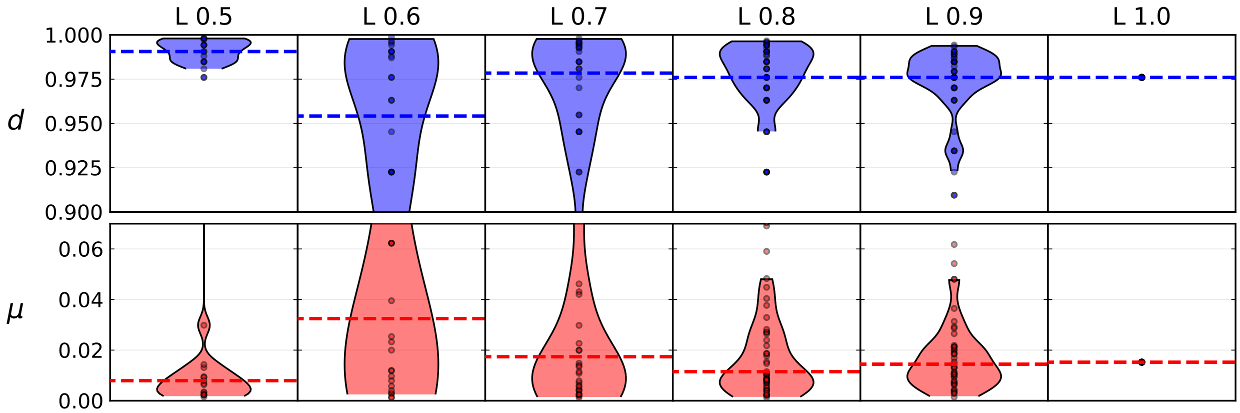

**Figure S30: Violin plots of model parameters (number of mutations per generation and relative death rate) inferred from different fractions $L$ of the mutations present in the Ling et al. data.**

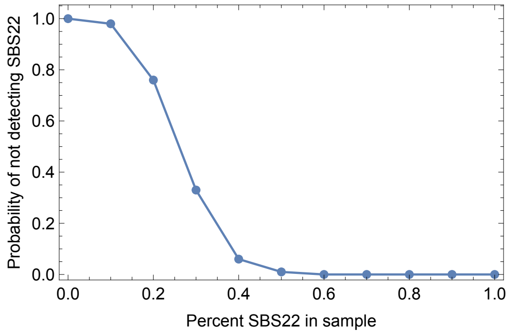

**Figure S31: Sampling mutations with different weights of a particular signature (SBS22).** Weights of SBS22 are shown on the $x$-axis, the probability of not detecting SBS22 given 66 mutations is shown on the $y$ axis and rapidly decays with the underlying weight of SBS22, see text.
